# Pathogenic variants in autophagy-tethering factor EPG5 drive neurodegeneration through mitochondrial dysfunction and innate immune activation

**DOI:** 10.1101/2025.05.23.655806

**Authors:** Kritarth Singh, Hormos Salimi Dafsari, Olivia Gillham, Haoyu Chi, Ivet Mandzhukova, Ioanna Kourouzidou, Preethi Sheshadri, Chih-Yao Chung, Valeria Pingitore, Fleur Vansenne, David L. Selwood, Diana Pendin, Gyorgy Szabadkai, Manolis Fanto, Heinz Jungbluth, Michael R. Duchen

**Author notes:** Correspondence to: Michael R. Duchen. equal contribution (Kritarth Singh, Hormos Salimi Dafsari). joint supervision (Manolis Fanto, Heinz Jungbluth, Michael R. Duchen).

## Abstract

The autophagy-tethering factor, ectopic P-granule 5 autophagy protein (EPG5), plays a key role in autophagosome-lysosome fusion. Impaired autophagy associated with pathogenic variants in *EPG5* cause a rare devastating multisystem disorder known as Vici syndrome, which includes neurodevelopmental defects, severe progressive neurodegeneration and immunodeficiency. The pathophysiological mechanisms driving disease presentation and progression are not understood. In patient-derived fibroblasts and iPS cells differentiated to cortical neurons, we found that impaired mitophagy leads to mitochondrial bioenergetic dysfunction. Physiological Ca^2+^ signals resulted in paradoxical mitochondrial Ca^2+^ overload attributed to downregulation of MICU1/3. Ca^2+^ signals caused mitochondrial depolarisation, mtDNA release and activation of the cGAS-STING pathway, reversed by pharmacological inhibition of the mitochondrial permeability transition pore (mPTP) or of the STING pathway. Thus, we have identified multiple potential therapeutic targets driving disease progression associated with pathogenic EPG5 mutations, including impaired mitochondrial bioenergetics, mitochondrial Ca^2+^ overload, vulnerability to mPTP opening and activation of innate immune signalling.

## Introduction

Autophagy is an evolutionarily conserved, fundamental intracellular homeostatic process with essential roles in metabolic adaptation, defence against infections and the quality control of defective proteins and organelles including mitochondria (Klionsky et al., 2021). Ectopic P-granules 5 autophagy tethering factor (EPG5), initially reported in *Caenorhabditis elegans*, functions as a tethering protein in concert with specific SNARE complexes to facilitate specific fusion events between autophagosomes and lysosomes and the formation of degradative autolysosomes (Tian et al., 2010; Guo et al., 2014; Wang et al., 2016; Hori et al., 2017). Recessive mutations in human EPG5 cause a spectrum of neurodevelopmental disorders termed as *EPG5*-related disorders (*EPG5*-RD) ranging from the severe early-onset multisystem disorder Vici syndrome to relatively milder neurodevelopmental manifestations with progressive neurodegeneration (Cullup et al., 2013; Byrne et al., 2016b; Dafsari et al., 2022). Vici syndrome patients are characterized by the key diagnostic features of microcephaly, callosal agenesis, cataracts, hypopigmentation, cardiomyopathy, (combined) immunodeficiency and failure to thrive, followed by severe progressive neurodegeneration with a median life expectancy of approximately 24 months (Byrne et al., 2016a; Deneubourg et al., 2022). While most bi-allelic loss-of-function EPG5 mutations are associated with Vici syndrome, pathogenic missense variants in *EPG5* can lead to less severe clinical presentations. An attenuated EPG5 function is predicted to underlie the phenotypic variability seen in *EPG5*-RD (Vansenne et al., 2022; Dafsari et al., 2024). However, it is unclear whether the residual function of pathogenic *EPG5* variants correlates directly to a corresponding defect in autophagy alone or if other downstream cellular processes are also disrupted. Interestingly, preliminary observations indicate an increased frequency of adult-onset neurodegenerative disorders in (putative) heterozygous *EPG5* variant carriers, suggesting a potential dosage effect.

A previous study has reported that the histopathological appearance of *EPG5*-related Vici syndrome, often including marked ultrastructural mitochondrial abnormalities and decreased respiratory chain enzyme activity, may mimic primary mitochondrial disorders (McClelland et al., 2010; Byrne et al., 2016b) and some patients have been suspected to have a mitochondrial cytopathy before the causative pathogenic *EPG5* variants were genetically confirmed (Balasubramaniam et al., 2018; Waldrop et al., 2018). Our recent findings have shown that defective mitophagy and associated mitochondrial dysfunction play a significant role in *EPG5*-related Vici syndrome and that some features of the disorder may be a direct consequence of mitochondrial dysfunction (Dafsari et al., 2024). Impaired mitochondrial energy metabolism has been recognized as a key factor in the pathogenesis of many common adult-onset neurodegenerative disorders including amyotrophic lateral sclerosis (ALS), Parkinson’s and Alzheimer’s disease (Duchen and Szabadkai, 2010; Schon and Przedborski, 2011; Area-Gomez et al., 2019; Chu, 2022; Suomalainen and Nunnari, 2024). Mitochondrial dysfunction resulting from impaired mitochondrial quality control can lead to impaired ATP homeostasis, oxidative stress, impaired Ca^2+^ buffering and altered mitochondrial Ca^2+^ signalling, all of which contribute to neuronal dysfunction and cell death characteristic of many neurodegenerative disorders (Duchen, 2012; Devine and Kittler, 2018; Boyman et al., 2020; Plotegher et al., 2020).

Mitochondria actively maintain neuronal Ca^2+^ homeostasis through Ca^2+^ uptake and efflux pathways during physiological cytosolic [Ca^2+^] ([Ca^2+^]_c_) signals (Rizzuto et al., 2012; Plotegher et al., 2021; Verma et al., 2022). Mitochondrial Ca^2+^ uptake is mediated by a Ca^2+^- selective ion channel, the mitochondrial calcium uniporter (MCU) located in the inner mitochondrial membrane (IMM) (Kamer and Mootha, 2015; Giorgi et al., 2018). This hetero-oligomeric channel is composed of the pore-forming protein, MCU (Baughman et al., 2011; De Stefani et al., 2011), a scaffold, essential MCU regulator (EMRE) (Sancak et al., 2013), and Ca^2+^-sensitive gatekeepers, MICU1, MICU2, and MICU3 (Perocchi et al., 2010; Plovanich et al., 2013; Patron et al., 2019). The dimer of MICU proteins in concert with EMRE exerts a tight control on mitochondrial Ca^2+^ entry during changes in local [Ca^2+^]_c_ (Csordas et al., 2013; Patron et al., 2014; Fan et al., 2020). A rise in mitochondrial matrix [Ca^2+^] ([Ca^2+^]_m_) increases the rate of TCA cycle-driven NADH generation and the rate of oxidative ATP production (Duchen, 1992; Jouaville et al., 1999; Denton, 2009). Accumulation of Ca^2+^ in mitochondria is balanced by Ca^2+^ efflux through the NLCX exchanger which normally maintains a low [Ca^2+^]_m_ (Palty et al., 2010; Garbincius and Elrod, 2022). Supraphysiological accumulation of Ca^2+^, however, can trigger the opening of a large conductance channel in the IMM, the mitochondrial permeability transition pore (mPTP), resulting in the collapse of ΔΨ_m_, ATP hydrolysis by the F_O_F_1_ ATPase and mitochondrial osmotic swelling (Brenner and Moulin, 2012; Bhosale et al., 2015; Briston et al., 2019). This mitochondrial catastrophe seems to be a common final path driving mitochondrial Ca^2+^ overload-induced excitotoxic injury and neuronal cell death as evident in *MICU1*-KO mice (Singh et al., 2022) and implicated in neurodegenerative diseases such as ALS (Lautenschlager et al., 2013), Alzheimer’s (Jadiya et al., 2021) and Parkinson’s disease (Choi et al., 2022) as well as in several muscular dystrophies and myopathies (Palma et al., 2009; Rao et al., 2014).

Mitochondrial Ca^2+^ overload may also trigger the release of mtDNA via selective or limited mPTP opening in a subset of damaged mitochondria, activating a cytoprotective innate inflammatory response and evade cell death similar to the phenomenon described recently during cellular senescence (Victorelli et al., 2023). Cytosolic mtDNA is sensed as genotoxic stress by the DNA-sensing cyclic GMP–AMP synthase (cGAS)–stimulator of interferon response cGAMP interactor 1 (STING) pathway that activates type I/III IFN and nuclear factor (NF)-κB signalling (West et al., 2015; Newman and Shadel, 2023). Recent findings indicate that a mtDNA-driven inflammatory response may trigger a more insidious, low-grade chronic neuronal loss, a feature of many late-onset neurodegenerative diseases, especially when coincident with impaired mitophagy (Sliter et al., 2018; Jauhari et al., 2020; Yu et al., 2020; Jimenez-Loygorri et al., 2024).

Mitochondrial Ca^2+^ overload culminating in the loss of mitochondrial function, Ca^2+^-dependent cell death, inflammation, and progressive cell loss has been implicated in major neurodegenerative diseases (Filadi and Pizzo, 2020). Impaired mitochondrial Ca^2+^ homeostasis may underlie progressive infantile epileptic encephalopathy found in nearly two-thirds of patients with *EPG5*-RD (Deneubourg et al., 2025) and an associated chronic inflammatory response may explain high levels of cytokines measured in Vici syndrome patients (Piano Mortari et al., 2018). However, evidence for a specific mechanism driving aberrant mitochondrial Ca^2+^ signalling leading to mitochondrial dysfunction and inflammatory response is scarce and poorly understood.

Our recent findings point towards mitochondrial dysfunction and the failure of autophagic removal of damaged mitochondria allowing progressive accumulation of mitochondrial defects as a major factor underlying the neurodegenerative disease and inflammatory dysregulation seen in patients with Vici syndrome and other *EPG5*-RDs (Dafsari et al., 2024). We now identify impaired mitochondrial Ca^2+^ homeostasis as the causative mechanism for mtDNA-driven STING-type I IFN inflammatory response in patient-derived fibroblasts with truncating and missense *EPG5* variants. iPSC-derived cortical neurons carrying a founder *EPG5* pathogenic variant showed increased sensitivity to excitotoxicity in response to low concentrations of glutamate that were innocuous in control cells, with responses characterised by mitochondrial Ca^2+^ overload, delayed calcium deregulation (DCD), loss of ΔΨ_m_ and neuronal cell death. We demonstrate that impaired mitochondrial bioenergetic function and Ca^2+^ overload found in patient-derived cells and *EPG5* mutant neurons were attributable to the downregulation of MICU1 and increased susceptibility to mPTP opening. Remarkably, pharmacological inhibition of mPTP opening attenuated inflammatory signals and rescued mitochondrial bioenergetic function in patient-derived cells and *EPG5* mutant neurons. These findings highlight a key pathophysiological mechanism in Vici syndrome and other *EPG5*-RD and signpost potential therapeutic strategies for the rapidly expanding clinical and genetic spectrum of patients with *EPG5*- and other autophagy-related disorders.

## Results

### Mitochondrial bioenergetic function is impaired in patient-derived fibroblasts bearing pathogenic EPG5 mutations

To characterise the impact of pathogenic EPG5 mutations on mitochondrial metabolism and bioenergetic function, we examined human dermal fibroblasts derived from patients (Supplementary Table S1) either carrying homozygous missense mutations (p.Gln336Arg, p.Arg1621Gln, hereafter referred to as patient 1 and patient 2 respectively) or compound heterozygous and homozygous truncating mutations (p.Arg299*/Pro1827Ala, p.Phe1604Glyfs*20, hereafter referred to as patient 3 and patient 4 respectively) as well as healthy control cell lines from donors matched for age and sex. Protein expression analysis by immunoblotting in control and patient fibroblasts revealed an almost complete absence of normal EPG5 protein in all patient cells (Fig. 1A-B), even in missense mutations, while in comparison the *EPG5* mRNA expression showed a significant reduction in patient cells (Fig. S1A).

**Figure 1:**
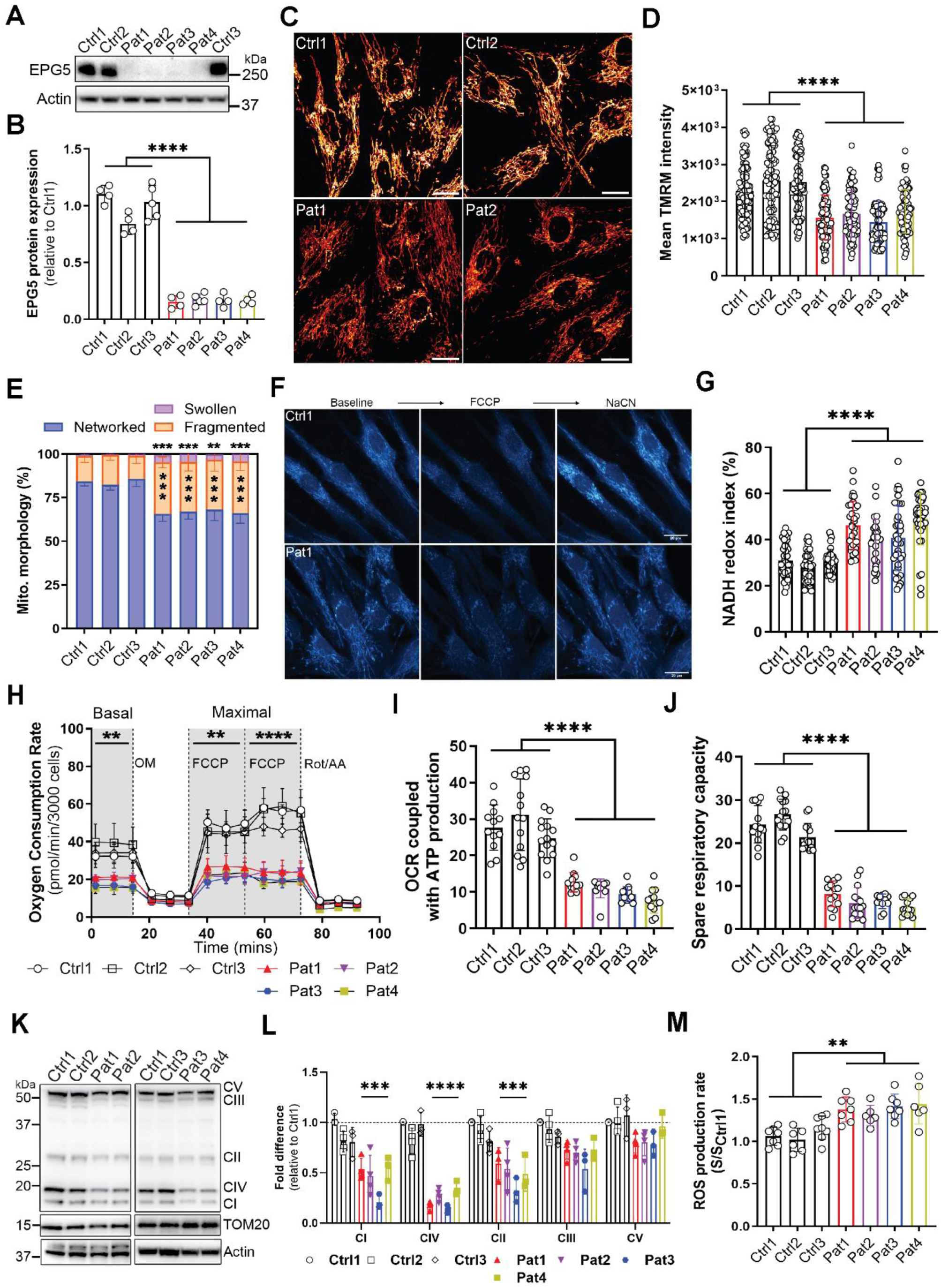
Patient-derived fibroblasts bearing pathogenic EPG5 mutations show impaired mitochondrial bioenergetic function and respiratory defect. **(A)** Immunoblot image of EPG5 protein expression in controls and patient-derived fibroblasts. Actin was used as a loading control. **(B)** Protein levels normalized to those in control 1, (n_exp_ = 4, \*\*\*\**p* <0.0001). **(C)** Representative confocal image of TMRM labelled cells showing ΔΨ_m_ in control 1, control 2, patient 1 and patient 2 fibroblasts. Scale bars: 20 μm. **(D)** Mean TMRM values showing steady state ΔΨ_m_, (n_exp_ = 4, n_cells_ analysed for each control and patient = 95-110, \*\*\*\**p* <0.0001). **(E)** Mitochondrial morphometric analysis of all TMRM confocal images classified into networked, fragmented and swollen mitochondria represented as percentage of total mitochondrial population, (n_exp_ = 4, n_cells_ analysed for each control and patient = 80-91, \*\**p=* 0.0034, \*\*\**p=* 0.0002). **(F)** Representative NADH confocal image of control 1 and patient 1 fibroblasts at baseline and following sequential application of the protonophore FCCP and cyanide (NaCN; complex IV inhibitor), Scale bars: 20 μm. **(G)** NADH redox index plotted as percentage of the minimum (with FCCP) and maximum (with NaCN) values, (n_exp_ = 3, n_cells_ analysed for each control and patient = 33-40, \*\*\*\**p* <0.0001). **(H)** Normalised OCR traces from controls and patient fibroblasts, (n_exp_ = 3, n_rep_ = 4, \*\**p=* 0.0043 and 0.0038, \*\*\*\**p* <0.0001). **(I-J)** Normalised ATP-linked respiration and spare reserve capacity of controls and patient fibroblasts, (n_exp_ = 3, n_rep_ = 4, \*\*\*\**p* <0.0001). **(K)** Immunoblot image of OXPHOS protein subunits expression from whole cell lysates of controls and patient-derived fibroblasts. TOM20 and Actin were used as loading controls. **(L)** Protein expression levels normalized and plotted as fold difference relative to control, (n_exp_ = 3-4, \*\*\**p=* 0.0003 and 0.0001, \*\*\*\**p* <0.0001). **(M)** DHE oxidation rates plotted as ROS production rate and normalized to control 1, (n_exp_ = 3, \*\**p=* 0.0015). Data (B, D, E, G, I, J, L and M) are expressed as mean ± SD and individual data points from independent experiments are shown in each plot. Statistical analysis was carried out using one/two-way ANOVA followed by posthoc Tukey’s test.

In order to assess mitochondrial bioenergetic function in patient fibroblasts, we first used equilibration of tetramethyl rhodamine methyl ester (TMRM) to measure ΔΨ_m_ by confocal imaging (Fig. 1C and Fig. S1B). Single cell analysis of mitochondrial TMRM fluorescence intensity showed reduced ΔΨ_m_ in all the patient cells compared to controls (Fig. 1D). Quantitative analysis of mitochondrial volume occupancy by co-labelling the cells with Calcein AM, to measure cytosolic area, showed no change between the cell lines (Fig. S1C). However, quantification of mtDNA copy number revealed a significant increase in mtDNA copy number in all patient fibroblasts (Fig. S1D). Morphometric analysis of TMRM-labelled mitochondria showed that a large proportion of the mitochondrial network was fragmented while a small pool showed swollen morphology in patient cells (Fig. 1E). To investigate the cause of the reduced ΔΨ_m_ in patient fibroblasts, we measured the redox state of the NADH pool, the main substrate for the mitochondrial electron transport chain (ETC). The resting level of NADH autofluorescence was quantified through the experimental determination of the “redox index,” a ratio of the signal representing the maximally oxidized pool (response to 1 μM FCCP) and the maximally reduced pool (the response to 1 mM NaCN) (Fig. 1F and Fig. S1E). Patient fibroblasts exhibited a significant increase in the NADH redox state (i.e. less oxidised NADH/more reduced NAD^+^) in comparison to control cells, suggesting an impaired function of the mitochondrial ETC (Fig. 1G) consistent with the previous reports (Byrne et al., 2016b). To confirm this, the oxygen consumption rate was measured using the Seahorse XFe96 extracellular flux analyser (Fig. 1H). Both the ATP-linked respiratory rate and spare respiratory capacity were substantially reduced in patient cells compared to controls (Fig. 1I-J).

In order to further identify the cause of the mitochondrial bioenergetic defect observed in the patient cells, steady-state levels of ETC components were analysed by SDS-PAGE (Fig. 1K) as well as the native assembly of these components into respiratory supercomplexes by BNGE and immunoblotting (Fig. S1F). The expression levels of CI, CII and CIV were all significantly reduced as was the assembly of CI, CIII and CIV compared to control cells except for CV which remained unaffected in all patient cells (Fig. 1L and Fig. S1G). Patient cells also showed an increased rate of production of intracellular reactive oxygen species (ROS), measured using dihydroethidium (DHE) fluorescence (Fig. 1M). Together, these data show an impaired mitochondrial bioenergetic function and a respiratory chain defect associated with the decreased expression and assembly of the OXPHOS complexes in patient fibroblasts carrying pathogenic EPG5 mutations.

### Pathogenic *EPG5* variants causes dysregulation of mitochondrial calcium signalling

The Ca^2+^-dependent regulation of mitochondrial metabolism, mediated by Ca^2+^ entry into mitochondria through the MCU complex, plays a key role in driving respiratory chain activity that maintains the rate of ATP synthesis in response to increased energy demand (Szabadkai and Duchen, 2008). As mitochondrial Ca^2+^ uptake is potential-dependent, we investigated whether impaired mitochondrial Ca^2+^ uptake due to the reduced ΔΨ_m_ might amplify the bioenergetic defect in the patient cells. We therefore measured the mitochondrial matrix [Ca^2+^]_m_ response in control and patient cells expressing mitochondrial-targeted aequorin following a challenge with 10 µM histamine (Fig. 2A). Surprisingly, mitochondrial Ca^2+^ uptake was increased in patient cells with significantly larger peak of [Ca^2+^]_m_ upon physiological histamine stimulation (Fig. 2B). This striking finding suggested that the increase in histamine-induced [Ca^2+^]_m_ in patient fibroblasts could be due to altered endoplasmic reticulum (ER) Ca^2+^ content and ER-mitochondria proximity or dysregulation of the MCU/NCLX complex. To test this, cells were labelled with Fluo-4 AM and mito-Fura-2 AM (De Nadai et al., 2021; Pendin et al., 2019) to simultaneously monitor the changes in [Ca^2+^]_c_ and [Ca^2+^]_m_ (Fig. 2C-D). Application of 10 µM histamine to both cell types produced a comparable rise in [Ca^2+^]_c_ with no significant changes in total Ca^2+^ released from the ER (Fig. S2A) or its clearance as measured by the time taken from peak [Ca^2+^]_c_ to half the final baseline value (Fig. S2B). However, the increase in both resting and histamine-stimulated [Ca^2+^]_m_ was confirmed by ratiometric measurement of mito-Fura-2 intensity in patient cells (Fig. 2E-G). Notably, the initial rate of histamine-induced mitochondrial Ca^2+^ uptake was also significantly increased in patient fibroblasts (Fig. 2F). We further examined the ER-mitochondria contact sites by analysing the fraction of mitochondrial surface involved in contact with ER by transmission electron microscopy (TEM) (Fig. S2C). We found no significant change in the frequency and total number of contacts in the 0-20 nm range, the gap width relevant for Ca^2+^ transfer, between both the cell types (Fig. S2D).

**Figure 2:**
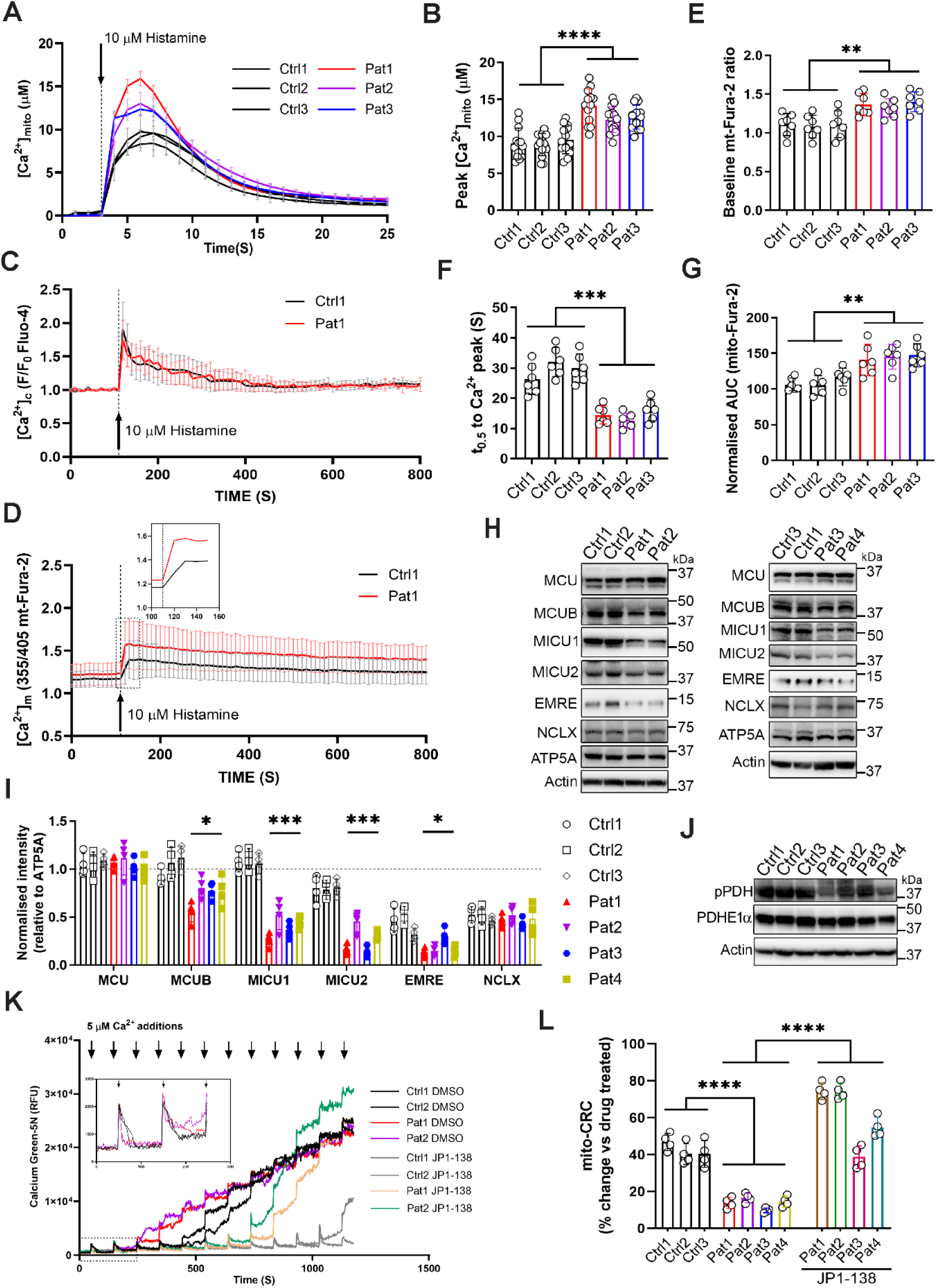
Patient-derived fibroblasts bearing pathogenic EPG5 mutations show impaired mitochondrial Ca^2+^ signalling. **(A)** Mean traces for [Ca^2+^]_m_ uptake measured using a mitochondria-target aequorin plate reader assay in response to 10 μM histamine, (n_exp_ = 3, n_rep_ = 5). **(B)** Maximum [Ca^2+^]_m_ induced by 10 μM histamine in control and patient fibroblasts, (n_exp_ = 3, n_rep_ = 5, \*\*\*\**p* <0.0001). **(C)** Mean [Ca^2+^]_c_ traces measured in control 1 and patient 1 fibroblasts upon 10 μM histamine stimulation by Fluo-4 AM, (n_exp_ = 5-7). **(D)** Mean [Ca^2+^]_m_ traces measured in control 1 and patient 1 fibroblasts upon 10 μM histamine stimulation by mito-Fura-2 AM, inset shows rate of [Ca^2+^]_m_ rise, (n_exp_ = 5-7). **(E)** Measurement of resting [Ca^2+^]_m_ levels in the control and patient fibroblasts loaded with mito-Fura-2 AM before histamine stimulation, calculated from the traces in D, (\*\**p=* 0.0065). **(F)** Quantitative analysis of time when [Ca^2+^]_m_ rise was at the half maxima upon histamine stimulation, calculated from the traces in D, (\*\*\**p=* 0.0003). **(G)** Quantification of normalised areas under the curve (AUC) of the mito-Fura-2 AM traces, representing total mitochondrial Ca^2+^ uptake over time, calculated from the traces in D, (\*\**p=* 0.0094). **(H)** Immunoblot images of proteins involved in mitochondrial Ca^2+^ signalling from whole cell lysates of all the control and patient fibroblasts. ATP5A and Actin were used as loading controls. **(I)** Protein levels relative to loading control ATP5A were normalized to those in control 1, (n_exp_ = 4, \**p=* 0.0243 and 0.0105, \*\*\**p=* 0.0002 and 0.0005). **(J)** Immunoblot images for pPDH (PDH-E1α pS293) and total PDH (PDH-E1α) from whole cell lysates of control and patient fibroblasts. Actin was used as a loading control. **(K)** Ca^2+^ retention capacity measured on isolated mitochondria from control and patient fibroblasts, mean traces showing the extra-mitochondrial calcium measured using Calcium Green-5N after repetitive addition of 5 μM CaCl_2_ boluses in the presence or absence of JP1-138 (100 nM, added at 0 s), inset shows rate of Ca^2+^ in control and patient mitochondria. **(L)** Mitochondrial Ca^2+^ retention capacity calculated as the percentage inhibition as compared to untreated mitochondria, (n_exp_ = 4, \*\*\*\**p* <0.0001). Data (A, B, C, D, E, F, G, I, and L) are expressed as mean ± SD and individual data points from independent experiments are shown in each plot. Statistical analysis was carried out using two-way ANOVA followed by posthoc Tukey’s test (non-significant *p* values are denoted with numeric values).

The increased velocity of mitochondrial Ca^2+^ uptake observed in patient fibroblasts prompted us to measure the expression levels of regulatory components of the MCU complex (Fig. 2H). MCU and NCLX protein levels were not altered but surprisingly, the expression of MICU1, MICU2 and EMRE was significantly reduced in patient cells indicating that the ‘gatekeeper’ function of MICU1/MICU2 in mitochondrial Ca^2+^ uptake is compromised in patient cells allowing rapid entry of Ca^2+^ into the mitochondria (Fig. 2I). Mitochondrial Ca^2+^ signalling directly impacts TCA cycle intermediates by the allosteric activation of pyruvate dehydrogenase (PDH) and α-ketoglutarate dehydrogenase (αKGDH). Accumulation of mitochondrial Ca^2+^ activates PDH phosphatase (PDP1), which dephosphorylates the PDH E1α subunit and thereby increases PDH activity to convert pyruvate to acetyl-CoA (Denton et al., 1972). Immunoblotting analysis of phosphorylated PDH (p-PDH E1α, inactive) revealed a significantly reduced p-PDH E1α/PDH ratio in patient cells compared to controls (Fig. 2J and Fig. S2E), consistent with the chronic elevation of resting [Ca^2+^]_m_ and enhanced PDP activity in EPG5-deficient patient cells. Remarkably, a high resting [Ca^2+^]_m_ and the associated mitochondrial bioenergetic deficiency are the characteristic cellular features of fibroblasts derived from patients with MICU1 loss-of-function mutations, a rare childhood disorder characterised by proximal myopathy, cognitive impairment and a progressive neurodegeneration (Logan et al., 2014; Lewis-Smith et al., 2016).

Supraphysiological mitochondrial Ca^2+^ accumulation can trigger the opening of mPTP and Ca^2+^ dependent cell death (Briston et al., 2019). We therefore asked whether diminished expression of the MCU regulatory proteins renders patient cells vulnerable to [Ca^2+^]_m_ overload and mPTP opening. To examine this, isolated mitochondria from control and patient fibroblasts were challenged with a series of 5 μM Ca^2+^ boluses to evoke [Ca^2+^]_m_ overload. We found that mitochondria from patient cells tolerated fewer Ca^2+^ pulses before the precipitous increase in fluorescence signal indicating mPTP opening (Fig. 2K). Consistent with the above results, the rate of Ca^2+^ uptake was also increased in patient cells in this assay, as measured by the time taken from peak Ca^2+^ to half the final baseline value (Fig. S2F). Interestingly, preincubation with JP1-138, a highly specific novel mitochondrial cyclophilin D (CypD) targeting molecule and a potent inhibitor of PTP (Pingitore et al., 2024), significantly increased the mitochondrial retention capacity in patient cells (Fig. 2K-L). Together, these data demonstrate that the downregulation of Ca^2+^-sensing regulators, MICU1and MICU2 lowers the threshold for Ca^2+^ uptake and renders patient cells more vulnerable to [Ca^2+^]_m_ overload and mPTP opening.

### Mitochondrial calcium overload triggers mtDNA release in patient-derived fibroblasts bearing pathogenic EPG5 mutations

Pathological excessive mitochondrial Ca^2+^ uptake can irreversibly lead to bioenergetic collapse, mitochondrial swelling, and mPTP opening. This catastrophic loss of mitochondrial function is a major trigger for acute cell death (Bauer and Murphy, 2020). Alternatively, limited mPTP opening in a subset of damaged mitochondria may lead to a chronic low-grade innate inflammatory response. Previous studies have suggested that mPTP opening may allow mtDNA release into the cytosol thus driving activation of innate immune signalling and inflammation, a disease hallmark of ALS and other chronic metabolic pathologies (Yu et al., 2020; Xian et al., 2022). We therefore asked whether [Ca^2+^]_m_ overload in patient cells drives this pathway. Control and patient fibroblasts immunolabelled with anti-TOM20, anti-Citrate synthetase (CS) and DNA antibodies to label OMM, IMM and mtDNA nucleoids, respectively, were imaged using Airyscan microscopy which provides near super-resolution imaging, sufficient to visualise the integrity of the mitochondrial outer and inner membrane and mtDNA nucleoids (Fig. 3A). Both cell types displayed mtDNA nucleoids contained within a continuous IMM and OMM colocalizing together (Fig. S3A). However, we found cytosolic localisation of DNA puncta in over 30% of patient cells as well as a significant increase in the average cytosolic DNA puncta per cell but none at all in control cells (Fig. 3B-C). Notably, patient cells also displayed a fragmented mitochondrial network with swollen morphology (Fig. 3A). The presence of cytosolic DNA puncta in patient cells under normal conditions supported our hypothesis that increased resting [Ca^2+^]_m_ in patient cells could cause chronic mtDNA release.

**Figure 3:**
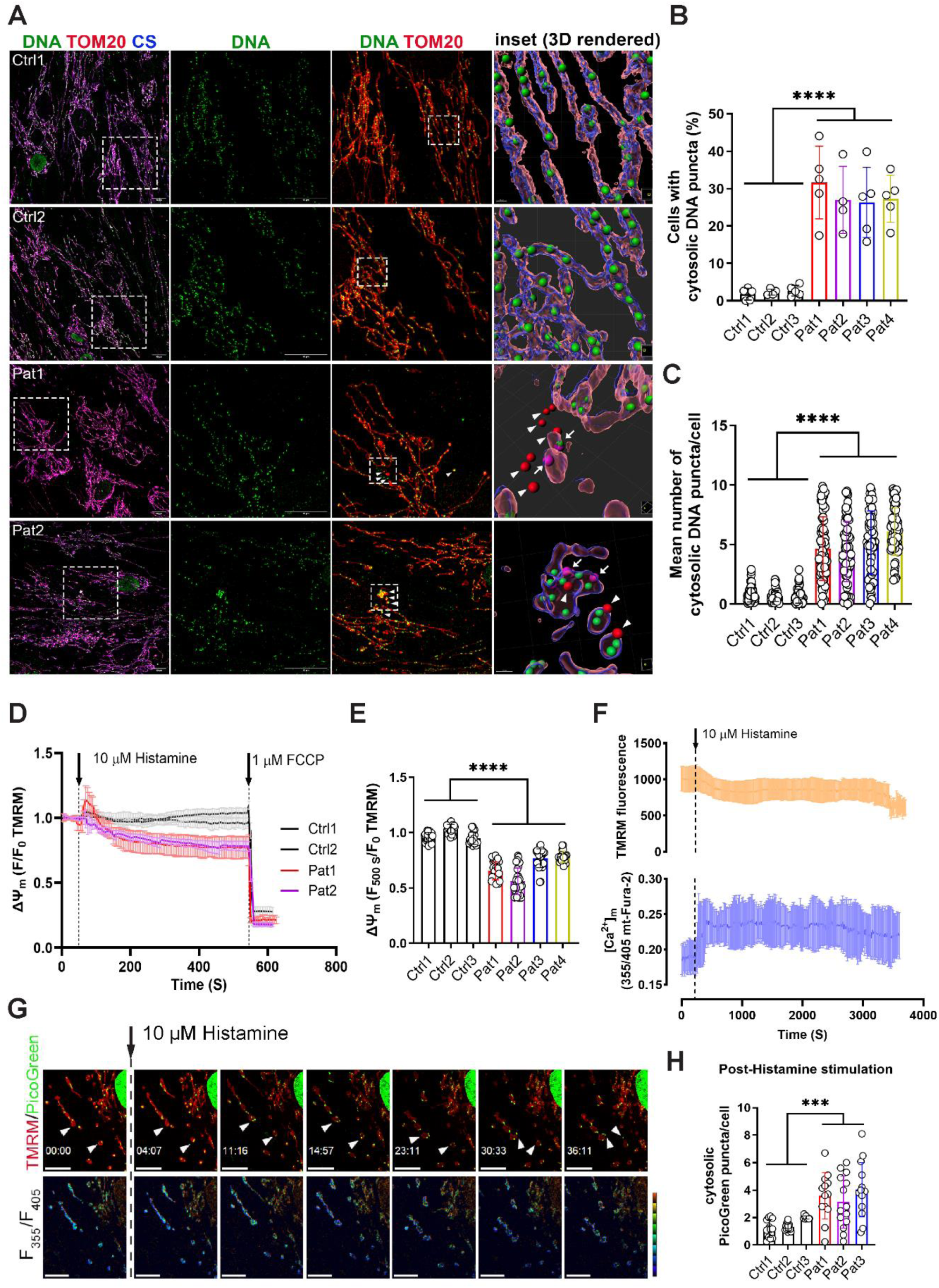
Mitochondrial Ca^2+^ overload induces the release of cytosolic mtDNA in patient-derived fibroblasts. **(A)** Representative super-resolution Airyscan images of control and patient fibroblasts immunolabelled with DNA (green), TOM20 (red) and citrate synthetase (blue). The magnified images on the right show areas in which DNA (green) does not co-localize with TOM20 (red) in patient cells. 3D representations of the inset show OMM (red), IMM (blue) and mtDNA (green), where all the mtDNA puncta (green spots) are located within the mitochondria of control fibroblasts with some mtDNA puncta (red spots indicated with arrowheads) in the cytoplasm of patient fibroblasts. mtDNA nucleoids partially outside the OMM and IMM surface are shown in magenta and indicated by arrows. Overview scale bars, 10 μm and inset scale bars, 0.5 μm. **(B)** Percentage of control and patient fibroblasts showing cytosolic DNA puncta, (n_exp_ = 4, \*\*\*\**p* <0.0001). **(C)** Quantification of number of cytosolic DNA puncta released per cell, (n_exp_ = 4, n_cells_ analysed for each control and patient = 61-71, \*\*\*\**p* <0.0001). **(D)** Traces showing mean change ± SD in TMRM (representing ΔΨ_m_) fluorescence intensity in response to 10 μM histamine challenge and FCCP-induced depolarization in control and patient fibroblasts, (n_exp_ = 4). **(E)** Measurement of the mitochondrial depolarization at 500 s after 10 μM histamine stimulation, (n_exp_ = 4, n_rep_ analysed for each control and patients = 12-14, \*\*\*\**p* <0.0001). **(F)** Mean traces of TMRM fluorescence intensity and [Ca^2+^]_m_ change after 10 μM histamine challenge in patient 1 fibroblasts co-labelled with TMRM, mito-Fura-2 and PicoGreen, (n_exp_ = 3). **(G)** Snapshots from time-lapse confocal imaging of patient fibroblasts co-labelled TMRM (red) and PicoGreen (green), upper panels and mito-Fura-2 AM, ratiometric lower panels, quantified in F. Elapsed time after 10 μM histamine challenge is indicated (Video S2). White arrowheads denote nucleoid externalization events. Scale bars, 5 μm. **(H)** Quantitative analysis of the average number of PicoGreen puncta released from TMRM-labelled mitochondria into the cytosol in response to 10 μM histamine, (n_cells_ analysed for each control and patient = 15, \*\*\**p=* 0.0006). Data (B, C, D, E, F and H) are expressed as mean ± SD and individual data points from independent experiments are shown in each plot. Statistical analysis was carried out using two-way ANOVA followed by posthoc Tukey’s test.

To recapitulate the progressive chain of events which could lead to [Ca^2+^]_m_ overload-induced mtDNA release, we first monitored ΔΨ_m_ in both control and patient fibroblasts following challenge with 10 µM histamine (Fig. 3D). Control cells maintained ΔΨ_m_ over time until the uncoupler-induced loss of ΔΨ_m_. In contrast, patient cells exhibited a steep decrease in membrane potential after the histamine challenge leading to more than a 30% reduction in ΔΨ_m_ over time (Fig. 3E). Loss of ΔΨ_m_ is one of the hallmark features of [Ca^2+^]_m_ overload-induced mPTP opening. Therefore, we next co-labelled the cells with TMRM, mito-Fura-2 and PicoGreen (a potential-dependent DNA binding probe) to monitor the intra-mitochondrial dynamics of ΔΨ_m_, [Ca^2+^]_m_ and mtDNA respectively, over a prolonged time interval of 30 min to 1 h. Live-cell imaging of control fibroblasts following histamine challenge showed a slow increase in [Ca^2+^]_m_ which coincided with a small increase in ΔΨ_m_ which was maintained over time (Fig. S3B-C). An increased fragmentation of the mitochondrial network was also observed after histamine stimulation (Video S1 and Fig. S3D) suggesting a Ca^2+^-induced remodelling of mitochondrial morphology as previously reported (Chakrabarti et al., 2018). In comparison to control cells, histamine challenge in patient cells caused a rapid increase in peak [Ca^2+^]_m_ followed by a simultaneous fall in ΔΨ_m_ and mitochondrial swelling (Fig. 3F-G and Video S2). Analyses of individual mitochondria in patient cells revealed the release of PicoGreen-labelled mtDNA associated with depolarization and swelling of mitochondria (Fig. 3H) that were never seen in control cells. Together, these data demonstrate the mechanism of [Ca^2+^]_m_ overload-induced mtDNA release in patient cells by identifying a progressive cascade of events starting from uncontrolled [Ca^2+^]_m_ uptake and overload to loss of ΔΨ_m_ and mitochondrial swelling that culminates in mPTP opening and mtDNA release.

### Release of mtDNA drives the activation of the cGAS-STING pathway and interferon response in patient-derived fibroblasts

To characterize the cell-intrinsic changes in gene expression which may impact cellular function after mtDNA release, we performed bulk RNA sequencing (RNA-seq) on a control (control 1) and a pair of patient fibroblasts (patient 1 and patient 3). Gene set enrichment analysis (GSEA) of the gene signature enriched in patient fibroblasts revealed the top upregulated genes involved in the immune response (Fig. 4A-B and Fig. S4A-B). Among the most significantly upregulated pathways, nine from the Gene Ontology biological process and seven from the KEGG pathway are related to immune activation and response. A gene expression profile of the differentially expressed genes associated with the enriched pathways, collected using a significance level of false discovery rate (FDR) <0.05, uncovered a striking upregulation of the type I/III IFN signature gene set in the patient fibroblasts (Fig. 4C and Fig. S4C). Among the list of 45 IFN-stimulated genes (ISGs), we observed an upregulation of the genes involved in direct antiviral activity (*Ifit1*, *Ifit3*, *Ifi44*, *Oasl*), members of the ISG transcription factors (*Irf9* and *Stat1*) which are activated downstream of type I/III IFN receptor and chemokines (*Ccl2* and *Cxcl10*) and pro-inflammatory cytokines (*Tnf* and *Il-1b*) involved in adaptive immune response. Upregulation of these ISGs and pro-inflammatory genes was also confirmed in the remaining patient fibroblasts with pathogenic *EPG5* variants (Fig. 4D-E and Fig. S4D) suggesting that the induction of the IFN response pathway is likely a common feature of EPG5 deficiency.

**Figure 4:**
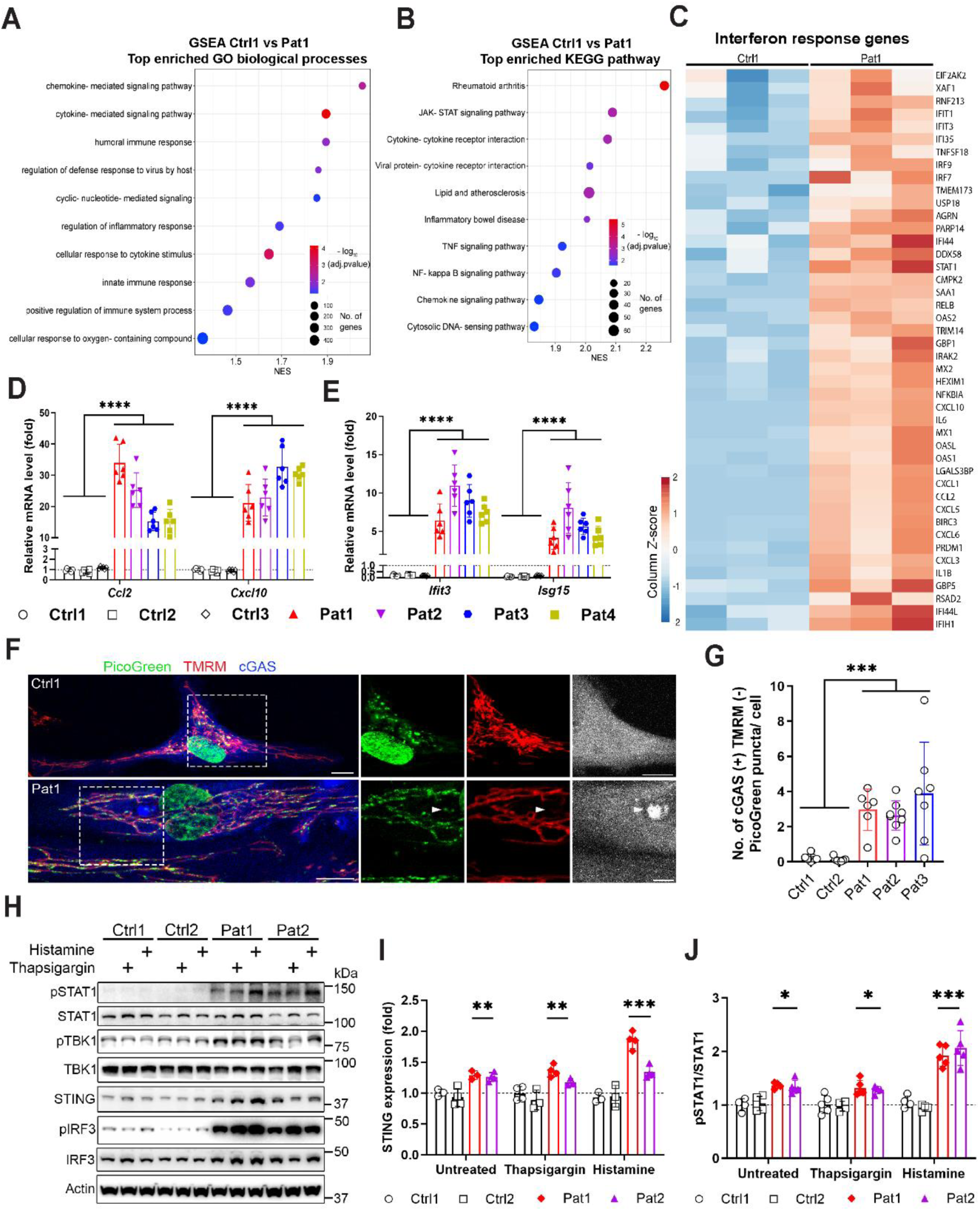
Activation of cGAS-STING pathway and interferon response triggered by cytosolic release of mtDNA in patient-derived fibroblasts. **(A-B)** Pathway analysis of RNA-seq data from patient 1 compared to control 1. Dot plots and GSEA performed using two types of gene set: the Gene Ontology (GO) biological process and KEGG pathway. NES, normalized enrichment score. **(C)** Heatmap of RNA-seq data displaying the top 50 upregulated differentially expressed type I/III IFN genes (DEGs) in control 1 and patient 1 fibroblasts, (n_exp_ = 3). **(D-E)** qRT-PCR analysis of ISGs expression in all control and patient fibroblasts, (n_exp_ = 3, \*\*\*\**p* <0.0001). **(F)** Representative confocal images of control 1 and patient 1 fibroblasts transfected with BFP-cGAS and co-labelled with TMRM (red) and PicoGreen (green). The magnified images on the right show areas in which a mtDNA nucleoids (green) devoid of TMRM fluorescence co-localize with cytosolic cGAS puncta (grey) in patient cells indicated by a white arrowhead. Overview scale bars, 10 μm and inset scale bars, 5 μm. **(G)** Quantification of the number of cGAS(+) and TMRM (-) PicoGreen puncta per cell, (n_exp_ = 3, n_cells_ analysed for each control and patient = 26-30, \*\*\**p=* 0.0004). **(H)** Immunoblot images of proteins involved in the cGAS-STING signalling and ISGs induction from whole cell lysates of control and patient fibroblasts treated with 10 μM histamine and 1 μM thapsigargin for 24 h. Actin was used as a loading control. **(I)** Protein expression levels of STING normalized and plotted as fold difference relative to control, (n_exp_ = 4, \*\**p=* 0.0056 and 0.0096, \*\*\**p=* 0.0006). **(J)** The ratio of pSTAT1 (p Tyr701) and total STAT1 band intensities normalised to the control 1 ratio, (n_exp_ = 4, \**p=* 0.124 and 0.0161, \*\*\**p=* 0.0008). Data (D, E, G, I and J) are expressed as mean ± SD and individual data points from independent experiments are shown in each plot. Statistical analysis was carried out using two-way ANOVA followed by posthoc Tukey’s test.

To investigate the signalling events upstream of IFN induction in patient fibroblasts, we asked if the presence of cytosolic mtDNA could induce type I IFN by activating the canonical cGAS-STING pathway. cGAS is predominantly a nuclear protein, however, immunofluorescence analysis revealed a significant increase in the intensity of the cytosolic cGAS in patient cells (Fig. S4E-F) indicating its cytosolic relocalization in response to immunogenic intracellular DNA (de Oliveira Mann and Hopfner, 2021). Live-cell imaging of fibroblasts over-expressing BFP-cGAS and co-labelled with TMRM and PicoGreen showed cytosolic cGAS puncta colocalizing with free mtDNA devoid of mitochondria in patient cells (Fig. 4F-G) supporting the notion that cytosolic cGAS observed in patient cells binds to mtDNA after its release from mitochondria. Cytosolic detection of mtDNA by cGAS activates classic STING signalosomes at the Golgi apparatus which in turn activates TBK1, a central kinase involved in the integration of innate immune signals from cytosolic DNA/RNA sensors to IFN induction by activating interferon-regulatory factors (IRFs) (Liu et al., 2015; Zhang et al., 2019). In control and patient fibroblasts treated with histamine or thapsigargin, to increase [Ca^2+^]_c_ and amplify [Ca^2+^]_m_ overload, immunoblotting for active pTBK1 showed a substantial increase both under resting and histamine/thapsigargin-stimulated conditions in patient compared to control cells (Fig. 4H and Fig. S4G-H). Consistent with TBK1 activation, a significant increase in STING expression was also detected in patient fibroblasts which further increased with histamine challenge (Fig. 4I and Fig. S4I). Notably, we observed a robust activation of downstream transcription factors IRF3 and STAT1 in patient cells under resting condition which increased by more than two-fold after histamine challenge (Fig. 4J and Fig. S4J-K). Together, these results support the RNA-Seq and mRNA expression data and suggests that the cGAS-STING signalling is constitutively active in patient-derived fibroblasts and further amplified by increased [Ca^2+^]_m_ during physiological histamine stimulation.

### Inhibition of mPTP opening by JP1-138 attenuates STING-dependent IFN response in patient-derived fibroblasts

Given that the constitutive activation of cGAS-STING signalling drives the downstream IFN pathway in patient fibroblasts, we reasoned that STING inhibition might curtail this inflammatory response. Treatment with H-151, a covalent inhibitor that blocks activation-induced palmitoylation of STING (Haag et al., 2018), dampened the IFN response signalling in patient cells by reducing STING levels and normalizing active IRF3 and STAT1 comparable to the levels seen in control cells (Fig. S5A-D). However, treatment with G140, a cGAS-specific inhibitor (Lama et al., 2019) showed only a marginal effect on STING, pIRF3 and pSTAT1 levels in patient cells (Fig. S5A-D). Furthermore, H-151 treatment also substantially reduced the expression levels of ISGs in patient cells (Fig. S5E-F) indicating that STING inhibition might represent a potential therapeutic target in patients with pathogenic *EPG5* variants.

We next asked whether the STING-IFN response pathway might act in a feedback manner to further impair mitochondrial bioenergetic function in patient fibroblasts. Patient fibroblasts exhibited no significant change in resting ΔΨ_m_ after 24 h treatment with H-151 (Fig. 5A). Similarly, the NADH redox index remained unaltered in all the patient cells (Fig. 5B). Long-term treatment with H-151 for three days failed to show any significant increase in resting and maximal respiration compared to untreated patient cells (Fig. 5C and Fig. S5G-I). These results suggest that the inflammatory signature observed in patient fibroblasts is a consequence of impaired mitochondrial Ca^2+^ homeostasis but does not contribute to the underlying bioenergetic defect of *EPG5*-related disorders.

**Figure 5:**
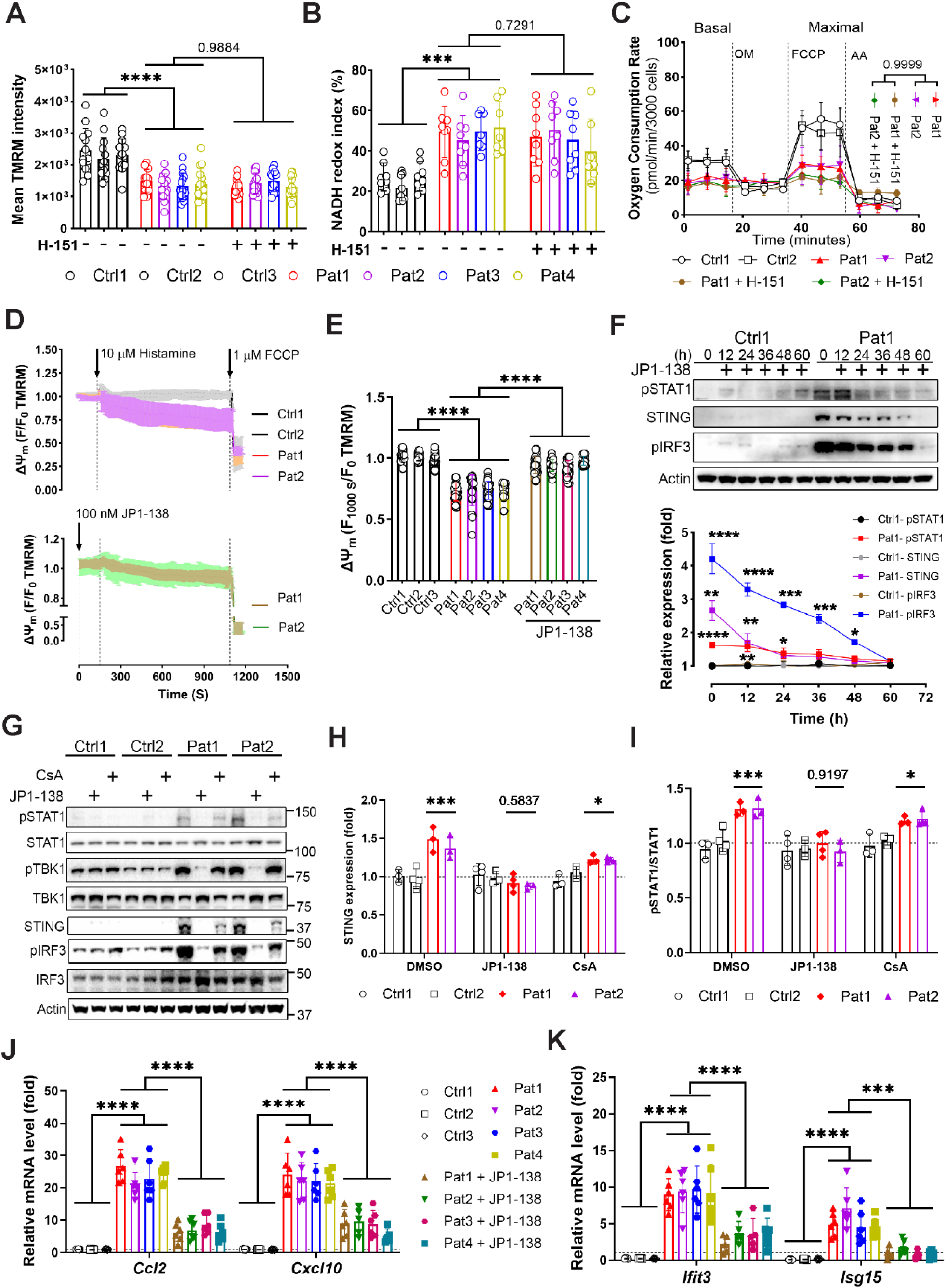
Attenuation of STING-dependent interferon response by JP1-138 treatment in patient-derived fibroblasts. **(A)** Quantitative analysis of steady state ΔΨ_m_ by measuring TMRM fluorescence intensity in control and patient fibroblasts either untreated or treated with STING inhibitor, H-151 (1 μM, 24 h) (n_exp_ = 3, n_cells_ analysed for each control and patient = 12-15, \*\*\*\**p* <0.0001). **(B)** NADH redox index of control and patient fibroblasts either untreated or treated with H-151, (n_exp_ = 3, n_cells_ analysed for each control and patient = 7-10, \*\*\**p=* 0.0001). **(C)** Normalised OCR traces from controls and patient fibroblasts either untreated or treated with H-151 for longer durations (0.5 μM, 3 days) (n_exp_ = 3, n_rep_ = 4). **(D)** Traces showing mean change ± SD in ΔΨ_m_ in response to 10 μM histamine challenge and FCCP-induced depolarization in the absence (upper panel) and presence of JP1-138 (100 nM) (lower panel) in control and patient fibroblasts, (n_exp_ = 4). **(E)** Quantification of the mitochondrial depolarization at 1000 s after 10 μM histamine stimulation, (n_exp_ = 4, n_rep_ analysed for each control and patient = 36-44, \*\*\*\**p* <0.0001). **(F)** Immunoblot images of cGAS-STING signalling proteins from whole cell lysates of control 1 and patient 1 fibroblasts treated with JP1-138 (100 nM) for the indicated time. Actin was used as a loading control. Relative expression levels of pSTAT1 (p Tyr701), STING and pIRF3 (pSer396) normalized and plotted as fold difference relative to control, (n_exp_ = 3, \**p=* 0.0235, \*\**p=* 0.0049, \*\*\**p=* 0.0007, \*\*\*\**p* <0.0001). **(G)** Immunoblot images of the cGAS-STING signalling proteins from whole cell lysates of control and patient fibroblasts treated with either CsA (1 μM) or JP1-138 (100 nM) for 3 days. Actin was used as a loading control. **(H)** Protein expression levels of STING normalized and plotted as fold difference relative to control, (n_exp_ = 4, \**p=* 0.0265, \*\*\**p=* 0.0004). **(I)** The ratio of pSTAT1 (p Tyr701) band intensities normalised to the control 1 ratio, (n_exp_ = 4, \**p=* 0.0197, \*\*\**p=* 0.0007). **(J-K)** qRT-PCR analysis of ISGs expression in all the control and patient fibroblasts either untreated or treated with JP1-138 (100 nM) for 3 days (n_exp_ = 3, \*\*\**p=* 0.0001, \*\*\*\**p* <0.0001). Data (A, B, C, D, E, F, H, I, J and K) are expressed as mean ± SD and individual data points from independent experiments are shown in each plot. Statistical analysis was carried out using one/two-way ANOVA followed by posthoc Tukey’s test (non-significant *p* values are denoted with numeric values).

Based on our finding that mPTP inhibition by JP1-138 led to more than 50% increase in mitochondrial calcium retention capacity of patient cells, we hypothesized that treatment with JP1-138 could also suppress downstream activation of the cGAS-STING signalling and IFN response by protecting patient cells from [Ca^2+^]_m_ overload and mtDNA release. To test this, we again monitored ΔΨ_m_ in both control and patient fibroblasts following a challenge with 10 µM histamine (Fig. 5D and Fig. S6A). As expected, patient cells showed a rapid loss of ΔΨ_m_ in comparison to control cells. However, preincubation with JP1-138 completely rescued the histamine-induced mitochondrial depolarization in patient cells (Fig. 5E) suggesting that treatment with JP1-138 increases [Ca^2+^]_m_ buffering capacity in intact patient fibroblasts. Next, control and patient fibroblasts were treated with JP1-138 for different time points and immunoblotted to detect changes in the activation of the cGAS-STING signalling (Fig. 5F). Interestingly, JP1-138 treatment was associated with a time-dependent decrease in the expression of STING, pIRF3 and pSTAT1 reaching levels seen in control cells after 60 h. Using this time point, we further tested the efficacy of JP1-138 in supressing cGAS-STING signalling by comparing it to another classic mPTP inhibitor, cyclosporin A (CsA) (Fig. 5G and Fig. S6B). Treatment with JP1-138 (100 nM) restored the activation of TBK1, IRF3 and STAT1 and reduced the expression of STING to normal levels in patient cells (Fig. 5H-I and Fig. S6C-F). In comparison, CsA treatment showed only a partial response and failed to effectively suppress the cGAS-STING signalling when used at a higher concentration (1 µM) demonstrating the specificity and potent activity of JP1-138 in comparison to CsA. Importantly, JP1-138 treatment also substantially reduced the expression levels of ISGs in patient cells (Fig. 5J-K). Together, these data show that JP1-138 treatment increases [Ca^2+^]_m_ buffering capacity and protects patient cells from [Ca^2+^]_m_ overload-induced mPTP opening and activation of the cGAS-STING signalling.

### JP1-138 suppresses mtDNA release and improves mitochondrial bioenergetic function in patient fibroblasts bearing pathogenic EPG5 mutations

To determine whether inhibition of mPTP by JP1-138 treatment also supressed chronic mtDNA release, control and patient fibroblasts immunolabelled with TOM20, CS and DNA were examined using Airyscan imaging (Fig. 6A and Fig. S7A). As expected, patient cells showed a significant increase in the cytosolic localization of DNA puncta as well as an increased proportion of the mitochondrial pool with fragmented and swollen morphology. Swollen mitochondria often displayed mtDNA extrusion in patient cells. Long term treatment with JP1-138 lead to a marked reduction in both the average number and the total percentage of cells with cytosolic DNA puncta in patient fibroblasts (Fig. 6B-C). Notably, we also observed remodelling of mitochondrial morphology in patient cells characterized by a significant reduction in the swollen mitochondria pool (Fig. 6D) suggesting that JP1-138 treatment rescues the mitochondrial bioenergetic defect in patient fibroblasts. Resting ΔΨ_m_ in patient cells increased to the levels seen in control cells after long-term treatment with JP1-138 (Fig. 6E-F and Fig. S7B) indicating a significant repolarization of mitochondria in patient fibroblasts. The increase in ΔΨ_m_ was accompanied by a decrease in the NADH redox index (i.e. more oxidised NADH/less reduced NAD^+^) in comparison to untreated patient fibroblasts (Fig. 6G). Improvement in these bioenergetic parameters was associated with a concomitant increase in mitochondrial respiration in patient fibroblasts (Fig. 6H and Fig. S7C). Long-term treatment with JP1-138 led to a small but significant increase in basal or ATP-linked respiration rate along with a large increase in the spare respiratory capacity when compared to the untreated patient group (Fig. S7D-E). However, compared to the changes in ΔΨ_m_ and NADH redox index, the increase in mitochondrial respiration after JP1-138 treatment was partial when compared with untreated control fibroblasts. Together these data show that JP1-138 treatment effectively suppresses mtDNA release and partially rescues mitochondrial bioenergetic function and morphology in patient fibroblasts carrying pathogenic EPG5 mutations.

**Figure 6:**
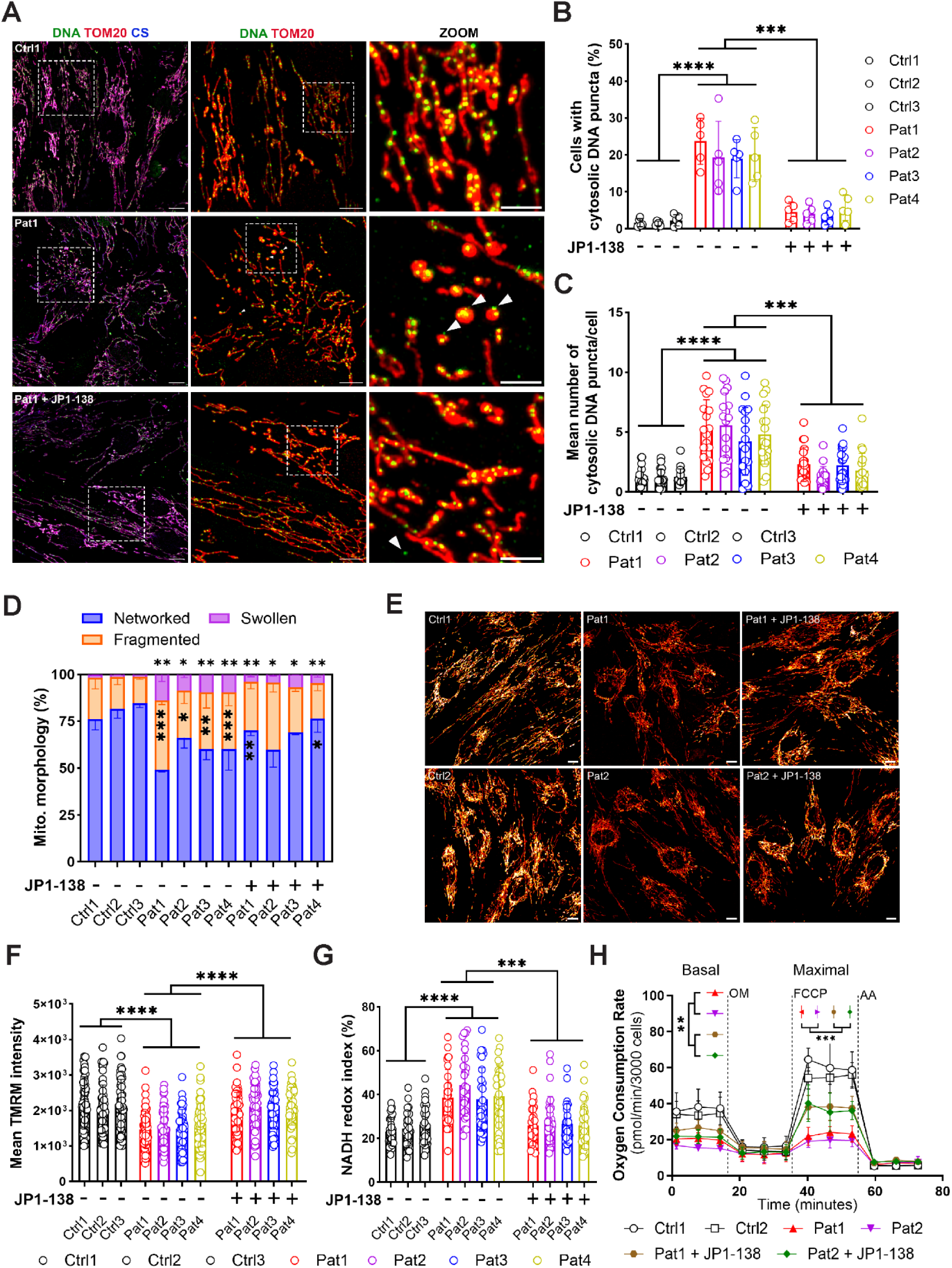
JP1-138 treatment reduces cytosolic mtDNA release and improves mitochondrial bioenergetic function in patient fibroblasts bearing pathogenic *EPG5* variants. **(A)** Representative Airyscan images of control 1 and patient 1 fibroblasts treated with either JP1-138 (100 nM) or vehicle (DMSO) for 3 days and immunolabelled with DNA (green), TOM20 (red) and citrate synthetase (blue). The magnified images on the right show areas in which DNA (green) does not co-localize with TOM20 (red) in patient cells marked by white arrow heads which is significantly reduced in JP1-138-treated patient fibroblasts. Overview scale bars, 10 μm and inset scale bars, 5 μm and 2.5 μm. **(B)** Percentage of control and patient fibroblasts treated with either JP1-138 (100 nM) or vehicle (DMSO) for 3 days and analysed for cytosolic DNA puncta, (n_exp_ = 5, \*\*\**p=* 0.0001, \*\*\*\**p* <0.0001). **(C)** Quantification of the number of cytosolic DNA puncta released per cell, calculated from B (n_exp_ = 4, n_cells_ analysed for each control and patient = 16-21, \*\*\**p=* 0.0001, \*\*\*\**p* <0.0001). **(D)** Mitochondrial morphometric analysis of all Airyscan confocal images in A classified into networked, fragmented and swollen mitochondria represented as percentage of total mitochondrial population, (n_exp_ = 4, n_cells_ analysed for each control and patient = 16-21, \**p=* 0.0140, \*\**p=* 0.0028, \*\*\**p=* 0.0006). **(E)** Representative confocal image of TMRM labelled control and patient fibroblasts treated with either JP1-138 (100 nM) or vehicle (DMSO) for 3 days. Scale bars: 20 μm. **(F)** Mean TMRM values showing steady state ΔΨ_m_, (n_exp_ = 4, n_cells_ = analysed for each control and patient = 42-59, \*\*\*\**p* <0.0001). **(G)** NADH redox index of control and patient fibroblasts treated with either JP1-138 (100 nM) or vehicle (DMSO) for 3 days (n_exp_ = 3, n_cells_ analysed for each control and patient = 27-31, \*\*\**p=* 0.0001, \*\*\*\**p* <0.0001). **(H)** Normalised OCR traces from controls and patient fibroblasts treated with either JP1-138 (100 nM) or vehicle (DMSO) for 3 days (n_exp_ = 3, n_rep_ = 4, \*\**p=* 0.0077, \*\*\**p=* 0.0009). Data (B, C, D, F, G, and H) are expressed as mean ± SD and individual data points from independent experiments are shown in each plot. Statistical analysis was carried out using one/two-way ANOVA followed by posthoc Tukey’s test.

### Impaired mitochondrial Ca^2+^ signalling sensitises Q336R iPSC-derived neurons to excitotoxicity

To investigate the impact of impaired mitochondrial bioenergetic function and mitochondrial Ca^2+^ signalling more specifically on the pathophysiology of neurodegeneration that is a severe feature of patients with *EPG5*-related disorders, we used an iPSC-derived cortical neuronal model, reprogrammed from patient 1 fibroblasts bearing the pathogenic founder Q336R missense mutation and a CRISPR/Cas9-edited isogenic control. In vitro neurodifferentiation of isogenic and Q336R iPSCs yielded mixed cultures of neurons and glial cells with no significant change in the proportion of cell types (Fig. S8A-B). The increased bioenergetic demand imposed during metabolic workload such as physiological glutamate-induced excitation may sensitize neurons with impaired mitochondrial function to glutamate excitotoxicity (Plotegher et al., 2021). To test this, mixed cultures of neurons and glial cells, loaded with low-affinity [Ca^2+^]_c_ indicator FuraFF-AM were challenged with near physiological (10 µM) and toxic (100 µM) glutamate concentrations (Fig. 7A-B). Early peak [Ca^2+^]_c_ (ΔFuraFF_early_) remained unchanged between isogenic and Q336R neurons in response to 10 µM glutamate (Fig. 7C). However, in the majority of Q336R neurons but never in the isogenic control cells, the initial transient response was followed by a progressive secondary increase of [Ca^2+^]_c_ (Fig. 7D-E). This characteristic response, referred to as delayed calcium deregulation (DCD) was evident only at 100 µM in control cells (Fig. 7A). Glutamate-induced DCD drives excitotoxic neuronal cell death through a cascade of pathways (Tymianski, 2011; Llorente-Folch et al., 2015). Disruption of mitochondrial Ca^2+^ homeostasis is the key contributing factor during glutamate excitotoxicity (Plotegher et al., 2021). Therefore, we first measured the expression of the regulatory components of the MCU complex using immunoblotting (Fig. 7F). As seen in the patient fibroblasts, protein expression of MICU1 and MICU3, a brain-specific Ca^2+^ sensing regulator, were significantly reduced in Q336R neurons along with EMRE, while the relative levels of MCU and NCLX remained unchanged (Fig. 7G). BN-PAGE analysis of mitochondria isolated from isogenic and Q336R neuronal cultures revealed the ∼1.1 MDa assembly representing MCU complex containing gatekeeper subunits in isogenic mitochondria and a prominent constitutively active ∼400 kDa MCU-EMRE complex devoid of gatekeeper proteins in Q336R mitochondria (Fig. S8C-E) as previously described in cortical mitochondria of *Afg3l2* neuron-specific knockout mice with spastic ataxia-neuropathy (Konig et al., 2016). In addition to defective neuronal MCU assembly, native complex I assembly was also reduced in Q336R mitochondria (Fig. S8C). Analysis of mRNA expression showed no significant change among the components of the MCU complex (Fig. S8F) suggesting a post-translational regulation of MICU1/MICU3 expression/assembly in Q336R neurons.

**Figure 7:**
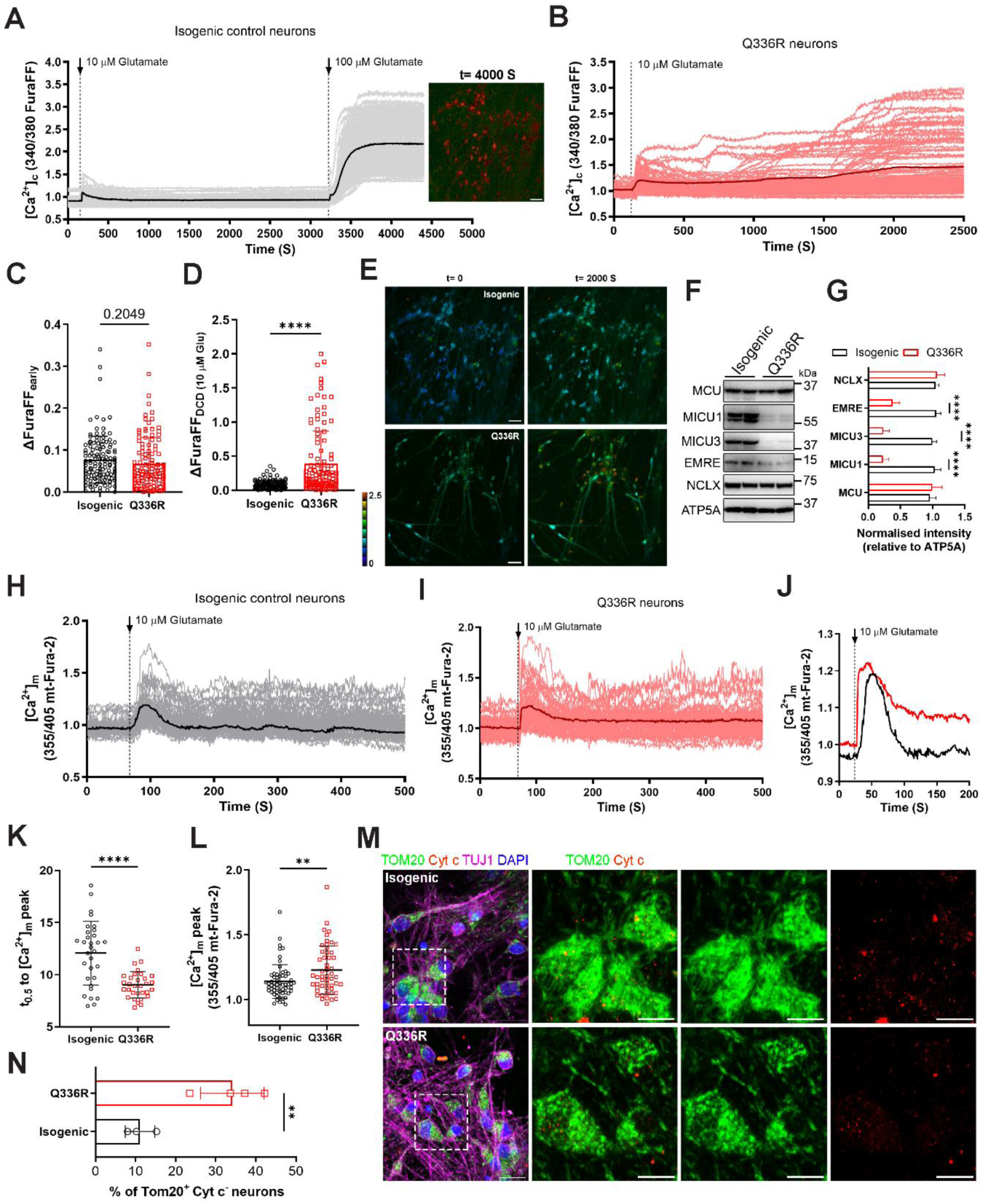
Enhanced susceptibility to glutamate-induced delayed Ca^2+^ dysregulation and cell death in Q336R neurons. **(A-B)** Cytosolic calcium concentration in neurons labelled with the low-affinity calcium sensor, FuraFF AM and measured by fluorescence imaging for the indicated time intervals. Changes in [Ca^2+^]_c_ measured from individual (grey) and mean (black) traces, following sequential exposure of isogenic neurons to 10 μM and 100 μM glutamate are plotted as a function of time. The traces reveal a significant difference between the responses of isogenic and Q336R neurons upon 10 μM glutamate stimulation which was followed by delayed calcium deregulation (DCD) in a large proportion of Q336R neurons, (n_exp_ = 3, n_cells_ analysed for isogenic and Q336R = 91-120). **(A)** Baseline subtracted peak values of the early response at about 150 s to 10 μM glutamate for isogenic and Q336R neurons. **(B)** Baseline subtracted peak values of DCD at 2000 s in response to 10 μM glutamate, (\*\*\*\**p* <0.0001). **(C)** FuraFF ratiometric images for isogenic and Q336R neurons shown at the start of the experiment (t = 0 s) and at 2000 s after exposure to glutamate, showing the full recovery of [Ca^2+^]_c_ in isogenic control neurons, while the sustained very high [Ca^2+^]_c_ levels in the Q336R neurons reflecting DCD, similar to response after 100 μM glutamate in isogenic control in A. Scale bars, 50 μm. **(D)** Immunoblot images of proteins involved in mitochondrial Ca^2+^ signalling from whole cell lysates of isogenic and Q336R neurons. ATP5A was used as a loading control. **(E)** Protein levels relative to loading control ATP5A were normalized to those in control 1, (n_exp_ = 4, \*\*\*\**p* <0.0001). **(H-I)** Mitochondrial Ca^2+^ concentration in neurons labelled with the mito-Fura-2 AM and measured by fluorescence imaging for the indicated time intervals. Changes in [Ca^2+^]_m_ measured from individual grey traces in isogenic control and red traces in Q336R neurons following exposure to 10 μM glutamate are plotted as a function of time, (n_exp_ = 3, n_cells_ analysed for isogenic and Q336R = 55-75). **(J)** Mean [Ca^2+^]_m_ traces measured with mito-Fura-2 upon exposure to 10 μM glutamate and calculated from individual traces in H and I. **(K)** Time when [Ca^2+^]_m_ rise was at the half maxima upon glutamate stimulation, calculated from the traces in H and I, (\*\*\*\**p* <0.0001). **(L)** Maximum [Ca^2+^]_m_ rise induced exposure to 10 μM glutamate, calculated from the traces in H and I, (\*\**p=* 0.0041). **(M)** Representative confocal image of isogenic control and Q336R neurons treated with either 10 μM glutamate or vehicle (PBS) for 12 h and immunolabelled with TOM20, Cytochrome c (Cyt c) and neuronal β-Tubulin III, Clone TUJ1. The magnified images on the right show areas in which TOM20 (green) does not co-localize with Cyt c (red) in Q336R neurons. Overview scale bars, 20 μm, inset scale bars, 5 μm. **(N)** Quantitative analysis of percentage of total mitochondrial area lacking Cyt c, (n_exp_ = 3, n_cells_ analysed for isogenic and Q336R = 32-40, \*\**p=* 0.0059). Data (C, D, G, K, L and N) are expressed as mean ± SD and individual data points from independent experiments are shown in each plot. Statistical analysis was carried out either using two-tailed unpaired Student’s t-test or one-way ANOVA followed by posthoc Tukey’s test (non-significant *p* values are denoted with numeric values).

Consistent with these results, the measurement of [Ca^2+^]_m_ in isogenic and Q336R neurons loaded with mito-Fura-2 AM, showed a higher resting [Ca^2+^]_m_ in Q336R neurons (Fig. S8G). Moreover, confocal images analysis of ratiometric images (F_355_/F_405_) revealed a swollen mitochondrial pool with a high F_355_/F_405_ ratio in Q336R neurons compared to isogenic control neurons which showed elongated mitochondrial morphology with a low F_355_/F_405_ ratio (Fig. S8H). Exposure to a physiological concentration of glutamate evoked an early rise in [Ca^2+^]_m_ and a larger [Ca^2+^]_m_ peak in Q336R neurons compared to isogenic control neurons (Fig. 7H-L). Ratiometric image analysis also showed a rapid [Ca^2+^]_m_ rise in Q336R neurons in response to glutamate challenge (Fig. S8I). Glutamate-induced [Ca^2+^]_m_ overload in Q336R neurons was confirmed using the low-Ca^2+^ affinity mito-aequorin probe (Fig. S8J). These results are consistent with [Ca^2+^]_m_ overload as a cause of excitotoxic glutamate-induced DCD and neuronal cell death. Therefore, we analysed cytochrome c (Cyt c) release in neurons exposed to 10 µM glutamate overnight (12 h) by immunolabelling neurons for the TOM20 and Cyt c (Fig. 7M). The release of cytochrome c from mitochondria (TOM20**^+^** and Cyt c**^-^**) is the major trigger for caspase activation during the initial events of apoptotic cell death cascade. After glutamate challenge, Q336R neurons showed a marked increase in TOM20 intensity without Cyt c intensity suggesting Cyt c release in comparison to isogenic control neurons (Fig. 7N). Notably, Q336R neurons also showed an aberrant remodelling of mitochondrial morphology accompanied by a significant increase in swollen mitochondria pool in the somas and a more fragmented form in the axons (Fig. S8K). Altogether, these results show that impaired mitochondrial Ca^2+^ signalling and the ensuing [Ca^2+^]_m_ overload increases the vulnerability to glutamate-induced DCD and cell death in Q336R neurons.

### JP1-138 improves mitochondrial Ca^2+^ buffering capacity and bioenergetic function in Q336R neurons

An increase in [Ca^2+^]_m_ is the most potent inducer of mPTP opening and therefore provides a direct link between sustained increases in [Ca^2+^]_m_ and mitochondrial dysfunction during excitotoxic injury in neurons (Keelan et al., 1999; Abramov and Duchen, 2008). Given that the pharmacological inhibition of mPTP in patient fibroblasts suppresses mtDNA-induced IFN response and partially restores mitochondrial bioenergetic function, we first determined mitochondrial Ca^2+^ retention capacity of neurons which were digitonin permeabilised after co-labelling with Fluo-4 AM (which is lost from the cytosol but retained in mitochondria after permeabilization (McKenzie et al., 2017) and TMRM to monitor dynamic changes in [Ca^2+^]_m_ and ΔΨ_m,_ respectively, using sequential additions of exogenous CaCl_2_ as described in Fig. S9A. The highest fold-change in Fluo-4 intensity was evident only after the fourth successive Ca^2+^ addition (6.91 µM final Ca^2+^) in permeabilised isogenic neurons, indicating the Ca^2+^ uptake threshold and cooperative activation of MCU by Ca^2+^ (Fig. 8A). Subsequent Ca^2+^ addition (13.3 µM final Ca^2+^) induced permeability transition culminating in the collapse of ΔΨ_m_ as indicated by the rapid loss of the TMRM signal. In contrast, mitochondria from Q336R neurons showed an almost linear increase in Fluo-4 intensity with successive Ca^2+^ additions indicating a lower threshold for activation and a constitutively active MCU complex. As a consequence of [Ca^2+^]_m_ overload, mPTP opening and rapid dissipation of ΔΨ_m_ was observed at lower concentrations of Ca^2+^ (between 1.32 and 3.28 µM) in the Q336R cells (Fig. 8B). Notably, the preincubation of Q336R mitochondria with JP1-138 significantly increased the threshold of mPTP opening and the loss of ΔΨ_m_ without affecting the rate of Ca^2+^ uptake indicating the protective effect of JP1-138 from [Ca^2+^]_m_ overload-induced mPTP opening and mitochondrial depolarisation (Fig. 8A-B). To demonstrate this under (patho)physiological state, isogenic and Q336R neurons were loaded with Rhodamine-123 in ‘dequench mode’ to monitor time-dependent changes in ΔΨ_m_ following exposure to glutamate. After an initial transient depolarization indicated by an early rise in fluorescence in response to glutamate, ΔΨ_m_ recovered to the baseline in almost all the isogenic neurons (Fig. 8C and 8F). In contrast, Q336R neurons showed a delayed secondary depolarisation in the majority of the neurons (Fig. 8D), coincident with the glutamate-induced DCD response (Abramov and Duchen, 2008). Moreover, an uncoupler-induced mitochondrial depolarization caused a smaller additional depolarization in comparison to isogenic neurons confirming near total dissipation of ΔΨ_m_ induced by glutamate in the Q336R neurons. Importantly, preincubation with JP1-138 completely protected the Q336R neurons from glutamate-induced collapse of ΔΨ_m_ (Fig. 8E-G). To determine the bioenergetic differences under unstimulated resting state, we measured basal and maximal respiration in isogenic and Q336R neurons. Both ATP-linked respiration rate and spare respiratory capacity were reduced in Q336R neurons (Fig. 8H). However, long-term treatment with JP1-138 resulted in a small but significant increase in both basal and maximal respiration rate in Q336R neurons (Fig. 8I-J). Together, these results demonstrate that JP1-138 effectively increases mitochondrial Ca^2+^ buffering capacity and protects Q336R neurons from glutamate-induced loss of ΔΨ_m_, and improves mitochondrial respiration, thus highlighting a potential therapeutic strategy to target neurodegeneration in *EPG5*-related disorders.

**Figure 8:**
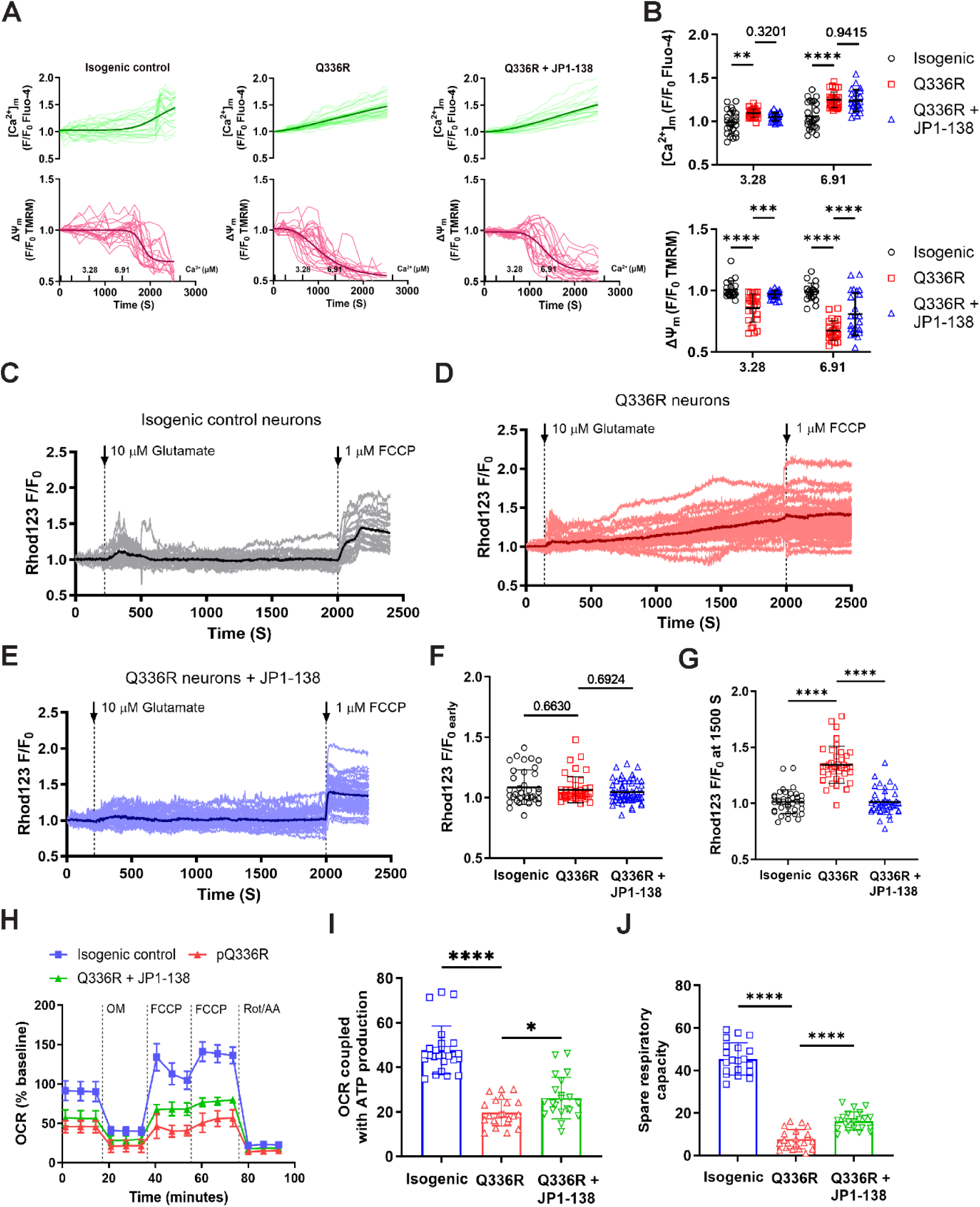
JP1-138 increases mitochondrial Ca^2+^ buffering capacity and partially rescues bioenergetic function in Q336R neurons. **(A)** Traces showing mean change ± SD in Fluo-4 AM (representing [Ca^2+^]_m_) and TMRM (representing ΔΨ_m_) fluorescence intensity in response to increasing concentrations of exogenous Ca^2+^ (upward tick on the x-axis) to digitonin-permeabilised isogenic and Q336R neurons bathed in a pseudo-intracellular recording solution with or without JP1-138 (0.5 μM). **(B)** Quantitative analysis of Fluo-4 intensity (upper panel) and TMRM fluorescence (lower panel) showing peak change in permeabilised isogenic and Q336R neurons treated with or without JP1-138 in response to ascending concentrations of Ca^2+^ in the recording buffer, (n_exp_ = 3, n_cells_ analysed for isogenic and Q336R = 32-45, \*\**p=* 0.0050, \*\*\**p=* 0.0005, \*\*\*\**p* <0.0001). **(C-E)** Mitochondrial membrane potential measured using Rhodamine-123 (with the ‘dequench protocol’) in responses to exposure to 10 μM glutamate stimulation and plotted as a function of time for isogenic and Q336R treated with either JP1-138 (0.5 μM) or vehicle (DMSO). An increase in Rhodamine-123 fluorescence intensity reports mitochondrial depolarization. The responses to 10 μM glutamate were significantly different between isogenic and Q336R revealing a large depolarization in the Q336R neurons. Pre-incubation with JP1-138 completely rescues glutamate-induced depolarisation in Q336R neurons. FCCP-induced mitochondrial depolarization represents total ΔΨ_m_ after glutamate stimulation, (n_exp_ = 3, n_cells_ analysed for isogenic and Q336R = 42-55). **(F-G)** Quantitative analysis of the mitochondrial depolarization at early time point and 1500 s after 10 μM glutamate stimulation, expressed as Rhod123 F/F_0_ for isogenic and Q336R neurons with or without JP1-138 pretreatment, (\*\*\*\**p* <0.0001). **(F)** Normalised OCR traces from isogenic and Q336R neurons treated with either JP1-138 (0.5 μM) or vehicle (DMSO) for 3 days (n_exp_ = 3, n_rep_ = 4). **(I-J)** Normalised ATP-linked respiration and spare reserve capacity of isogenic and Q336R neurons calculated from traces in G (n_exp_ = 3, n_rep_ = 4, \**p=* 0.0177, \*\*\*\**p* <0.0001). Data (B, F, G, H, I and J) are expressed as mean ± SD and individual data points from independent experiments are shown in each plot. Statistical analysis was carried out using one-way ANOVA followed by posthoc Tukey’s test.

## Discussion

Whilst defective autophagy has been identified as a key mechanism in *EPG5*-related disorders, downstream pathogenic effects remain poorly understood. Our recent findings suggested impaired mitochondrial homeostasis due to defective PINK1/PARKIN-dependent mitophagy in patient-derived cells (Dafsari et al., 2024). Here, we show that impaired mitochondrial Ca^2+^ signalling and the associated bioenergetic insufficiency could be major contributing factors to the neurodegeneration observed in patients with *EPG5*-RD. We found that mitochondrial Ca^2+^ overload and increased vulnerability to mPTP opening drives mtDNA release and the cGAS-STING dependent IFN response in patient fibroblasts bearing pathogenic *EPG5* variants. Patients with either truncating or missense mutations showed almost complete deficiency of normal EPG5 protein (∼280 kDa) suggesting the reduced stability of dysfunctional protein. Similar results were also reported in hESC (Vidyawan et al., 2025) and LCL (Piano Mortari et al., 2018) model with EPG5 haploinsufficiency. An impaired mitochondrial bioenergetic function and morphology was found in all patient fibroblasts. The mitochondrial respiratory defect was associated with downregulation of OXPHOS subunit expression despite an increase in mtDNA copy number. Gene enrichment analysis showed upregulation of PPARGC1A transcription factor encoding PGC1α, a master regulator of mitochondrial biogenesis (Supplementary Table S2), indicating an adaptive reaction that could compensate in part for impaired mitophagy in patient fibroblasts. Mitophagy defects may also impact downstream mitochondrial quality control regulated by mitochondrial proteases involved in remodelling of IMM proteome and initiating mitochondrial unfolded protein response (Uoselis et al., 2023; Chakrabarty et al., 2024; Yamada et al., 2025).

Unexpectedly, we found downregulation of MICU1/2 in patient fibroblasts and MICU1/3 in EPG5-mutant neurons as the causative mechanism for [Ca^2+^]_m_ overload and ensuing mPTP opening. Pathogenic mutations in MICU1 itself are associated with a rare childhood disorder that include neurodevelopmental, neurodegenerative and neuromuscular phenotypes with some close parallels to *EPG5*-RD (Logan et al., 2014; Bhosale et al., 2017; Kohlschmidt et al., 2021). Patients with loss-of-function MICU1 mutations present with a progressive extrapyramidal neurodegenerative disorder (Logan et al., 2014), features that have a significant overlap with the clinical phenotype associated with *EPG5*-RD, including ataxia, dystonia, and tremor as well as basal ganglia abnormalities in magnetic resonance imaging. Similarly, microcephaly, myopathy, seizures and immunodeficiency also present in MICU1 deficiency as well as in EPG5 deficiency. Furthermore, [Ca^2+^]_m_ overload and mitochondrial dysfunction due to downregulation of NLCX and MCU remodelling has been implicated in the neuropathology and memory decline in Alzheimer’s disease (Jadiya et al., 2019). Although the mechanism of MCU remodelling in patient fibroblasts and EPG5 mutant neurons defined by downregulation of MICUs and EMRE protein levels has not been investigated here, it seems likely that stress-induced activity of quality control proteases in IMM may control the protein turnover of MCU regulatory subunits. Proteostasis stress driven by impaired mitophagy may lead to the processing of DELE1 by OMA1 protease, culminating in activation of the integrated stress response (ISR) (Uoselis et al., 2023; Yamada et al., 2025) as evident by the ISR signature from RNA-seq data in patient fibroblasts (Supplementary Table S2). It is possible that the activity of alternative IMM or matrix proteases such as YME1L1 (Tsai et al., 2022) or AFG3L2 (Konig et al., 2016) respectively may control the processing and degradation of MICUs and EMRE respectively during the ISR.

Widespread mPTP opening causes pathological collapse of mitochondrial bioenergetic function, ATP hydrolysis and depletion, mitochondrial swelling and rupture culminating in neuronal cell death as evident during ischemic stroke (Schinzel et al., 2005). In contrast, limited or selective opening of mPTP, a feature also evident during histamine-induced [Ca^2+^]_m_ overload in patient fibroblasts, may cause insufficient damage to lead to cell death, but enable the release of mtDNA into the cytosol, where it triggers the activation of the cGAS-STING pathway, driving an inflammatory IFN response. This protective inflammatory program could be constitutively driven by patient fibroblasts in a chronic low-grade manner that maintains proliferation and evades acute cell death. A recent study described a similar process termed minority mitochondrial outer membrane permeabilization (miMOMP) where MOMP occurring in a subset of mitochondria enables mtDNA efflux through BAK/BAX macropores and activates the cGAS-STING pathway to drive age-associated inflammation in mice (Ichim et al., 2015; Victorelli et al., 2023). Mice deficient in *Epg5* and other primary autophagy genes such as *Atg5* and *Atg7* exhibit elevated cytokine expression and cellular lung inflammation sufficient to inhibit influenza pathogenesis and evade cell death (Lu et al., 2016). Notably, these cytokine levels are also increased in patients with *EPG5*-RD (Piano Mortari et al., 2018) and show a hyperimmunity against influenza. Constitutive activation of STING-mediated antiviral signalling arising as a result of defective mitophagy and mtDNA release may provide the explanation for the lack of influenza cases in patients with *EPG5*-RD. Consistent with these reports, we also found upregulated expression of pro-survival NF-κB target genes such as cFLIP and cIAP (Supplementary Table S2) in patient RNA-seq datasets in addition to the type I/III IFN signature genes.

The pathogenic Q336R EPG5 mutation in iPSC-derived neurons impaired mitochondrial respiration and reduced CI assembly similar to patient fibroblasts. The exposure to glutamate at concentrations that are innocuous for isogenic neurons caused a rapid collapse of bioenergetic capacity, disruption of [Ca^2+^]_c_ homeostasis and excitotoxic death in Q336R neurons. Glutamate-induced excitotoxicity in Q336R neurons is attributed to a lower mitochondrial Ca^2+^ uptake threshold and [Ca^2+^]_m_ overload which combined with increased ROS production leads to increased susceptibility to mPTP opening caused by downregulation of MICU1/3 holocomplex and a constitutively active MCU complex. Thus, an increased bioenergetic demand during neuronal metabolic workload may amplify the compromised mitochondrial function due to [Ca^2+^]_m_ overload and trigger a rapid pathological cascade culminating in cell death. Our data demonstrate that an impaired mitochondrial metabolism and Ca^2+^ signalling sensitises neurons to excitotoxic injury which may play an important role in neurodegeneration in *EPG5*-RD and may also contribute to epileptogenesis observed in around two-thirds of patients (Deneubourg et al., 2025). Loss of AFG3L2 activity, a mitochondrial m-AAA protease associated with spinocerebellar ataxia and loss of Purkinje neuron, results in impaired processing of EMRE and assembly of MCU complex enabling [Ca^2+^]_m_ overload, mPTP opening and neuronal death in mice (Konig et al., 2016). Moreover, neuronal loss of MICU1 increased [Ca^2+^]_m_ overload-induced excitotoxicity and caused progressive degeneration of motor neurons in MICU1-KO mice (Singh et al., 2022). These findings further highlight the impaired mitochondrial Ca^2+^ homeostasis as a key driver of excitotoxic neuronal cell death in early-onset neurodegenerative diseases.

Prevention of [Ca^2+^]_m_ overload-induced mPTP opening by JP1-138 effectively increased mitochondrial Ca^2+^ buffering capacity and partially rescued mitochondrial bioenergetic function in both patient fibroblasts and Q336R neurons. Prolonged treatment with JP1-138 also suppressed mtDNA release and downstream activation of the cGAS-STING pathway and IFN response in patient fibroblasts in comparison to CsA, the prototypical inhibitor of mPTP with off-target effects (Briston et al., 2019). A significant but partial improvement of mitochondrial bioenergetic function by JP1-138 underscores the pathophysiological impact of impaired mitophagy in regulating the neuronal energy homeostasis. Further investigations based on the pharmacological modulation of impaired mitophagy are needed to examine the role of mitophagy in mitigating inflammation and restoring mitochondrial function in EPG5-deficient cells. Inhibition of STING using H-151 equally supressed STING-dependent IFN response, although it failed to rescue mitochondrial dysfunction. Several studies have demonstrated that genetic ablation or pharmacological inhibition of STING protects against inflammation-mediated neuronal loss in mouse models of PD, ALS and lysosomal storage diseases (Sliter et al., 2018; Yu et al., 2020; Wang et al., 2024). Our findings suggest that the pharmacological effects of cGAS/STING inhibitors could be dependent on the mechanism mediating the pathogenesis which, in patient fibroblasts with EPG5 mutations, is a consequence of impaired mitochondrial homeostasis. Submicromolar concentrations of JP1-138 were highly effective at maintaining ΔΨ_m_ even after [Ca^2+^]_m_ overload in both patient fibroblasts and Q336R neurons. Our group recently reported improved mitochondrial activity and preclinical safety of JP1-138 compound with 20-fold greater brain concentration following a single dose in mice (Pingitore et al., 2024), highlighting its therapeutic potential in targeting Ca^2+^ mishandling and excitotoxicity in *EPG5*-RD and other neurodegenerative diseases.

In summary, this study expands our understanding of how impaired mitochondrial Ca^2+^ signalling and dysfunction can contribute to the presentation and progression of *EPG5*-RD by identifying a cascade of events starting from bioenergetic deficiency, unregulated mitochondrial Ca^2+^ uptake and Ca^2+^ overload to mPTP opening that culminates in low-grade mtDNA-driven chronic inflammation in patient fibroblasts and excitotoxicity and cell death in neurons. This progressive chain of events may account for the clinically variable and multisystem pathophysiology of *EPG5*-RD but also help to identify multiple sites for pharmacological intervention as potential novel therapeutic targets. These findings will together serve to improve our understanding of the progressive neurodegeneration in patients with *EPG5*-related disorders and further aid patients and families in genetic counselling.

---

**Figure S1:**
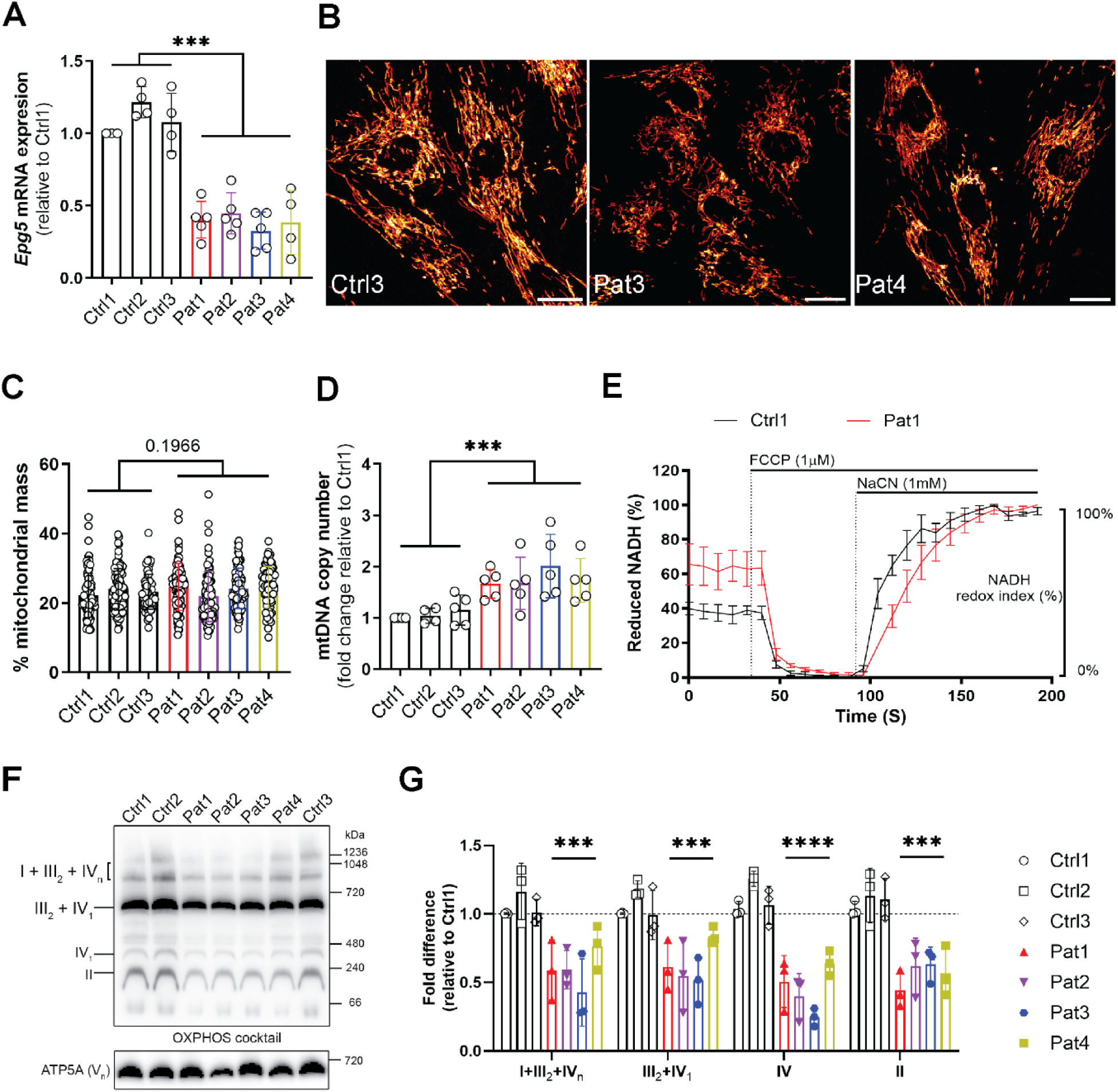
Mitochondrial dysfunction and increased mtDNA copy number in EPG5-deficient fibroblasts. **(A)** *EPG5* mRNA expression levels in patient fibroblasts determined with qRT-PCR on cDNA transcript of isolated mRNA and normalized to the levels measured in control, (n_exp_ = 4, \*\*\**p=* 0.0007). **(B)** Representative confocal image of TMRM labelled cells showing ΔΨ_m_ in control 3, patient 3 and patient 4 fibroblasts. Scale bars: 20 μm. **(C)** Quantification of mitochondrial volume occupancy calculated from total cytosolic volume (Calcein AM intensity value) and represented as percentage mitochondrial mass, (n_exp_ = 4, n_cells_ analysed for each control and patient = 95-110). **(D)** Quantification of mtDNA copy number measured by qPCR using a mtDNA-specific primer pair, from total genomic DNA isolated from control and patient fibroblasts, (n_exp_ = 4-5, \*\*\**p=* 0.0008). **(E)** Representative trace of quantified (mean ± SD) NADH autofluorescence in control and patient fibroblasts. Following the acquisition of baseline autofluorescence, application of 2.5 μM FCCP maximises the rate of respiration and oxidises all the mitochondrial NADH resulting in the lowest fluorescence signal (this level is considered as the minimum = 0% for NADH). Application of 1 mM NaCN blocks respiration and prevents NADH oxidation. This allows the NADH pool to be regenerated and the highest fluorescence signal is obtained (this level is considered as the maximum = 100% for NADH). The NADH redox index can be calculated using the obtained traces, (n_exp_ = 3). **(F)** Immunoblot image of native respiratory chain protein expression and supercomplex assembly from isolated mitochondria of control and patient fibroblasts analysed using blue native gel electrophoresis (BNGE). Supercomplex V_n_ was detected using ATP5A and used as a loading control. **(G)** Protein expression levels normalized and plotted as fold difference relative to control 1, (n_exp_ = 3, \*\*\**p=* 0.0005, 0.006 and 0.0001, \*\*\*\**p* <0.0001). Data (A, C, D, E, F, and H) are expressed as mean ± SD and individual data points from independent experiments are shown in each plot. Statistical analysis was carried out using two-way ANOVA followed by posthoc Tukey’s test (non-significant *p* values are denoted with numeric values).

**Figure S2:**
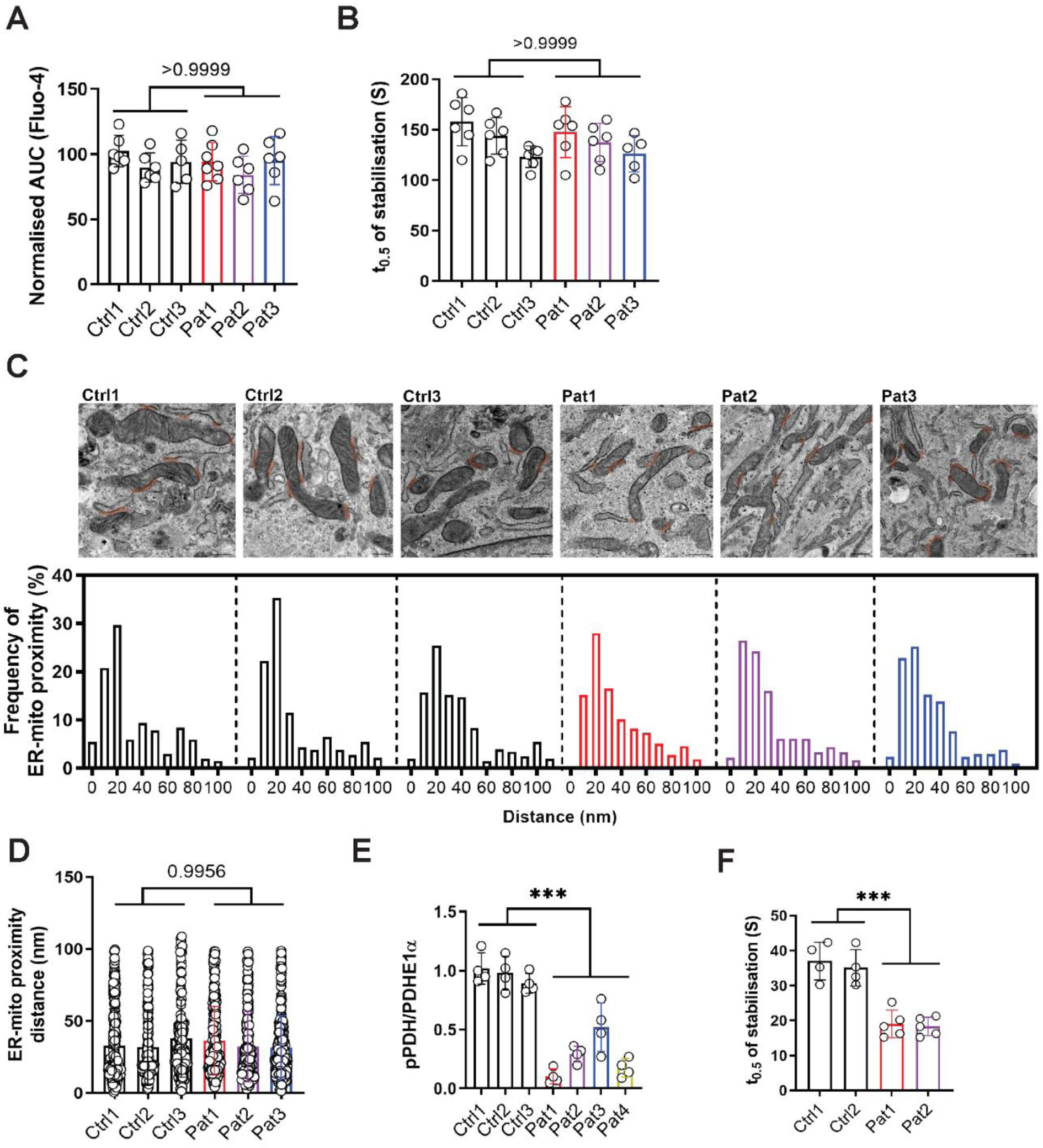
Both the ER Ca^2+^ stores and ER-mitochondria contact sites distribution are unaltered in control and patient-derived fibroblasts. **(A)** Quantification of normalised areas under the curve (AUC) of the Fluo-4 AM traces in response to 10 μM histamine, representing total ER Ca^2+^ released over time, calculated from the mean traces in Fig. 2B. **(B)** Measurement of time when 50% of released ER Ca^2+^ was cleared from the cytosol, calculated from the mean traces in Fig. 2B. **(C)** Representative TEM images of control and patient fibroblasts, with red segmented regions between ER and mitochondria marking ER-mito contact sites. Scale bars: 0.5 μm. Histograms of ER-mito contact widths distribution across control and patient fibroblasts, calculated from the segmented regions, (n_exp_ = 3, n_rep_ analysed for eachcontrol and patients =181-220). **(D)** Quantification of ER-mito contact widths from dataset in C. **(E)** The ratio of the pPDH (PDH-E1α pS293) and total PDH (PDH-E1α) band intensities normalised to the average control 1 ratio in Fig. 2J, (n_exp_ = 4, \*\*\**p=* 0.0004). **(F)** Time when [Ca^2+^]_m_ uptake was at the half maxima after the first 5 μM CaCl_2_ bolus, calculated from the inset in Figure 2K (\*\*\**p=* 0.0001). Data (A, B, D, E and F) are expressed as mean ± SD and individual data points from independent experiments are shown in each plot. Statistical analysis was carried out using two-way ANOVA followed by posthoc Tukey’s test (non-significant *p* values are denoted with numeric values).

**Figure S3:**
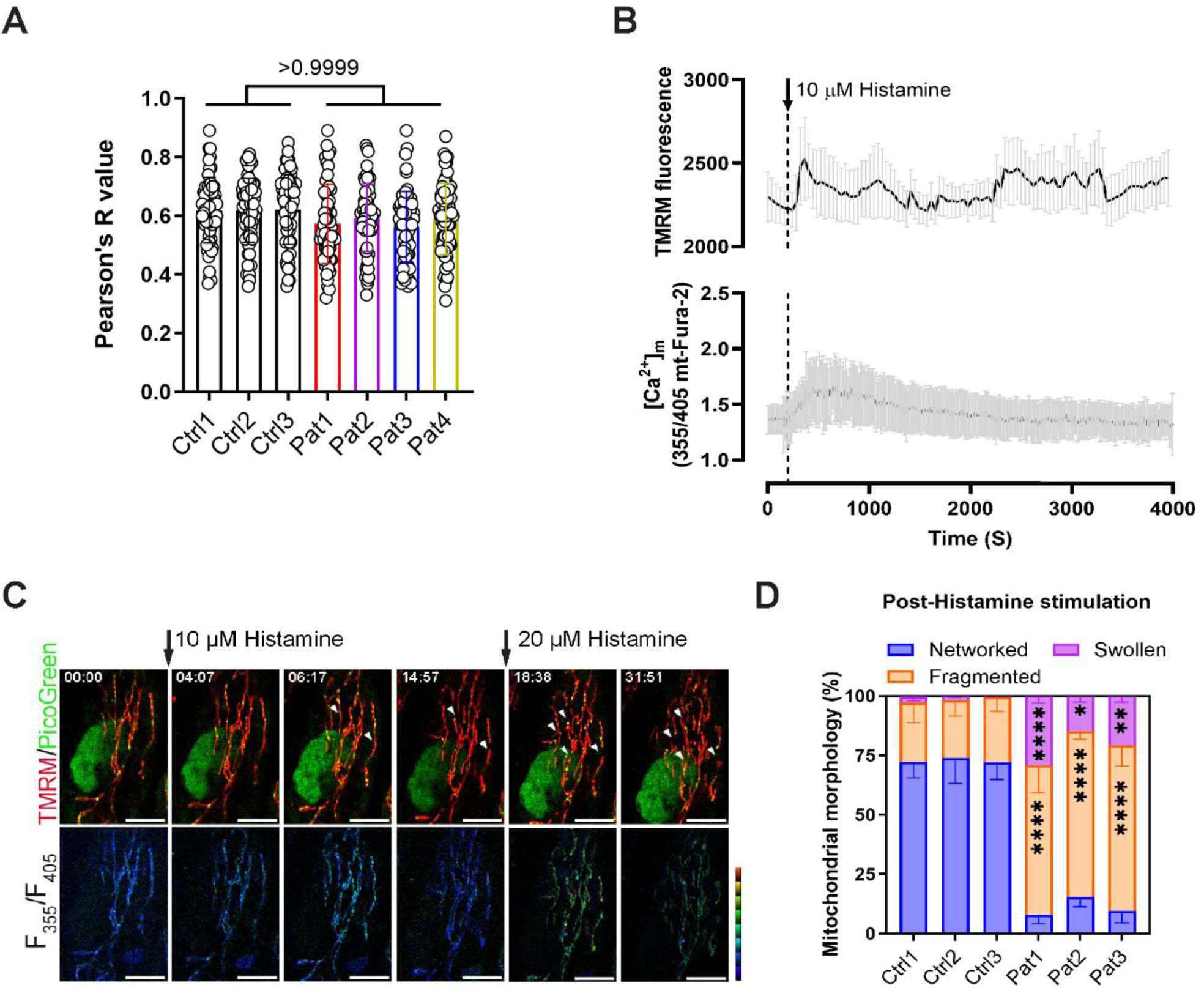
Physiological stimulation with histamine does not induces mitochondrial depolarisation or swelling in control fibroblasts. **(A)** Pearson’s R (colocalization) value of TOM20 (red) and citrate synthase (blue) labelled mitochondria from confocal images of control and patient fibroblasts represented in Fig. 3A, (n_exp_ = 4, n_cells_ analysed for each control and patient = 55-66). **(B)** Mean traces of TMRM fluorescence intensity and [Ca^2+^]_m_ change after 10 μM histamine challenge in control 1 fibroblasts co-labelled with TMRM, mito-Fura-2 AM and PicoGreen, (n_exp_ = 3). **(C)** Snapshots from time-lapse confocal imaging of control 1 fibroblasts co-labelled TMRM (red) and PicoGreen (green), upper panels and mito-Fura-2 AM, ratiometric lower panels, quantified in B. Elapsed time after 10 μM histamine challenge is indicated (Video S1). White arrowheads denote mitochondrial fragmentation events after 10 μM and 20 μM histamine challenge. Scale bars, 5 μm. **(D)** Quantitative morphometric analysis of TMRM labelled mitochondria in control and patient cells after stimulation with 10 μM histamine. Mitochondrial pool classified into networked, fragmented and swollen mitochondria are represented as a percentage of the total mitochondrial population, (n_exp_ = 3, n_cells_ analysed for each control and patients = 18-28, \**p=* 0.0313, \*\**p=* 0.0050, \*\*\*\**p* <0.0001). Data (A, B and D) are expressed as mean ± SD and individual data points from independent experiments are shown in each plot. Statistical analysis was carried out using one/two-way ANOVA followed by posthoc Tukey’s test (non-significant *p* values are denoted with numeric values).

**Figure S4:**
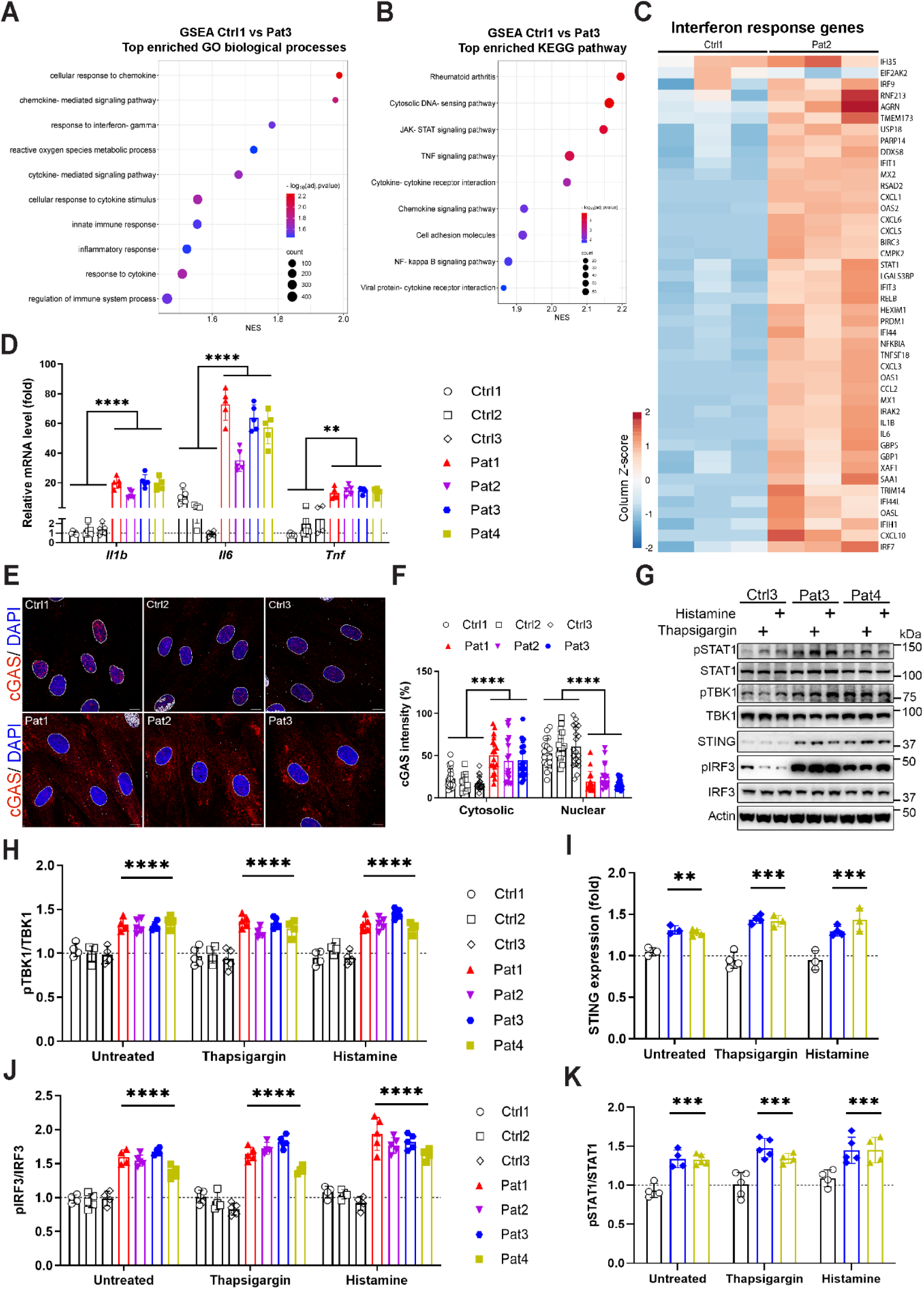
Additional characterisation of cGAS-STING activation and ISGs induction in patient-derived fibroblasts bearing EPG5 mutations. **(A-B)** Pathway analysis of RNA-seq data from patient 3 compared to control 1. Dot plots and GSEA performed using two types of gene set: the Gene Ontology (GO) biological process and KEGG pathway. NES, normalized enrichment score. **(C)** Heatmap of RNA-seq data displaying the top 50 upregulated differentially expressed type I/III IFN genes (DEGs) in control 1 and patient 3 fibroblasts, (n_exp_ = 3). **(D)** qRT-PCR analysis of pro-inflammatory ISGs expression in all control and patient fibroblasts, (n_exp_ = 3, \*\**p=* 0.0029, \*\*\*\**p* <0.0001). **(E)** Representative confocal images of control and patient fibroblasts immunolabelled with cGAS (red) antibody and stained with DAP1 to quantify cytoplasmic and nuclear localization of cGAS. **(F)** Quantitative analysis of relative cGAS fluorescence intensity from cytoplasmic and nuclear area to determine cytosolic cGAS translocation in control and patient fibroblasts, (n_exp_ = 3, n_cells_ analysed for each control and patient = 15-20, \*\*\*\**p* <0.0001). **(G)** Immunoblot images of proteins involved in cGAS-STING signalling and ISGs induction from whole cell lysates of control and patient fibroblasts treated with 10 μM histamine and 1 μM thapsigargin for 24 h. Actin was used as a loading control. **(H)** The ratio of pTBK1 (pSer172) and total TBK1 band intensities normalised to the control 1 ratio, (n_exp_ = 4, \*\*\*\**p* <0.0001). **(I)** Protein expression levels of STING normalized and plotted as fold difference relative to control, (n_exp_ = 4, \*\**p=* 0.0096, \*\*\**p=* 0.0003, 0.0006). **(J-K)** The ratio of pIRF3 (pSer396) and total IRF3 and pSTAT1 (p Tyr701) and total STAT1 band intensities normalised to the control 1 ratio, (n_exp_ = 4, \*\*\*\**p* <0.0001, \*\*\**p=* 0.0004, 0.0002). Data (D, F, H, I, J and K) are expressed as mean ± SD and individual data points from independent experiments are shown in each plot. Statistical analysis was carried out using two-way ANOVA followed by posthoc Tukey’s test.

**Figure S5:**
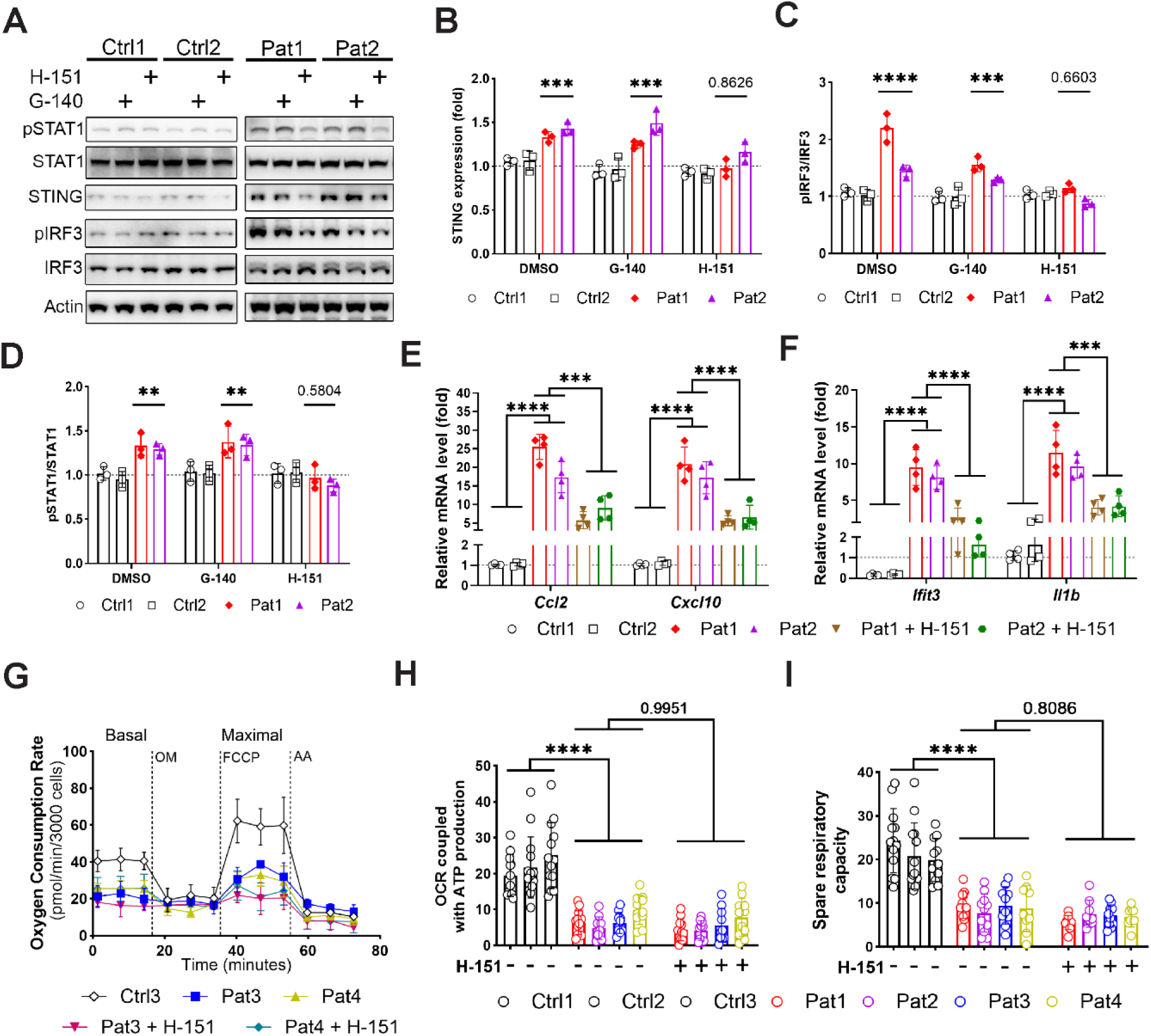
Inhibition of STING activation by H-151 attenuates the STING-dependent interferon response but does not improve mitochondrial function in patient-derived fibroblast. **(A)** Immunoblot images of the cGAS-STING signalling proteins from whole cell lysates of control and patient fibroblasts treated with either STING inhibitor, H-151 (1 μM) or cGAS inhibitor, G-140 (100 nM) for 24 h. Actin was used as a loading control. **(B)** Protein expression levels of STING normalized and plotted as fold difference relative to control, (n_exp_ = 3, \*\*\**p=* 0.0008). **(C-D)** The ratio of pIRF3 (pSer396) and total IRF3 and pSTAT1 (p Tyr701) and total STAT1 band intensities normalised to the control 1 ratio, (n_exp_ = 3, \*\**p=* 0.0034, \*\*\**p=* 0.0002, \*\*\*\**p* <0.0001). **(E-F)** qRT-PCR analysis of ISGs expression in the control and patient fibroblasts either untreated or treated with H-151 (1 μM) for 24 h (n_exp_ = 3, \*\*\**p=* 0.0006, \*\*\*\**p* <0.0001). **(G)** Normalised OCR traces from controls and patient fibroblasts either untreated or treated with H-151 for longer durations (0.5 μM, 3 days) (n_exp_ = 3, n_rep_ = 4). **(H-I)** Normalised ATP-linked respiration and spare respiratory capacity of all the controls and patient fibroblasts measured from traces in G and Fig. 5C (n_exp_ = 3, n_rep_ = 4, \*\*\*\**p* <0.0001). Data (B, C, D, E, F, G, H and I) are expressed as mean ± SD and individual data points from independent experiments are shown in each plot. Statistical analysis was carried out using two-way ANOVA followed by posthoc Tukey’s test (non-significant *p* values are denoted with numeric values).

**Figure S6:**
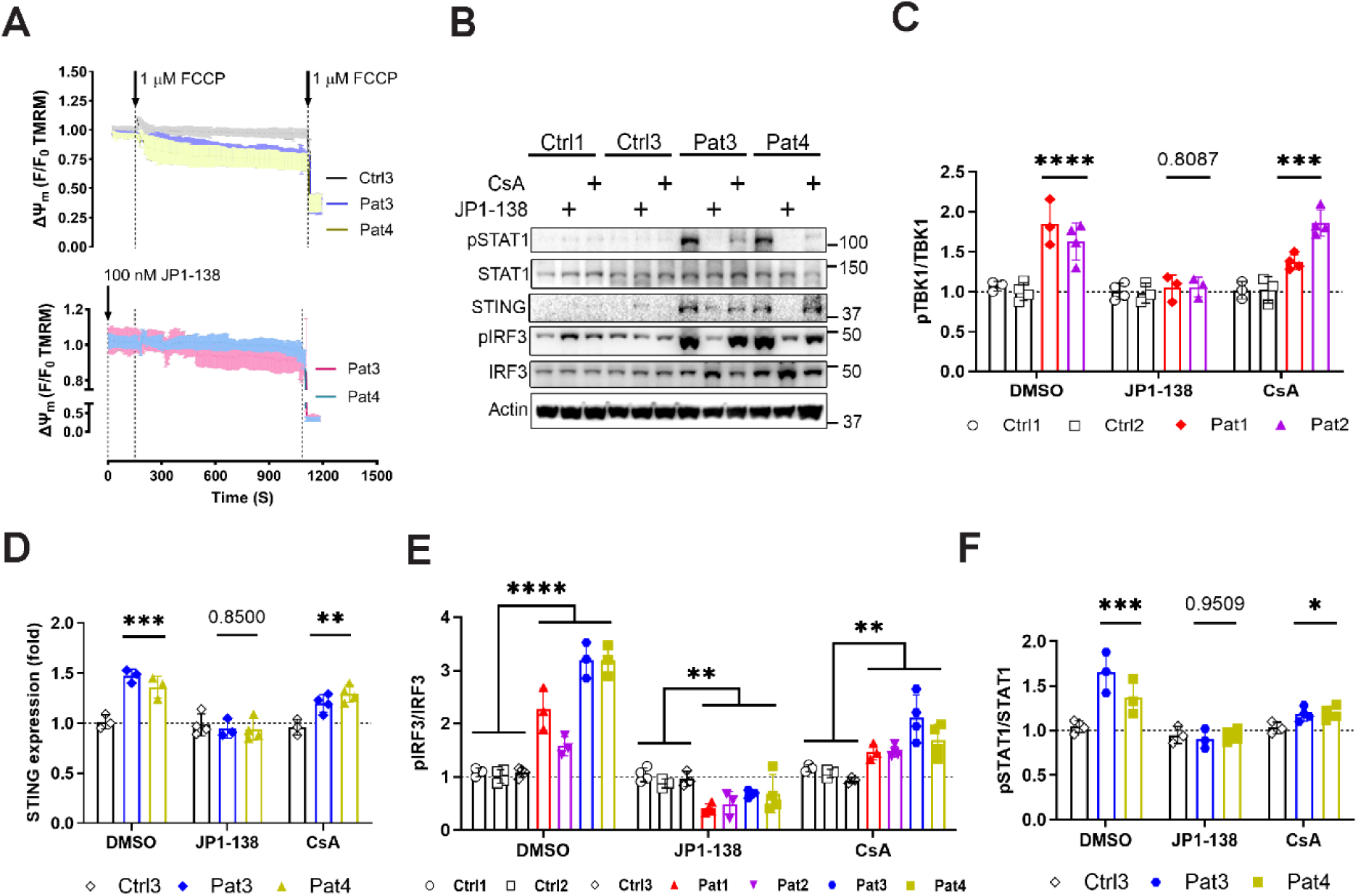
Additional characterisation of the effect of JP1-138 treatment on the STING-dependent interferon response in patient-derived fibroblasts. **(A)** Traces showing mean change ± SD in ΔΨ_m_ in response to 10 μM histamine challenge and FCCP-induced depolarization in the absence (upper panel) and presence of JP1-138 (100 nM) (lower panel) in control and patient fibroblasts, (n_exp_ = 4). **(B)** Immunoblot images of proteins involved in the cGAS-STING cascade from whole cell lysates of control and patient fibroblasts treated with either CsA (1 μM) or JP1-138 (100 nM) for 3 days. Actin was used as loading a control. **(C)** The ratio of pTBK1 (pSer172) and total TBK1 band intensities normalised to the control 1 ratio in Fig. 5G, (n_exp_ = 4, \*\*\**p=* 0.0001, \*\*\*\**p* <0.0001). **(D-F)** Relative expression levels of STING, ratio of pIRF3 (pSer396) and total IRF3 and pSTAT1 (p Tyr701) and total STAT1 band intensities normalized and plotted as fold difference relative to control, (n_exp_ = 3, \**p=* 0.0395, \*\**p=* 0.0020, \*\*\**p=* 0.0002, \*\*\*\**p* <0.0001). Data (A, C, D, E and F) are expressed as mean ± SD and individual data points from independent experiments are shown in each plot. Statistical analysis was carried out using two-way ANOVA followed by posthoc Tukey’s test (non-significant p values are denoted with numeric values, *p < 0.05, **p < 0.01, ***p < 0.001, ****p < 0.0001).

**Figure S7:**
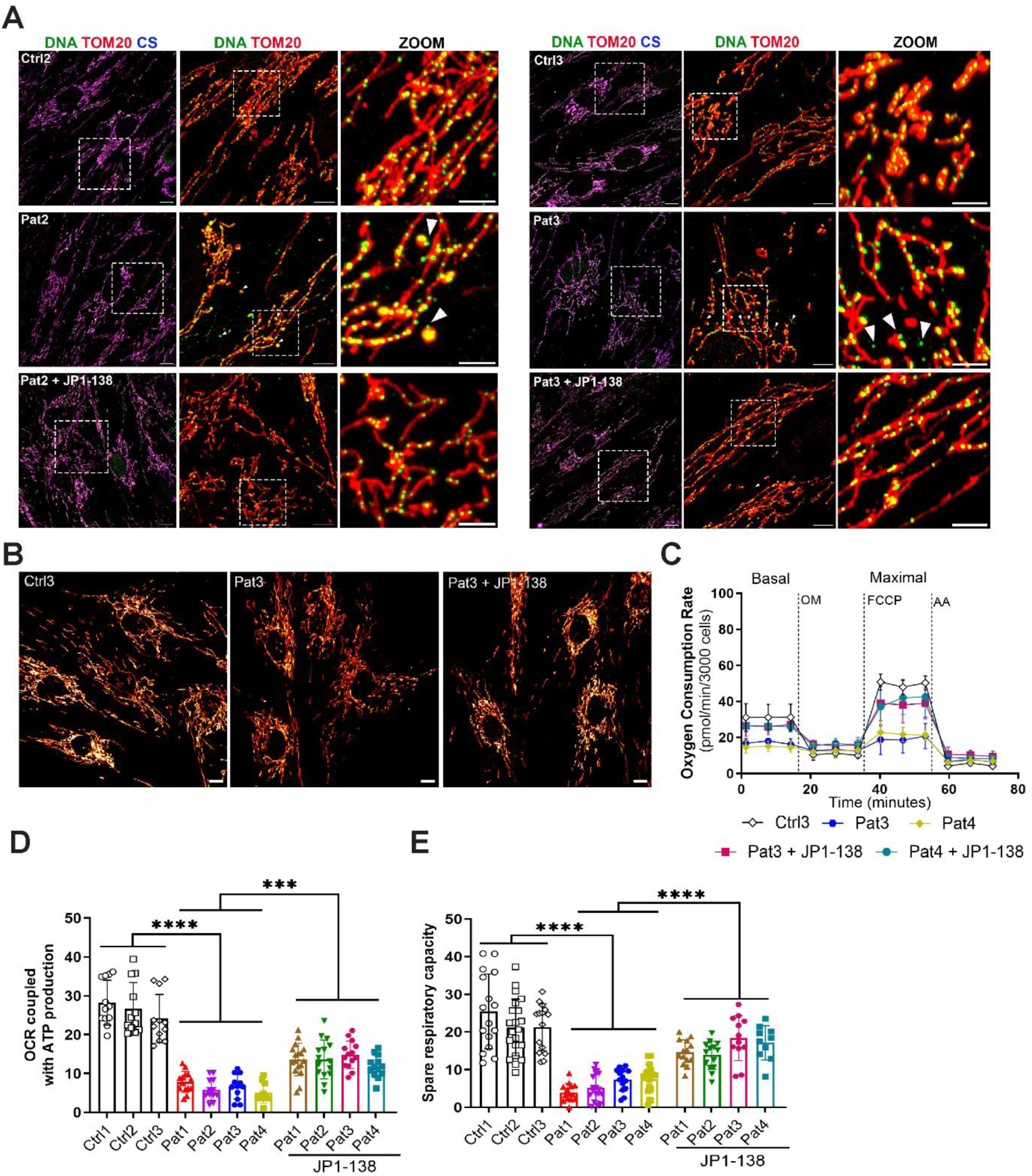
Additional characterisation of the effect of JP1-138 treatment on mitochondrial respiration and cytosolic mtDNA release in patient-derived fibroblast. **(A)** Representative Airyscan images of control and patient fibroblasts treated with either JP1-138 (100 nM) or vehicle (DMSO) for 3 days and immunolabelled with DNA (green), TOM20 (red) and citrate synthetase (blue). The magnified images on the right show areas in which DNA (green) does not co-localize with TOM20 (red) in patient cells marked by white arrow heads which is significantly reduced in JP1-138-treated patient fibroblasts. Overview scale bars, 10 μm and inset scale bars, 5 μm and 2.5 μm. **(B)** Representative confocal image of TMRM labelled control and patient fibroblasts treated with either JP1-138 (100 nM) or vehicle (DMSO) for 3 days. Scale bars: 20 μm. Normalised OCR traces from controls and patient fibroblasts treated with either JP1-138 (100 nM) or vehicle (DMSO) for 3 days (n_exp_ = 3, n_rep_ = 4). **(D-E)** Normalised ATP-linked respiration and spare reserve capacity of controls and patient fibroblasts calculated from traces in C and Fig. 6H (n_exp_ = 3, n_rep_ = 4, \*\*\*\**p* <0.0001). Data (C, D and E) are expressed as mean ± SD and individual data points from independent experiments are shown in each plot. Statistical analysis was carried out using two-way ANOVA followed by posthoc Tukey’s test.

**Figure S8:**
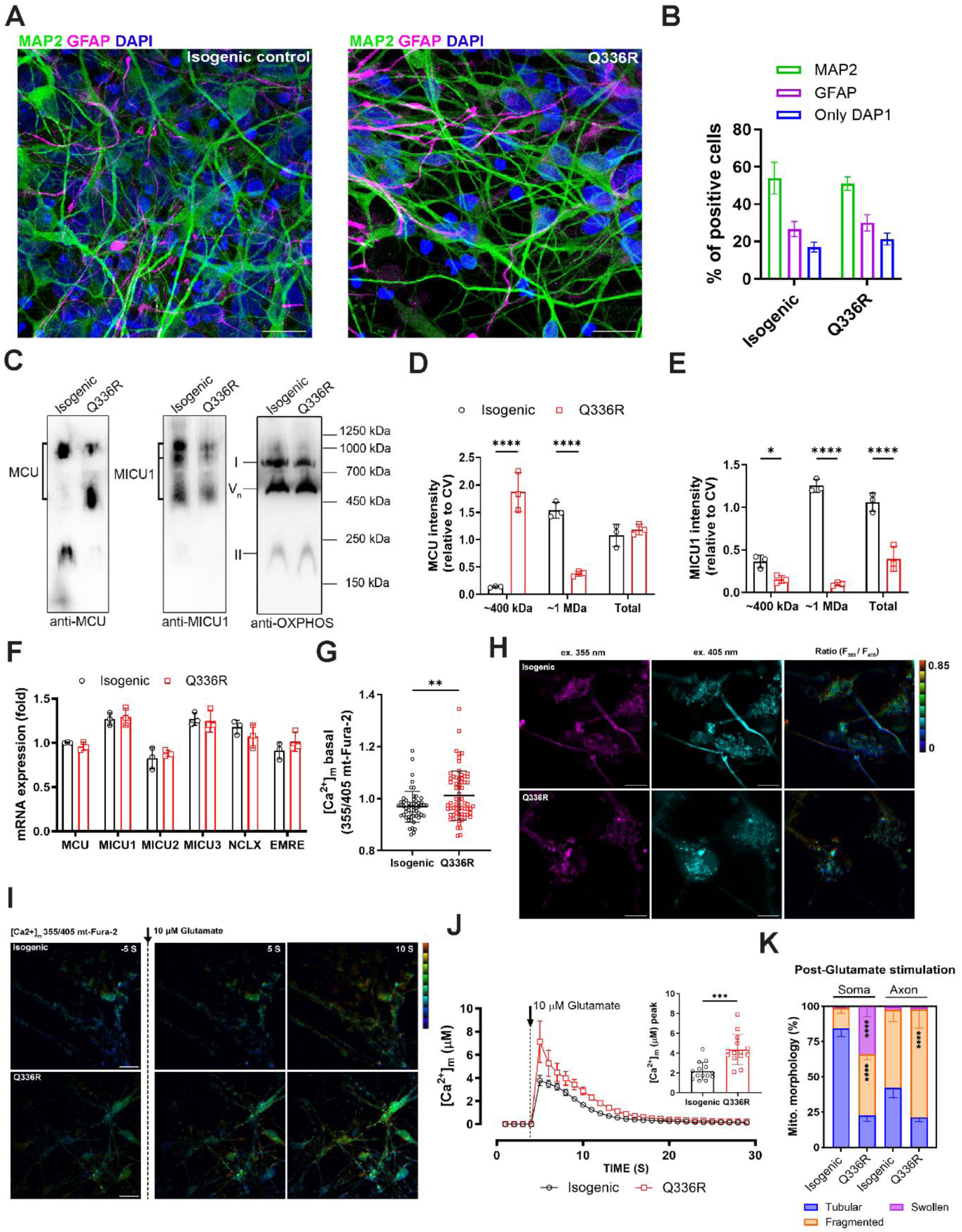
Altered mitochondrial Ca^2+^ homeostasis in Q336R neurons. **(A)** The neural cells were differentiated from NPCs and immunostained with the neuronal marker, MAP2 and the glial cell marker, GFAP. Scale bars, 20 μm. **(B)** The MAP2 (green) expression and GFAP (magenta) fluorescence levels were used to determine the percentage of targeted cells, while DAPI (blue) staining was used to determine the total number of cells in the field of view, (n_exp_ = 3). **(C)** Immunoblot image of native MCU complex and respiratory chain protein expression and supercomplex assembly identified using indicated antibodies from isolated mitochondria of isogenic and Q336R neurons and analysed using BNGE. **(D-E)** Quantitative analysis of high and low molecular weight MCU and MICU1 containing complexes analysed using BNGE in C, (n_exp_ = 3, \**p=* 0.0258, \*\*\*\**p* <0.0001). **(F)** mRNA levels of genes involved mitochondrial Ca^2+^ signalling using qPCR on cDNA transcript of isolated mRNA and normalized to the levels measured in isogenic control, (n_exp_ = 3). **(G)** Quantification of resting [Ca^2+^]_m_ in isogenic and Q336 neurons, calculated from calculated from the traces in Fig. 8H and I before glutamate stimulation, (\*\**p=* 0.0084). **(H)** Representative mito-Fura-2 ratiometric images for isogenic and Q336R neurons (the image excited at 355 nm divided by that excited at 405 nm) showing a higher steady-state [Ca^2+^]_m_ as well swollen morphology of mitochondrial in Q336R neurons. Scale bars, 10 μm. **(I)** Representative mito-Fura-2 ratiometric images for isogenic and Q336R neurons obtained at the start of the experiment (t = −5 s) and at 5 s and 10 s after exposure to glutamate, showing the respective rate of increase in [Ca^2+^]_m_ in isogenic control and Q336R neurons. Scale bars, 50 μm. **(J)** Mean traces for [Ca^2+^]_m_ uptake measured in isogenic and Q336R neurons using a mitochondria-target aequorin plate reader assay in response to 10 μM glutamate. Inset plot shows maximum [Ca^2+^]_m_ induced by 10 μM glutamate (n_exp_ = 3, n_rep_ = 5, \*\*\**p=* 0.0006). **(K)** Quantitative morphometric analysis of mitochondrial population in somas and axons of isogenic and Q336R neurons treated with either 10 μM glutamate or vehicle (PBS) for 12 h. All immunofluorescence images including representative images shown in Fig. 7M were classified into networked, fragmented and swollen mitochondria represented as a percentage of the total mitochondrial population, (n_exp_ = 3, n_cells_ analysed for isogenic and Q336R = 32-45, \*\*\*\**p* <0.0001). Data (B, D, E, F, G, J and K) are expressed as mean ± SD and individual data points from independent experiments are shown in each plot. Statistical analysis was carried out either using two-tailed unpaired Student’s t-test or one-way ANOVA followed by posthoc Tukey’s test.

**Figure S9:**
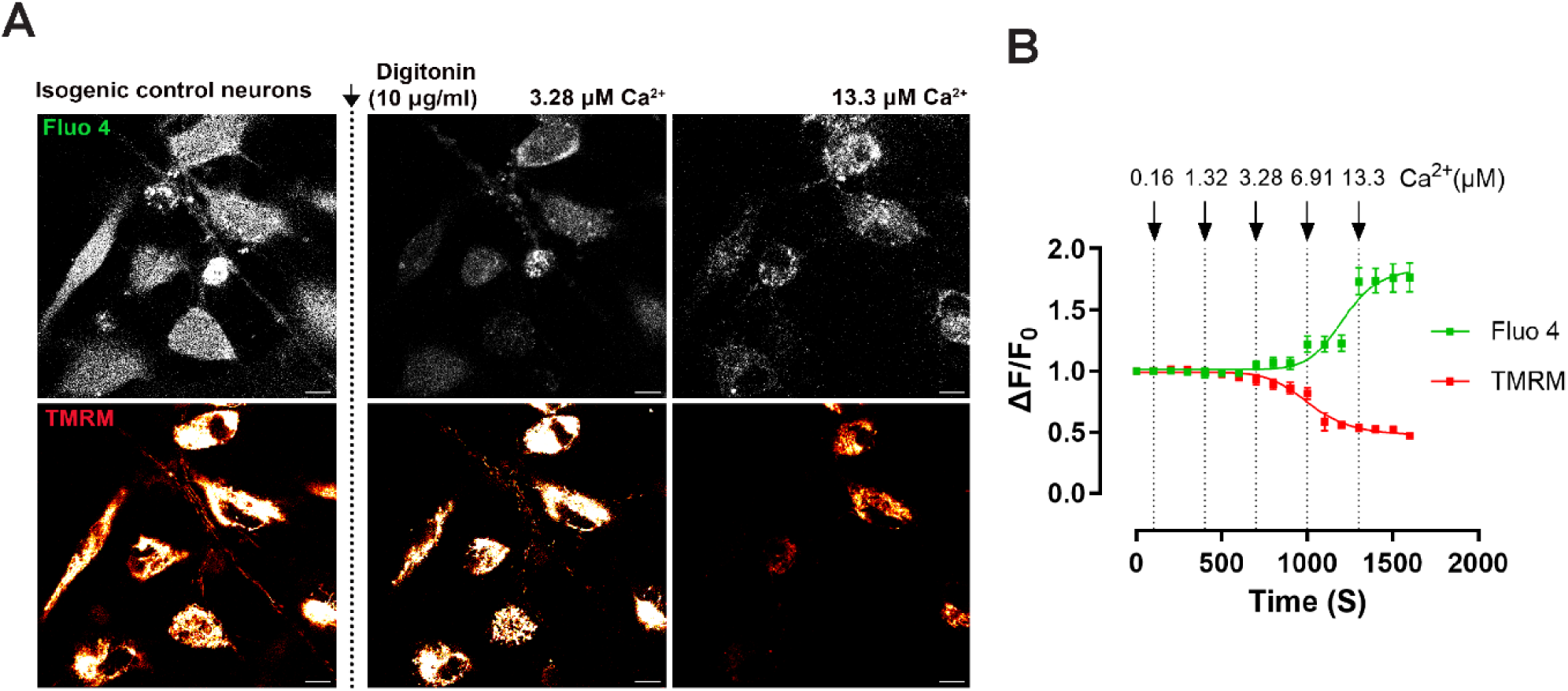
Mitochondrial Ca^2+^ buffering capacity determined by simultaneous measurement of [Ca^2+^]_m_ and ΔΨ_m_ in Q336R neurons. **(A)** Representative confocal images of permeabilised isogenic control neurons show changes in fluorescence of TMRM (red) and Fluo-4 AM (grey) in response to increasing concentrations of calcium (3.28 and 13.3 μM). Before digitonin permeabilization, Fluo-4 intensity shows cytosolic localization of the dye along with TMRM signal localizing to tubular mitochondria both in somas and axons of isogenic control neurons. The punctate staining pattern of Fluo 4 and TMRM following permeabilization confirms mitochondrial localisation of the dyes. **(B)** Traces showing mean change ± SEM in Fluo-4 (representing [Ca^2+^]_m_) and TMRM (representing ΔΨ_m_) fluorescence intensity in response to increasing concentrations of Ca^2+^ in the media. Quantified data from one representative control experiment depict the characteristic responses. The final free Ca^2+^ in the media is indicated for each respective addition.

## Video description

**Video S1: Time-lapse imaging of ΔΨ_m_, [Ca^2+^]_m_ and mtDNA dynamics in response to histamine in control 1 fibroblasts.** Control 1 fibroblasts co-labelled with TMRM, mito-Fura-2 and PicoGreen were imaged every 12.5 s to simultaneously monitor the change in ΔΨ_m_, [Ca^2+^]_m_ and mtDNA dynamics, respectively, in response to the successive application of 10 μM and 20 μM histamine. Movie playback 30fps. Scale bar, 10 μm.

**Video S2: Time-lapse imaging of ΔΨ_m_, [Ca^2+^]_m_ and mtDNA extrusion in response to 10 μM histamine in patient 1 fibroblasts.** Patient 1 fibroblasts co-labelled with mito-Fura-2, TMRM and PicoGreen were imaged every 12.5 s to simultaneously monitor the increase in [Ca^2+^]_m_, collapse of ΔΨ_m_, and mtDNA extrusion, respectively, in response to 10 μM histamine challenge. Movie playback 30fps. Scale bar, 10 μm.

## Materials and Methods

### Cell culture and treatments

Fibroblasts carrying specific EPG5 mutations (Table S1) were isolated from skin biopsies obtained from patients as part of the diagnostic process. Age and sex matched healthy control fibroblasts were obtained from the MRC Centre for Neuromuscular Disorders Biobank, London. All cell lines were cultured and maintained in Dulbecco’s modified Eagle’s medium (10566016, Gibco) supplemented with 10% fetal bovine serum (16140071, Gibco), and 1% Antibiotic-Antimycotic (15240096, Gibco) and incubated at 37°C with 5% CO_2_. Fibroblasts lines were maintained at maintained at sub-confluence (80%) and cultured between 3 to 10 passages. All cell cultures were routinely checked for mycoplasma using MycoAlert™ Mycoplasma Detection Kit (LT07-118, Lonza). For transfection, Human Dermal Fibroblasts Nucleofector Kit (VPI-1002, Lonza) was used according to the manufacturer’s instructions. Fibroblasts were treated with either freshly prepared histamine (CAY33828, Cambridge Bioscience) or a thapsigargin (586005, Merck), H-151 (S6652, Selleck) and G140 (S9945, Selleck) stock solutions made in DMSO, at indicated working concentrations for 24 h. Fibroblasts and neuronal cultures were preincubated with JP1-138 in imaging media for 10 minutes before live cell imaging. For long-term treatments with either JP1-138, CsA (1101, Tocris) or H-151, the culture medium was changed every two days.

### hiPSCs derivation and culture

The iPSC lines, GM27291 and GM28930 used in this study were generated by Coriell Institute. iPSC cell line GM27291 was derived from GM26636 fibroblasts (Patient 1) (Mitchell et al., 2022) and iPSC cell line GM28930 is the isogenic control for the patient-derived line GM27291. iPSCs were thawed, expanded, and maintained using hESC-qualified matrigel-coated plates (356277, Corning) with mTeSR1 iPSC medium (85850, STEMCELL Technologies) as per the commercially available protocol from STEMCELL Technologies. The iPSCs were passaged when they reached approximately 70%–80% confluency using 0.5 mM EDTA (S311-500, Fisher Scientific) prepared in PBS (10010023, Thermo Fisher Scientific).

### Differentiation of human iPSCs to neuronal networks

Cortical neuronal cultures were generated from human iPSCs according to (Shi et al., 2012). Neural maintenance media (NMM) comprised a 1:1 mixture of DMEM/F-12 GlutaMAX (10565018, Thermo Scientific) with Neurobasal media (12348-017, Thermo Scientific), supplemented with 2.5 µg/ml insulin (I9278, Sigma-Aldrich), 1 mM L-glutamine (25030-024Thermo Scientific), 50 µm nonessential amino acids (NEAA) (11140-050, Thermo Scientific), 50 µM β-mercaptoethanol (31350010, Gibco), 0.5x N-2 supplement (17502001, Thermo Scientific), 0.5x B-27 (17504001, Thermo Scientific), and 0.25% penicillin/streptomycin (15140122, Gibco). Neural induction media (NIM) consisted of NMM with 1 µM Dorsomorphin (3093, Tocris) and 10 µM SB431542 (1614, Tocris). In brief, iPSCs were cultured as above transferred at a seeding ratio of 2:1 to hESC-qualified Matrigel coated wells, in mTeSR1 supplemented with 10 µM ROCK inhibitor (1254/10, Bio-techne). Cells were incubated at 37°C for 24 h-48 h to reach 100% confluence. Cell layers were gently washed with PBS and medium replaced with Neural Induction Media (NIM), which was then replaced daily, until a dense neuro-epithelial sheet formed at around 8-12 days after neural induction. At this point, cells were passaged using 1 mg/ml dispase (17105041, Thermo Scientific) in NIM, and lifted as aggregates. These aggregates were washed and seeded at a ratio of 1:2 in fresh NIM, onto laminin (L2020-1MG, Sigma-Aldrich) wells. Cells were incubated overnight to allow attachment, and clumps that had not been attached were transferred to fresh laminin coated wells. Upon the formation of neural rosettes, culture medium was replaced with NMM supplemented with 20 ng/ml recombinant human FGF2 (100-18B, PeproTech), which was changed every 48 h. FGF2 was withdrawn from the culture medium after 4 days, and cells were expanded using dispase to lift neural rosettes. On day 25 (±1 day) after induction, cultures were dissociated into single cells using Accutase (A1110501, Thermo Scientific), and seeded in NMM at a ratio of 1:1 in fresh laminin coated wells. From this point onwards, cells were expanded as single cell suspensions with Accutase, and media was replaced at least every 48 h. At ∼ day 36 of neural induction, iPSC-NPCs (neural progenitor cells) were seeded in poly-L-lysine (P4707, Sigma) and laminin coated plates at a density of 5×10^4^ cells/cm^2^ for terminal neuronal differentiation, and NMM was replaced at least every 48 h. Neuronal cultures were matured until at least day 62.

For neuronal characterization, Neuronal cultures in glass-bottom 96-well plate (655892, SensoPlate^TM^) were fixed for 15 min at room temperature using a 4% paraformaldehyde solution, washed and permeabilized for 5 min with 0.1% Triton X-100 (AC215682500, Fisher Scientific) in PBS. Fixed cells were blocked with 3% BSA (A6003, Sigma) in PBS to block the nonspecific bindings for 60 min. Cells were then incubated overnight at 4°C with the GFAP and MAP2 primary antibodies prepared in blocking solution. The following day, the cells were washed with PBS and incubated with secondary antibodies, Alexa Fluor 488 or Alexa Fluor 568 based on the primary antibody host species, for 1 h. Then, the cells were washed with PBS and incubated with 4,6-diamidino-2-phenylindole (DAPI) solution (1:10,000, D1306, Thermo Scientific) for 5 min to detect all nuclei and imaged using confocal microscope as described below. Details of the antibodies used in this study can be found in Supplementary Table S3.

### RNA-sequencing

Fibroblasts for RNA-sequencing were seeded at a density of 1 × 10^6^ cells/dish and cultured in 10 cm dishes with regular media before harvesting on day two. RNA sequencing was performed at the UCL Genomics Core Facility using the Mag-Bind® Total RNA 96 Kit for RNA-sequencing polyA capture on the Illumina NextSeq 2000 P2 sequencer. Reads were mapped and analyzed by the SARTools R package. Differential expression analysis was carried out with the Bioconductor package DESeq2 (v.1.48.0) (Love et al., 2014). Genes were annotated using the org.Hs.eg.db package (Genome wide annotation for Human, v.3.21.0) and differentially expressed genes (DEG) with an adjusted p-value cut-off of 0.05 were identified as statistically significant. Gene ontology (GO) and Kyoto Encyclopedia of Genes and Genomes (KEGG) enrichment analysis were performed using the ClusterProfiler package (Yu et al., 2012). Results of DEG analysis were visualized with heatmaps plotted in ClustVis (Metsalu and Vilo, 2015).

### Transmission Electron Microscopy

Fibroblasts cultured on coverslips were fixed in EM fixative (2% glutaraldehyde + 2% paraformaldehyde in 0.1M sodium cacodylate) for 1 h followed by washings with 0.1M Cacodylate Buffer. The Coverslips were then fixed in 1% osmium tetroxide and 1% potassium ferricyanide in 0.1M sodium cacodylate, followed by sequential dehydration in ethanol. The coverslips were embedded in Epoxy Resin (Araldite CY212) Kit (Agar Scientific Ltd.) according to the standard protocol. Embedded coverslips were sectioned to 50 nm using an ultra-microtome fitted with a diamond knife, mounted onto TEM compatible copper grids. The grids were then stained with Lead Citrate for 3 min before proceeding for Imaging. Images were acquired using Jeol 1400 Transmission Electron Microscope and Gatan software. Images were taken at a magnification of 1200X (digital magnification). Interfaces between ER and mitochondria were segmented using DeepMIB software (Belevich and Jokitalo, 2021) analyzed using a custom ImageJ script: https://sites.imagej.net/MitoCare/.

### ROS measurements

The rate of intracellular ROS production in fibroblasts was measured using a superoxide indicator, dihydroethidium (DHE; D11347, Thermo Scientific) as described previously (Chung and Duchen, 2022). Following the measurements of fluorescence intensity, cells were stained with Hoechst 33342 (62249, Thermo Scientific) for 10 min to label and count the numbers of cell nuclei representing cell numbers in each well using an automated fluorescent image acquisition system (ImageXpress MicroXL). Subsequently, the fluorescence intensity was normalized based on the relative cell numbers obtained.

### SDS-PAGE and immunoblotting

Fibroblast cultures from 60 mm dishes were washed with PBS and collected by trypsinization (0.5% trypsin-EDTA; 15400054, Gibco). Similarly, mixed neuronal cultures were gently lifted and collected using Accutase. To prepare the samples for Western blot analysis, cell pellets were homogenized in 100–150 μl RIPA buffer (R0278, Sigma-Aldrich) containing 1x Halt Protease and Phosphatase Inhibitor Cocktail (78440, Thermo Scientific). Following homogenization, cell lysates were centrifuged at 16,000g at 4 °C for 30 min and the protein concentration in the supernatant was quantified using the Pierce BCA Assay Kit (23227, Thermo Scientific). Equivalent amounts of total protein (30 µg) samples in NuPAGE 4x LDS Sample Buffer (NP0007, Invitrogen) and 2% β-mercaptoethanol (63689, Sigma-Aldrich) were boiled at 95 °C for 10 min. Proteins were separated on either 12% Bolt Bis-Tris Plus (NW04127, Invitrogen) or 4–12% NuPAGE Bis-Tris polyacrylamide gels (NP0335, Invitrogen) immersed in MOPS running buffer (NP0001, Invitrogen) and transferred onto PVDF membranes (1620175, Bio-Rad). Membranes were then blocked in SuperBlock Blocking Buffer (37545, Thermo Scientific) for 1 h at room temperature, and probed overnight at 4°C using indicated primary antibodies. After incubation with appropriate secondary antibodies, protein bands were detected using a chemiluminescent reagent (Luminata Forte Western HRP substrate; WBLUF0100, Merck) and imaged using a ChemiDoc system (Bio-Rad). Membranes were further stripped using Restore Western Blot Stripping Buffer (21059, Thermo Scientific) and re-probed with additional primary antibodies. Quantification of the protein bands was performed using with ImageJ (NIH) and Image Lab software, v6.0.1 (Bio-Rad). Details of all the antibodies used in this study can be found in Table S3.

### Blue native gel electrophoresis (BNGE) and immunoblotting

Mitochondria from the fibroblasts were isolated using mitochondria isolation buffer 1 (MIB1; 225 mM Mannitol, 75 mM Sucrose, 5 mM HEPES, 1 mM EGTA and 1 mg/ml fatty acid free BSA) and MIB2 (same as MIB1 but without BSA) according to the method described previously (Singh and Duchen, 2022). Mitochondria isolation from the neuronal cultures were performed similarly, with the exception of homogenization step, performed using a Dounce tissue grinder tube. Protein concentration of the isolated mitochondria was quantified using the Pierce BCA Assay Kit (23227, Thermo Scientific) and an equivalent amounts of total protein (50 µg) was solubilized with digitonin followed by centrifugation at 20,000g for 20 min at 4°C. Digitonin-solubilized mitochondria were separated on 3–12% NativePAGE Bis-Tris gels (BN1001, Invitrogen) and electroblotted onto PVDF membrane (1620175, Bio-Rad) according to the manufacturer’s instructions. Membranes were then blocked and probed with indicated primary antibodies as described above. Quantification of the protein bands was performed using ImageJ. Details of all the antibodies used in this study can be found in Table S3.

### Quantitative reverse transcription PCR (RT-qPCR)

Total RNA was extracted from fibroblasts and neuronal culture using the RNeasy Plus Mini Kit (74104, Qiagen) according to the manufacturer’s instructions. Quality check and the quantification of isolated mRNA was done on the Nanodrop 2000c (ND-2000, Thermo Scientific). cDNA was synthesized from 1 μg of total RNA using SuperScript IV First-Strand Synthesis System Kit (18091050, Invitrogen) and quantitative PCRs was performed using SYBR Green JumpStart Taq ReadyMix (Sigma-Aldrich) on a CFX96 Real-Time PCR Detection System (Bio-Rad). Data were analysed using the comparative 2^−ΔΔCt^ method. Ct of the gene of interest was normalized to that of β-actin.

For the quantitative analysis of the relative mtDNA copy number, total genomic DNA was extracted from fibroblasts using the DNeasy Blood & Tissue Kit (69506, Qiagen) and the quantitative PCR was performed similarly with primers for the mtDNA tRNALeu (UUR) and for the nuclear B2M (β-2-microglobulin) to determine the relative mtDNA copy number of cells (Rooney et al., 2015). The following equation was used to determine the relative mitochondrial DNA content, 2 x 2^ΔCt^, where ΔCt is nuclear DNA Ct value subtracted by mtDNA Ct value. All primer pairs used can be found in Table S4.

### Mitochondrial oxygen consumption rate (OCR)

Measurements of mitochondrial respiration in fibroblasts and neurons were conducted with the Seahorse Bioscience XFe96 bioanalyzer using the Seahorse XF Cell Mito Stress Test Kit (103015-100, Agilent). Fibroblasts were seeded at a density of 1 × 10^4^ cells/well on XF96 cell culture microplates (102416-100, Agilent) and cultured for 1-2 days and/or treated for the indicated time. For neuronal cultures, iPSC-derived NPCs were seeded at a density of 5 × 10^4^ cells/well on XF96 cell culture microplates coated with poly-L-lysine and laminin for terminal neuronal differentiation and maturation using the method describe above. On the day of the experiment, the culture medium was replaced with Seahorse XF Base medium (103334-100, Agilent) supplemented with 1mM pyruvate (11360070, Gibco), 2 mM glutamine (25030081, Gibco) and 10mM glucose (A2494001, Gibco) and incubated in a CO_2_-free incubator for 30 min at 37 °C before loading into the Seahorse Analyser. After the measurement of basal respiration, the drugs oligomycin (5 μM), FCCP (1 μM, 2 μM), and rotenone/antimycin A (0.5 μM/ 0.5 μM) were added to each well in sequential order. Data was analysed using the XF Cell Mito Stress Test Report Generator. The OCR data from fibroblasts were normalized to the cell number obtained after counting the numbers of cell nuclei with ImageXpress MicroXL as described above. The OCR data from neurons were also normalized to the cell counts obtained after constructing a calibration curve and calculating the normalized cell number per well with CyQuant Direct Cell Proliferation Assay Kit (C35011, Invitrogen) and expressed as a percentage of the baseline measurement of untreated line.

### Mitochondrial membrane potential (ΔΨ_m_) in fibroblasts and neurons

The steady-state and time-lapse measurement of ΔΨ_m_ in fibroblast was carried out using tetramethylrhodamine methyl ester (TMRM) in the redistribution mode where a lower fluorescence intensity indicates reduced ΔΨ_m_ and vice versa. For the steady-state measurement of ΔΨ_m_, cells were seeded at a density of 1 × 10^5^ cells/dish on fluorodishes (FD35-100, WPI) and cultured for 1 day and/or treated for the indicated time. On the day of the experiment, cells were washed once with recording buffer 1 (RB1; 150mM NaCl, 4.25mM KCl, 4 mM NaHCO_3_, 1.25mM NaH_2_PO_4_, 2 mM CaCl_2_, 1.2 mM MgCl_2_, 10mM D-glucose, and 10mM HEPES at pH 7.4) and incubated with 25 nM TMRM (T668, Invitrogen) and 1 μM Calcien-AM (C3100MP, Invitrogen) in RB1 for 30 minutes at 37°C. Following incubation, cells were washed twice with RB1 and TMRM was added in the RB1 to avoid its depletion while imaging. z-stacks with 0.45 µm thickness with a pixel dwell time of 1.54 μs were acquired using a Zen Black software-controlled LSM 880 confocal microscope (Carl Zeiss) equipped with a plan-Apochromat 63x/1.4 oil DIC objective lens and an Ar (λ_ex_ = 488 nm, λ_em_ = 500–550 nm for Calcien-AM)and DPSS laser source (λ_ex_ = 561 nm, λ_em_ = 575–625 nm for TMRM) at 37 °C. Mean TMRM fluorescence intensity was quantified using the same threshold across all the samples and the percentage mitochondrial mass was calculated from the area of the binarized images of CalcienAM and TMRM using ImageJ.

For the time-lapse measurement of ΔΨ_m_, images were acquired from a single z-plane every 2 s interval with a pixel dwell time of 1.54 μs. After acquiring baseline TMRM images, 10 uM histamine was applied directly into the fluorodishes using a micropipette and subsequent images were obtained to monitor the effect of histamine on ΔΨ_m_. At the end of each experiment, 2.5 μM Carbonyl cyanide 4-(trifluoromethoxy) phenylhydrazone (FCCP; C2920, Sigma-Aldrich) was added as a positive control which depolarized the mitochondria. Time series were analysed using ImageJ by selecting and measuring the mean fluorescence intensity in the regions of interests (ROIs) in each field. Individual ΔΨ_m_ traces were normalized to their baseline intensity obtained before stimulation.

Measurement of ΔΨ_m_ in neurons was carried out using Rhodamine-123 in the dequench mode where dequenching or increase in Rhodamine-123 fluorescence indicates mitochondrial depolarization and loss of ΔΨ_m_. Neuronal cultures seeded in a glass-bottom 96-well plate (655892, SensoPlate) were labelled with 10 μg/ml Rhodamine-123 (R8004, Sigma-Aldrich) in BrainPhys Imaging Optimized Medium (05796, STEMCELL Technologies) at 37 °C and 5% CO_2_ for 20 min. After incubation, cells were washed thrice and imaged as above using a Plan-Neofluar 40x/1.30 oil objective and an Ar laser source (λ_ex_ = 488 nm, λ_em_ = 510–600 nm). For all experiments, the laser illumination intensity was kept to a minimum (max 0.5%) to avoid phototoxicity and photobleaching. After acquiring baseline Rhodamine-123 images, neurons were stimulated using 10 μM glutamate (G1626, Sigma-Aldrich) and subsequent images were acquired. At the end of each experiment, 1 μM FCCP was added to evaluate the Rhodamine-123 fluorescence intensity corresponding to the loss of ΔΨ_m_. Time series were analysed using ImageJ as described above.

### Mitochondria morphology analysis

Morphometric analysis of TMRM-labelled or TOM20-immunolabelled raw confocal images was performed using a 2D cell segmentation model, MitoSegNet (Fischer et al., 2020). Neuronal somas and axons were manually segmented in ImageJ using β-tubulin III staining for morphological distinction. Briefly, images were pre-processed to 8-bit format and mitochondria were first segmented using the MitoS basic toolbox (https://github.com/MitoSegNet). Segmentation masks were then run on MitoA analyzer tool for the quantification of the morphological features broadly categorized into shape descriptors such as area, eccentricity and perimeter and the network descriptors such as branch length, number of branches and curvature index. The shape descriptors values were then used to calculate the percentage of elongated, fragmented or swollen mitochondrial pool.

### Measurement of mitochondrial NAD(P)H

NAD(P)H autofluorescence imaging was carried out to investigate the mitochondrial redox state in fibroblasts. Cells were seeded at a density of 1 × 10^5^ cells/dish on fluorodishes and cultured for 1 day and/or treated for the indicated time. On the day of the experiment, cells were washed once with RB1 and imaged using the Carl Zeiss LSM 880 confocal microscope equipped with an ultraviolet (UV) laser (λ_ex_ = 355 nm, λ_em_ = 410–480 nm) and quartz Plan-Apochromat 63x /1.4 oil objective at 37 °C. Images were acquired from a single z-plane with the pinhole wide open to maximize signal and laser illumination intensity kept to a minimum (0.1-0.2%) to avoid phototoxicity. After acquiring baseline images, cells were first exposed to 2.5 μM FCCP to depolarize the mitochondria completely and achieve maximal respiration. The oxidation of the mitochondrial pool of NADH into non-fluorescent NAD^+^ led to the lowest fluorescence signal, which was considered as 0%. Thereafter, 1 mM cyanide (NaCN), an inhibitor of mitochondrial respiration, was added to allow the regeneration of the mitochondrial pool of NADH (the highest fluorescence signal was considered 100%). The baseline autofluorescence acquired from each cell was normalized between the minimal (0%) and maximal (100%) fluorescent signals to calculate the redox index.

### Mitochondrial Ca^2+^ concentration measurements with aequorin

[Ca^2+^]_m_ measurements in fibroblasts and neuronal cultures were carried out using the mitochondria-targeted luminescent aequorin probe, mtAEQ as previously described (Bonora et al., 2013). Fibroblasts were seeded at a density of 2 × 10^4^ cells/well on white 96-well plate (6005680, PerkinElmer) and cultured for 1-2 days. Similarly, iPSC-derived NPCs were seeded at a density of 5 × 10^4^ cells/well on white 96-well plate coated with poly-L-lysine and laminin for terminal neuronal differentiation and maturation using the method describe above. Two days before the experiment, cells were transduced with mtAEQ adenovirus. Following incubation, media was replaced with 5 μM coelenterazine (C2944, Invitrogen) in Krebs Ringer Buffer (125mM NaCl, 5.5mM D-Glucose, 5 mM KCl, 20mM HEPES, 1 mM Na_3_PO_4_, 1mM Glutamine, 100mM Pyruvate, and 1.2 mM CaCl_2_ at pH 7.4). The plate was then incubated in the dark for 2 h at 37 °C. Baseline luminescence signals were acquired using a plate reader (CLARIOstar, BMG Labtech) every 1 s followed by fluidic additions of either 10 μM histamine or 10 μM glutamate using integrated syringe injectors. At the end of each experiment, maximal aequorin signal was obtained by permeabilizing the cells with 1 mM digitonin and exposing the cells to a saturating Ca^2+^ concentration of 10 mM CaCl_2_. For analysis, luminescence values were converted into Ca^2+^ concentration as previously described (Bonora et al., 2013).

### Simultaneous measurement of [Ca^2+^]_c_ and [Ca^2+^]_m_ in fibroblasts

Dynamic measurements of cytosolic and mitochondrial Ca^2+^ concentrations were carried out in fibroblasts loaded with 2.5 μM Fluo-4 AM and 1 μM mt-Fura-2.3 AM (a modified version of mt-Fura-2 (Pendin et al., 2019), referred here as mito-Fura-2 AM) in RB1 containing 0.005% Pluronic F-127 for 30 minutes at 37°C. Following incubation, cells were washed thrice with RB1 and imaged using the Carl Zeiss LSM 880 confocal microscope equipped with an UV (λ_ex_ Ca^2+^ bound mito-Fura-2 = 355 nm, λ_ex_ Ca^2+^ unbound mito-Fura-2 = 405 nm, λ_em_ = 470–600 nm) and Ar laser (λ_ex_ = 488 nm, λ_em_ = 505–560 nm for Fluo-4) and quartz Plan-Apochromat 63x /1.4 oil objective at 37 °C. Images were acquired sequentially from a single z-plane every 12.5 s interval with a pixel dwell time of 1.54 μs. For all experiments, the laser illumination intensity was kept to a minimum (0.25-0.5%) to avoid phototoxicity. After acquiring baseline images for about 100 s, 10 μM histamine was applied directly into the fluorodishes using a micropipette and subsequent images were obtained. A total of 1 μM ionomycin was added at the end of each course as a positive control. Time series were analyzed in ImageJ as described above. After background subtraction, the change in Fluo-4 intensity was calculated relative to the baseline and shown as ΔF/F_0_. For mito-Fura-2, ratios between the fluorescence intensity excited at 405 nm and at 355 nm were calculated at each time point and plotted as 355/405 ratio representing [Ca^2+^]_m_. The values for area under the curve (AUC), basal mito-Fura-2 ratio, t_0.5_ to [Ca^2+^]_m_ peak and t_0.5_ of stabilization for [Ca^2+^]_c_ peak were calculated for each independent experiment in GraphPad Prism.

### Calcium retention capacity assay

The capacity of isolated mitochondria to accumulate Ca^2+^ until the mPTP opens was determined using the method described earlier (Bhosale and Duchen, 2019). Isolated mitochondria from the fibroblasts (0.5mg/ml), using the method described above, were resuspended in the RB2 (75 mM D-mannitol, 25 mM sucrose, 5 mM KH_2_PO_3_, 20 mM Tris-HCl, 100 mM KCl, and 0.1% BSA fatty acids free at pH 7.4) supplemented with 10 mM succinate and 1 μM rotenone to energize the mitochondria. 100 μl of the mitochondrial suspension was plated in a glass-bottom 96-well plate in triplicate for each condition. Extramitochondrial Ca^2+^ levels were quantified by measuring fluorescence intensity of the Ca^2+^-sensitive dye, Calcium Green-5N (C3737, Invitrogen) at 1μM concentration The fluorescence intensity was recorded using a plate reader (CLARIOstar, BMG Labtech) at 30 C° with the following filters: ex/em: 480 nm/520 nm. Ca^2+^ additions were achieved using integrated syringe injectors, where subsequent 10 μl additions of 50 μM CaCl_2_ were added for a total of 12 injections. The area under the curve was used as a measure of extramitochondrial Ca^2+^, which was expressed as a proportion of total Ca^2+^ added (Ca^2+^ free condition was used for background subtraction). This value was used to calculate the proportion of buffered Ca^2+^, and subsequent percentage inhibitions were calculated compared to untreated.

### Cytosolic and mitochondrial calcium imaging in neurons

Measurements of cytosolic Ca^2+^ concentrations were performed in neuronal cultures seeded on fluorodishes and loaded with 2.5 μM FuraFF AM (CAY20416, Cambridge Bioscience) in BrainPhys Imaging Optimized Medium containing 0.001% Pluronic F-127 for 30 minutes at 37°C. Following incubation, cells were washed thrice and imaged on a custom-made Olympus IX71 inverted epifluorescence microscope equipped with a UAPO/340 20x/0.70 water objective and a Xenon arc lamp. FuraFF was excited alternately at 340 nm ± 20 nm and 380 nm ± 20 nm and emitted light was collected through a dichroic T510lpxru (Chroma). Images were acquired with a Zyla CMOS camera (Andor) every 2 s using MetaFluor 7.8.12.0 (Molecular Devices). After acquiring baseline images for about 100 s, neurons were stimulated with 10 μM and/or 100 μM glutamate applied directly into the fluorodishes using a micropipette. A total of 2 μM ionomycin was added at the end of each experiment as positive control. For analysis, time series were imported and processed in ImageJ as described above. Ratios between the fluorescence intensity excited at 380 nm and at 340 nm were calculated at each time point and plotted as 340/380 ratio representing [Ca^2+^]_c_. Representative FuraFF ratiometric images at indicated time were obtained using Metamorph 7.8.12.0 (Molecular Devices).

Mitochondrial Ca^2+^ concentrations were measured in neurons loaded with 1 μM mito-Fura-2 AM in BrainPhys imaging medium containing 0.001% Pluronic F-127 for 30 minutes at 37°C. Following incubation, cells were washed thrice and imaged using the Carl Zeiss LSM 880 confocal microscope equipped with an UV laser and quartz Plan-Apochromat 63x/1.4 oil objective, as described above. Images were acquired from a single z-plane every 5 s interval with a pixel dwell time of 2.05 μs. After acquiring baseline images, 10 μM glutamate was applied directly into the fluorodishes using a micropipette and subsequent images were obtained. Time series were analyzed in ImageJ and the ratios between the fluorescence intensity excited at 405 nm and at 355 nm were calculated at each time point and plotted as 355/405 ratio representing [Ca^2+^]_m_. Basal mito-Fura-2 ratio, t_0.5_ to [Ca^2+^]_m_ peak and peak amplitude values were calculated for each cell in GraphPad Prism. Representative mito-Fura-2 ratiometric images were generated using Metamorph.

### Mitochondrial calcium retention assay in permeabilized neurons

To measure the capacity of neuronal mitochondria to accumulate Ca^2+^ until the mPTP opens, digitonin-permeabilized neurons were loaded with Fluo-4 AM and TMRM to simultaneously measure [Ca^2+^]_m_ uptake and the loss of ΔΨ_m_ as a readout for mPTP opening. Briefly, neuronal cultures seeded on fluorodishes, were washed once with RB1 and loaded with 5 μM Fluo-4 AM, 25 nM TMRM and 0.1% Pluronic F-127 dissolved in RB1 for 30 minutes at room temperature (RT). Following incubation, cells were permeabilized with 10 μg/ml digitonin in RB3 (5 mM NaCl, 130 mM KCl, 1 mM KH_2_PO_4_, 20 mM HEPES, 6.5 mM MgCl_2_, 1.5 mM

EGTA, 6 mM EDTA, 0.4 mM CaCl_2_, 2 mM malate, 2 mM succinate, 2 mM glutamate and 2.5 mM ADP; pH adjusted to 7.3 with 1 M KOH) containing 250 nM TMRM and 2 μM thapsigargin for 10 minutes. After permeabilization, excess digitonin was washed off and the neurons were imaged in RB3 with 250 nM TMRM and 2 μM thapsigargin to inhibit sarco/endoplasmic reticulum Ca^2+^ ATPase. Imaging was performed as described above, after baseline TMRM and Fluo-4 images, small volumes (5 or 10 μl) of 50 mM CaCl_2_ were carefully added to the chamber using a P10 micropipette until the loss of TMRM signal. Images were acquired every 120 seconds over a total time of 30 minutes. The final free Ca^2+^ ion concentration in RB3 was calculated using WEBMAXC Extended software: https://somapp.ucdmc.ucdavis.edu/pharmacology/bers/maxchelator/webmaxc/webmaxcE.htm. TMRM and Fluo-4 fluorescence intensities were calculated relative to baseline and shown as ΔF/F_0_ where ΔF is the difference in fluorescence between baseline and post CaCl_2_ addition and F_0_ is the basal fluorescence. Ratios of Fluo-4 and TMRM fluorescence intensities (ΔF/F_0_) were plotted against time and fitted with a nonlinear sigmoidal curve function.

### Live cell imaging of mtDNA release in fibroblasts

Quantitative analysis of mtDNA dynamics in fibroblasts was carried by co-labelling cells seeded in fluorodishes with 25 nM TMRM, 1 μM PicoGreen (P11495, Invitrogen), 1 μM mito-Fura-2 and 0.005% Pluronic F-127 in RB1. After incubation for 30 min at 37 °C, cells were washed thrice and imaged on Carl Zeiss LSM 880 confocal microscope equipped with an UV (λ_ex_ Ca^2+^ bound mito-Fura-2 = 355 nm, λ_ex_ Ca^2+^ unbound mito-Fura-2 = 405 nm, λ_em_ = 470– 600 nm), Ar (λ_ex_ = 488 nm, λ_em_ = 505–560 nm for PicoGreen) and DPSS laser source (λ_ex_ = 561 nm, λ_em_ = 570–700 nm for TMRM) and quartz Plan-Apochromat 63×/1.4 oil objective. Images were acquired from a single z-plane every 12.5 s interval with a pixel dwell time of 0.85 μs. After acquiring baseline images for about 250 s, 10 μM histamine was applied directly into the fluorodishes using a micropipette and subsequent images were obtained. At the end of each experiment, 1 μM FCCP was added at the end of the time course to completely depolarize mitochondria. Time series were processed and analyzed in ImageJ and Metamorph. TMRM fluorescence intensities were normalized relative to baseline and shown as ΔF/F_0_, mito-Fura-2 fluorescence intensities were plotted as 355/405 ratio representing [Ca^2+^]_m_ as described above. For the quantification of cytosolic PicoGreen puncta, number of PicoGreen puncta outside the nucleus and the TMRM-labelled mitochondrial perimeter were counted manually between the time points after the challenge with 10 μM histamine and before the application of 1 μM FCCP. Fibroblasts expressing mTagBFP-cGAS (Addgene:102603, a gift from Nicolas Manel) were co-labelled with 1 μM PicoGreen and 25 nM TMRM and imaged as described above with mTagBFP-cGAS visualized using an UV laser source (λ_ex_ = 405 nm, λ_em_ = 410–470 nm). For the quantification of the cytosolic PicoGreen puncta with cGAS, cells displaying puncta positive for cGAS and PicoGreen were selected and a binary image of TMRM and PicoGreen channels was generated in ImageJ. A segmented mask of non-mitochondrial Picogreen puncta was generated using image calculator and number of events with cGAS and PicoGreen overlap within each ROI were scored.

### Immunocytochemistry

#### Immunofluorescence analysis fibroblasts

Immunofluorescence analysis of cytosolic mtDNA puncta/ mtDNA release in fibroblasts was performed using Airyscan imaging. Briefly, cells were seeded at a density of 2 × 10^4^ cells/well in a glass-bottom 96-well plate and cultured for 1-2 days. On the day of the experiment, cells were fixed in 5% PFA in PBS for 30 min at RT, then washed three times with PBS, followed by quenching with 50 mM ammonium chloride in PBS. After fixation, cells were washed thrice with PBS and permeabilized in 0.1% Triton X-100 in PBS for 10 min, followed by three washes in PBS. Permeabilized cells were then blocked with 10% FBS in PBS, followed by incubation with primary antibodies in 5% FBS in PBS, for 2 h at RT. Cells were then washed three times with 5% FBS in PBS and labelled with the corresponding secondary antibodies prepared in 5% FBS in PBS for 1 h at RT. After three washes, super-resolution Airyscan images were acquired on a Zeiss LSM 880 with Airyscan detector in SR mode using all 32 pinholes. At least 8 z-stacks with optimal slice sizes 0.45 µm (overview image) and 0.185 µm (inset image) with a pixel dwell time of 4.10 μs were acquired. Prior to image analysis, raw confocal micrographs were automatically processed into deconvoluted Airyscan images using the Zen Black software. The number of DNA puncta outside the nucleus and mitochondrial perimeter were counted in ImageJ by generating a nuclear and mitochondrial mask and subtracting this segmented mask from DNA channel in image calculator. The colocalization coefficient (Pearson’s R-value) was quantified using Coloc 2 plugin. For 3D rendering and quantification of cytosolic mtDNA puncta or mtDNA release events, cells with enlarged nucleoids were selected and a segmented mask of IMM was created using surface function in Imaris 9.8 (Bitplane). Using this mask mtDNA nucleoids outside the surface were differentiated using spot function. A surface rendering of OMM was generated similarly and the transparency of IMM and OMM was adjusted to allow the visualization of mtDNA nucleoid spots. Details of all the antibodies used in this study can be found in Supplementary Table S3.

For the cGAS immunofluorescence, cells were cultured, fixed and permeabilized as above. After permeabilization, cells were blocked with 5% BSA in PBS for 1 h at the RT and incubated overnight with primary antibody diluted in PBS with 1% BSA and 0.1% Triton X-100. After three washes in PBS, cells were incubated with secondary antibody diluted in PBS with 1% BSA for 2 h at RT. Cells were then washed three times and incubated with DAPI for 5 min to label nuclei. After three washes, cells were imaged using the confocal microscope as described above. The nuclear cGAS intensity in each cell was quantified from the DAPI positive area and the cytosolic cGAS intensity per cell was calculated from subtracting the DAPI positive area from total cGAS mask area in ImageJ.

#### Immunofluorescence analysis neurons

Neuronal cultures were treated overnight with 10 μM glutamate and fixed with 4% PFA in 2x microtubule stabilization buffer (160 mM PIPES, 5 mM EGTA, 1 mM MgCl_2_, pH 7.2) for 10 min at RT. After fixation, cells were gently washed three times with PBS and 0.1% Triton X-100 for 10 min. Following permeabilization, cells were blocked with buffer containing 3% goat serum and 0.1% Triton X-100 in PBS for 30 minutes and incubated with primary antibodies prepared in blocking buffer for 2 h at RT. After three washes, cells were incubated with the corresponding secondary antibodies prepared in the blocking buffer for 1 h at RT. After three washes, nuclei were stained using DAPI and imaged as described above. z-stacks were imported into ImageJ and somas and axons were manually segmented using β-tubulin III staining. A colocalization mask of TOM20 and Cyt c channel was subtracted from the Cyt c channel allowing the separation of pixel intensities occurring outside the mitochondria. These binary images were used to quantify the area occupied by mitochondria without Cyt c.

### Statistical analysis

No formal statistical methods were used to predetermine sample sizes. Quantitative data are obtained from at least three independent biological replicates denoted by n_exp_, n_rep_ indicates the number of technical replicates within each independent experiment, and n_cells_ denotes the number of cells used for data analysis. Data are expressed as mean ± SD unless otherwise specified. All confocal images are representative of at least three biological replicates. Where applicable, curve fitting was performed using linear or nonlinear regression functions. Tests of normality were performed (Shapiro–Wilk test) to identify normal or non-normal populations. For normally distributed data, a paired or unpaired, two-tailed Student’s t-test was used to compare two groups. To compare more than two groups, multiple (paired or unpaired), two-tailed Student’s t-tests or a one-way or two-way analysis of variance (ANOVA) with appropriate multiple comparisons tests were used. Statistical differences were considered significant when the α-value of *p* was <0.05. Estimated *p* values are either stated as actual values or denoted by \**p* < 0.05, \*\**p* < 0.01, \*\*\**p* < 0.001, \*\*\*\**p* < 0.0001. For all experiments, the analysis was not blinded. Microsoft Excel was used to store all raw data. All statistical analyses were made using GraphPad Prism 10.4.1 (USA).

---

**Table S1.** *EPG5* mutations (NM_020964.2) in patient-derived fibroblasts.

| Patients | Variant 1 |  |  | Variant 2 |  |  | Source (Clinical presentation, outcome and Family/patient ID) |
| --- | --- | --- | --- | --- | --- | --- | --- |
|  | Nucleotide | amino acid | Exon | Nucleotide | amino acid | Exon |  |
| Patient 1 (Pat1) | c.1007A>G | p.Q336R | 2 | c.1007A>G | p.Q336R | 2 | Cullup et al., 2013, Patient ID- 7.1 |
| Patient 2 (Pat2) | c.4862G>A | p.R1621G | 28 | c.4862G>A | p.R1621G | 28 | Vansenne et al., 2022, Patient ID- 1.1 |
| Patient 3 (Pat3) | c.895C>T | p. R299* | 2 | c.5479C>G | p.P1827A | 31 | Dafsari et al., 2024, Patient ID- |
| Patient 4 (Pat4) | c.4952+1G>A | p.F1604Gfs*20 | 28 | c.4952+1G>A | p.F1604Gfs*20 | 28 | Byrne et al., 2016, Patient ID- 4.1 |

**Table S2.**
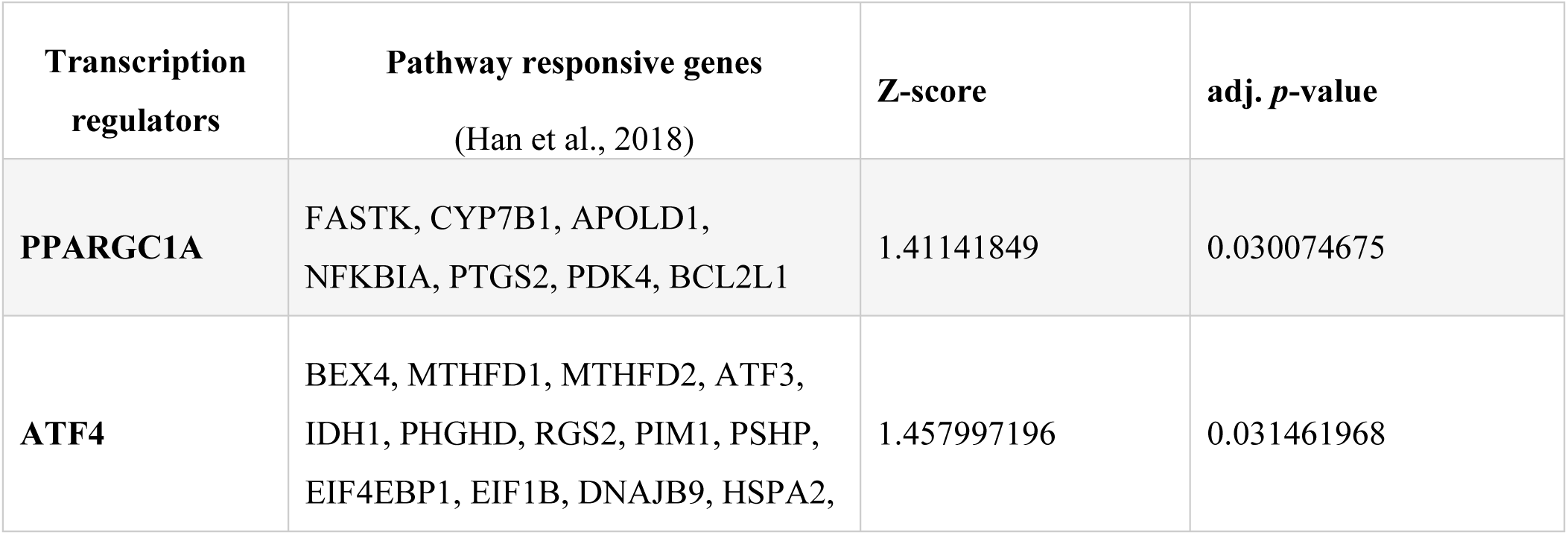
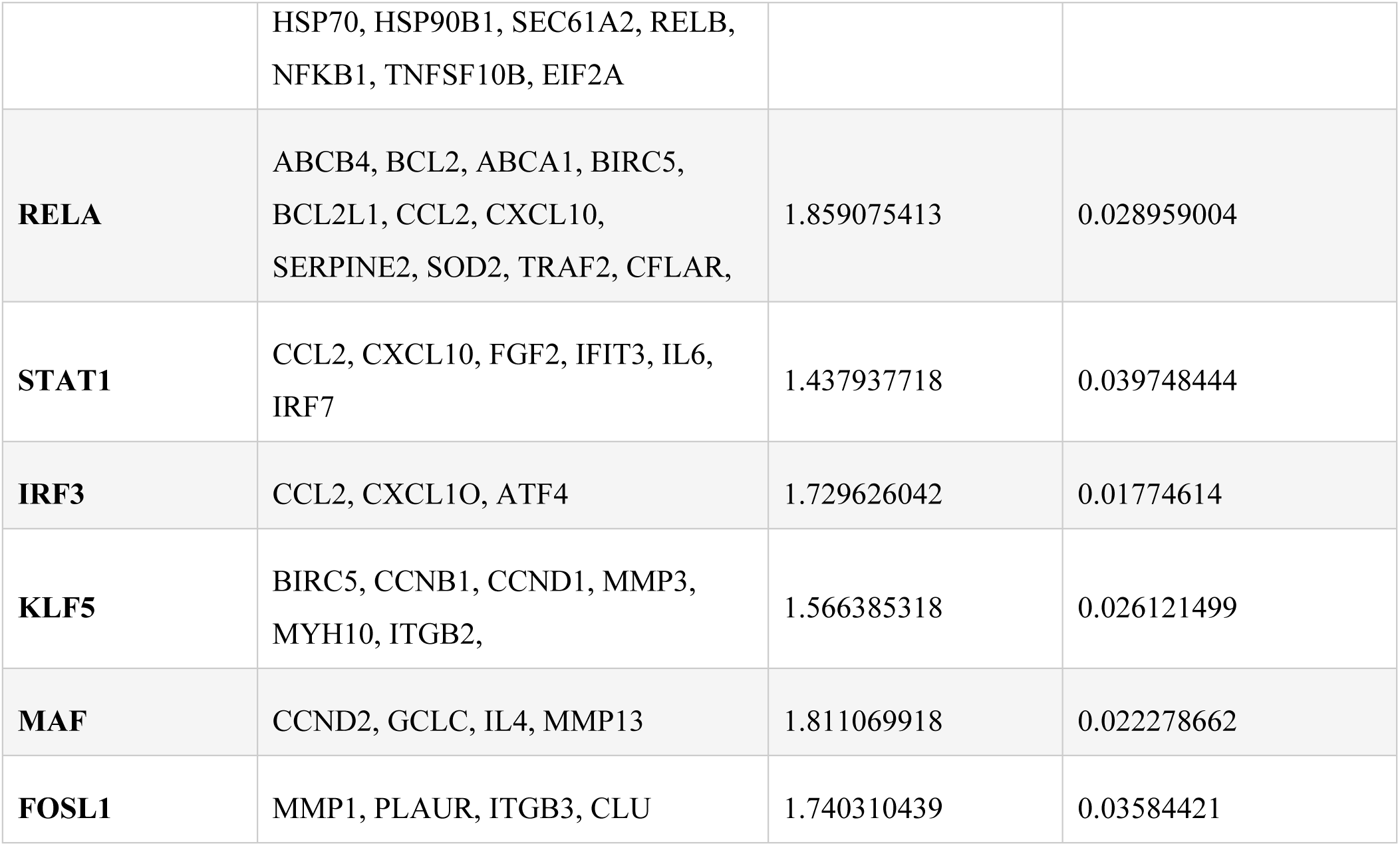
Upstream analysis of transcription regulators in patient fibroblasts.

| Transcription regulators | Pathway responsive genes<br>(Han et al., 2018) | Z-score | adj. *p*-value |
| --- | --- | --- | --- |
| **PPARGC1A** | FASTK, CYP7B1, APOLD1, NFKBIA, PTGS2, PDK4, BCL2L1 | 1.41141849 | 0.030074675 |
| **ATF4** | BEX4, MTHFD1, MTHFD2, ATF3, IDH1, PHGHD, RGS2, PIM1, PSHP, EIF4EBP1, EIF1B, DNAJB9, HSPA2, | 1.457997196 | 0.031461968 |
|  | HSP70, HSP90B1, SEC61A2, RELB, NFKB1, TNFSF10B, EIF2A |  |  |
| **RELA** | ABCB4, BCL2, ABCA1, BIRC5, BCL2L1, CCL2, CXCL10, SERPINE2, SOD2, TRAF2, CFLAR, | 1.859075413 | 0.028959004 |
| **STAT1** | CCL2, CXCL10, FGF2, IFIT3, IL6, IRF7 | 1.437937718 | 0.039748444 |
| **IRF3** | CCL2, CXCL10, ATF4 | 1.729626042 | 0.01774614 |
| **KLF5** | BIRC5, CCNB1, CCND1, MMP3, MYH10, ITGB2, | 1.566385318 | 0.026121499 |
| **MAF** | CCND2, GCLC, IL4, MMP13 | 1.811069918 | 0.022278662 |
| **FOSL1** | MMP1, PLAUR, ITGB3, CLU | 1.740310439 | 0.03584421 |

**Table S3:** Key resources table.

| Reagent or resource | Source | Identifier |
| --- | --- | --- |
| **Antibodies** |  |  |
| Rabbit pAb anti-EPG5 (1:1000) | Abcam | Cat# ab122186 |
| Mouse mAb anti-β-actin (1:10000) | Cell Signaling Technology | Cat# 3700 |
| Mouse Ab anti-OxPhos cocktail (1:1000) | Invitrogen | Cat# 45-8199 |
| Rabbit mAb anti-TOM20 (1:5000 IB, 1:200 IF) | Abcam | Cat# ab186735 |
| Mouse mAb anti-MCU (1:1000) | Sigma-Aldrich | Cat# AMAB91189 |
| Rabbit pAb anti-MCUB (1:1000) | Proteintech | Cat# 20387-1-AP |
| Rabbit pAb anti-MICU1 (1:1000) | Thermo Scientific | Cat# HPA037480 |
| Rabbit pAb anti-MICU2 (1:1000) | Sigma-Aldrich | Cat# HPA045511 |
| Rabbit pAb anti-MICU3 (1:1000) | Thermo Scientific | Cat# PA5-55177 |
| Rabbit pAb anti-EMRE (1:1000) | Abcam | Cat# ab157387 |
| Rabbit pAb anti-NCLX (1:1000) | Sigma-Aldrich | Cat# SAB2102181 |
| Mouse mAb anti-ATP5A (1:1000) | Abcam | Cat# ab14748 |
| Rabbit pAb anti-phospho PDH E1α (Ser293) (1:1000) | Sigma-Aldrich | Cat# AP1062 |
| Mouse mAb anti- PDH E1α (1:5000) | Abcam | Cat# ab110330 |
| Rabbit pAb anti-STAT1 (1:1000) | Cell Signaling Technology | Cat# 9172S |
| Rabbit mAb anti-phospho STAT1 (Tyr701) (1:1000) | Cell Signaling Technology | Cat# 9167S |
| Rabbit mAb anti-TBK1 (1:1000) | Cell Signaling Technology | Cat# 3504S |
| Rabbit mAb anti-phospho TBK1 (Ser172) (1:1000) | Cell Signaling Technology | Cat# 5483S |
| Rabbit mAb anti-STING (1:1000) | Cell Signaling Technology | Cat# 13647S |
| Rabbit pAb anti-IRF3 (1:1000) | Abcam | Cat# ab25950 |
| Rabbit mAb anti-phospho IRF3 (Ser396) (1:1000) | Cell Signaling Technology | Cat# 29047S |
| HRP-Goat Anti-Rabbit IgG (H+L) | Jackson ImmunoResearch | Cat# 111-035-045 |
| HRP-Goat Anti-Mouse IgG (H+L) | Jackson ImmunoResearch | Cat# 315-035-045 |
| Rabbit mAb anti-cGAS (1:100 IF) | Cell Signaling Technology | Cat# 15102S |
| Rabbit pAb anti-Citrate synthetase (1:100 IF) | Abcam | Cat# ab96600 |
| Mouse mAb anti-DNA (1:200 IF) | Sigma-Aldrich | Cat# CBL186 |
| Mouse mAb anti-Cytochrome C (1:200 IF) | BD Pharmingen | Cat# 556432 |
| Mouse mAb anti-Beta-Tubulin III (TUJ1) (1:100 IF) | STEMCELL Technologies | Cat# 60052 |
| Rabbit pAb anti-GFAP (1:100 IF) | Sigma-Aldrich | Cat# AB5804 |
| Chicken pAb anti-MAP2 (1:100 IF) | Abcam | Cat# ab92434 |
| Alexa Fluor 647 Donkey anti-Rabbit IgG (H+L) | Thermo Scientific | Cat# A-31573 |
| Alexa Fluor 488 Donkey anti-Mouse IgG (H+L) | Thermo Scientific | Cat# A-21202 |
| Alexa Fluor 568 Donkey anti-Rabbit IgG (H+L) | Thermo Scientific | Cat# A-10042 |
| Alexa Fluor 488 Goat anti-Rabbit IgG (H+L) | Thermo Scientific | Cat# A-11008 |
| Alexa Fluor 568 Goat anti-Mouse IgG (H+L) | Thermo Scientific | Cat# A-11004 |
| Alexa Fluor 647 Goat anti-Chicken IgG (H+L) | Thermo Scientific | Cat# A-21449 |
| **Commercial Kit Assays** |  |  |
| MycoAlert Mycoplasma Detection Kit | Lonza | Cat# LT07-118 |
| Seahorse XF Cell Mito Stress Test Kit | Agilent | Cat# 103015-100 |
| Seahorse XFe96/XF Pro FluxPak | Agilent | Cat# 103793-100 |
| Human Dermal Fibroblasts Nucleofector Kit | Lonza | Cat# VPI-1002 |
| NativePAGE Novex Bis-Tris Gel System | Thermo Scientific | Cat# BN2007, BN2008 |
| **Experimental Models: Cell Lines** |  |  |
| Control Fibroblasts | This paper | PMID: 24336167 and<br>MRC CNMD Biobank<br>London |
| Patient Fibroblasts | This paper | Table S1 |
| Isogenic iPSCs | Coriell Institute | RRID: GM28930 |
| Q336R iPSCs | Coriell Institute | RRID: GM27291 and<br>PMID: 35700637 |
| **Critical chemicals and reagents** |  |  |
| JP1-138 | Valeria Pingitore et al., 2024 | PMID: 38985859 |
| Mito-Fura-2 AM (v2.3) | Pendin et al., 2019 | PMID: 31132197 |
| **Plasmid and AAV vector** |  |  |
| pTRIP-CMV-mTagBFP2-2A-FLAG-ntcGAS | Addgene, (Gentili et al., 2015) | Cat# 102603 |
| Ade-mtaequorin | Gift from Rosario Rizzuto lab, (Tosatto et al., 2016) | PMID: 27138568 |
| **Software and algorithms** |  |  |
| FIJI (ImageJ); Version 1.54p | http://fiji.sc, (Schindelin et al., 2012) | RRID: SCR_002285 |
| Prism 10 | GraphPad | RRID: SCR_002798 |
| MitoSegNet | https://github.com/MitoSegNet, (Fischer et al., 2020) | PMID: 33083756 |
| Image Lab software; Version 6.1 | Bio-Rad | RRID: SCR_014210 |
| Microscopy Image Browser; Version 2.81 | https://mib.helsinki.fi/, (Belevich and Jokitalo, 2021) | RRID:SCR_016560 |
| Imaris Version 9.8 | Bitplane | RRID:SCR_007370 |
| MetaMorph Microscopy Automation and Image Analysis Software | Molecular Devices | RRID:SCR_002368 |
| MetaFluor Fluorescence Ratio Imaging Software | Molecular Devices | RRID:SCR_014294 |
| ZEISS ZEN Microscopy Software | Carl Zeiss | RRID:SCR_013672 |
| Seahorse Wave | Agilent | RRID:SCR_014526 |
| R Project | http://www.r-project.org/ | RRID:SCR_001905 |

**Table S4.**
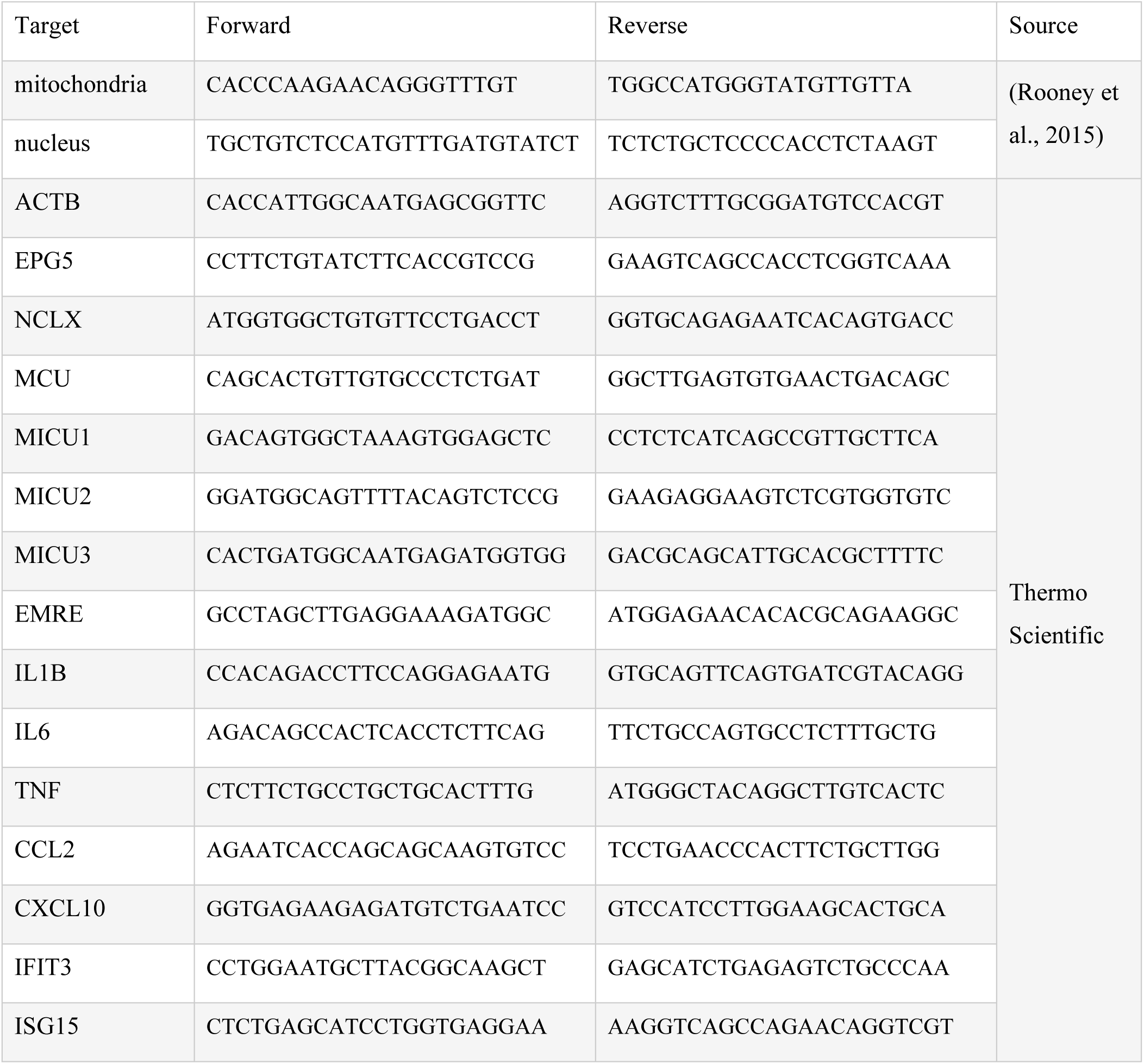
Primer pairs used in this study.

## Resource availability

### Lead Contact

Further information and requests for resources and reagents should be directed to and will be fulfilled by the Lead Contact, Michael R Duchen.

### Materials Availability

This study did not generate new unique reagents.

### Data and Code Availability

Raw FastQ files for RNA-sequencing analyses will be deposited to GEO upon publication. Gene-set Analyses using biological processes and the KEGG pathway analysis of the RNA-sequencing experiment as well as the unprocessed blot images deposited at Mendeley Data upon publication. This paper does not report original code.

## Acknowledgements

We would like to thank the Coriell Institute for Medical Research (Camden, NJ, USA) for their kind gift of patient fibroblasts with EPG5 mutations. We are thankful for the valuable discussions and support from all the members of M. Duchen, G. Szabadkai and M. Fanto labs. We acknowledge The MRC Centre for Neuromuscular Diseases Biobank (supported by the National Institute for Health Research Biomedical Research Centres at Great Ormond Street Hospital for Children, NHS Foundation Trust) for providing all age and sex-matched healthy control fibroblasts used in this study. We also acknowledge UCL Genomics Core Facility for the RNA-sequencing studies. This work was supported by grants from Action Medical Research (GN2959 to K.S. and M.R.D.) and Great Ormond Street Hospital for Children (V4218 to M.R.D.).

## Author contributions

**K.S.** and **M.R.D.**- Conceptualization, funding acquisition, investigation, visualization, methodology, writing–original draft, project administration, writing–review and editing. **H.J.**,

**H.S.D.** and **M.F.**- Conceptualization, resources, investigation, formal analysis, supervision, writing–review and editing. **O.G.** and **H.C.** - Formal analysis, methodology, investigation and data curation. **I.M., I.K.**, **P.S.** and **C.Y.C.**- Formal analysis, methodology and investigation, writing-review and editing. **V.P.**, **D.L.S., F.V.** and **D.P.**- Resources, writing-review and editing. **G.S.**- Conceptualization, writing-review and editing.

